# De Novo Design of C–H Hydroxylation Enzymes with Programmed Selectivities

**DOI:** 10.64898/2026.11.12.766315

**Authors:** Indrek Kalvet, Jihun Jeung, Shilong Gao, Alessia C. Andrews, Belinda B. Wenke, Ross Smithson, Johnny Mendoza, Hannah Nguyen, Emily Joyce, Alex Kang, Asim K. Bera, Anthony P. Green, David Baker

## Abstract

The ability to precisely functionalize individual C–H bonds in complex organic molecules would transform synthetic strategies to produce essential chemicals. In nature, C–H oxidations are commonly achieved using metalloenzymes such as cytochrome P450s, however adapting these systems to achieve selective conversions of complex substrates requires extensive enzyme discovery and engineering campaigns. Here we show that selective C–H oxidation biocatalysts can be programmed computationally from minimal structural inputs comprising a Cys-ligated heme cofactor, catalytic side chains required for H_2_O_2_ activation and heterolysis, and the target substrate precisely positioned adjacent to the resulting ferryl intermediate (Compound I). We designed heme hydroxylases for increasingly complex substrates, progressing from aromatic and aliphatic oxidations of fluorogenic model compounds to regiodivergent oxidation of estradiol at either the benzylic C6 (93% d.e.) or unactivated C16 (99% d.e.) positions. Crystal structures across all campaigns confirm the accuracy of the design pipeline. Together, these results establish a general framework for constructing selective heme hydroxylases around predefined substrates and reaction geometries, providing a path toward custom biocatalysts for site-selective C–H functionalizations.

## Introduction

Hydroxylation of carbon-hydrogen bonds plays a key role in biology and chemical synthesis (*1*). Installation of a hydroxyl group alters molecular properties, including solubility and interactions with biological targets, while also providing a synthetic handle for downstream functionalizations (Figure 1A). As drug discovery and chemical synthesis increasingly demand access to structurally complex molecules, there is a growing need for hydroxylation methods capable of targeting specific C–H bonds with high chemo-, regio-, and stereoselectivity (*2*). Synthetic transition metal catalysts enable C–H oxidation but are often constrained by the intrinsic reactivity of individual bonds (*3–5*). A structurally diverse array of enzymes has evolved to achieve remarkable selectivity in C–H hydroxylation (*1, 6, 7*). The most prominent of these are heme-dependent hydroxylases, which proceed through a highly reactive iron-oxo intermediate, Compound I (CpdI), capable of oxidizing even unactivated C–H bonds (Figure 1B). Cytochrome P450 monooxygenases access the CpdI intermediate through activation of molecular oxygen, but doing so requires delivery of reducing equivalents and coordinated electron-transfer machinery, adding substantial catalytic complexity (*8, 9*). In contrast, peroxygenases use H_2_O_2_ to access the same reactive iron-oxo intermediate directly (Figure 1C), bypassing the need for external electron-transfer partners and providing a simpler catalytic framework for design and engineering (*10, 11*). The operational simplicity of peroxide-driven catalysis has driven considerable interest in peroxygenase enzymes, with controlled peroxide delivery and enzyme engineering providing routes to mitigate peroxide sensitivity limitations (*12*). Beyond natural enzymes, peroxygenases have been created by repurposing P450 monooxygenases for peroxide-driven catalysis (*13–15*), by embedding synthetic cofactors to create new artificial metalloenzymes (*16, 17*), and their scope has been expanded using computational design (*18–20*). However, these approaches require adapting pre-existing catalytic frameworks, limiting the extent to which activity and selectivity can be specified a priori for a new target transformation. Retargeting natural scaffolds to new substrates or oxidation sites can also require extensive discovery and engineering campaigns (*21, 22*). Deep-learning-based atomistic design provides a promising platform for the creation of efficient and selective biocatalysts for synthetically valuable transformations (*23*), and incorporation of cofactors (*24–28*) has enabled design of metalloporphyrin-based catalysts for non-natural reactions such as carbene and nitrene transfer (*29–34*) and CO_2_ reduction (*35*). However, de novo design of hydroxylases tailored to a specific site on an arbitrary target molecule, which would provide a powerful complement to existing synthetic and biocatalytic approaches, remains an outstanding challenge.

**Figure 1.**
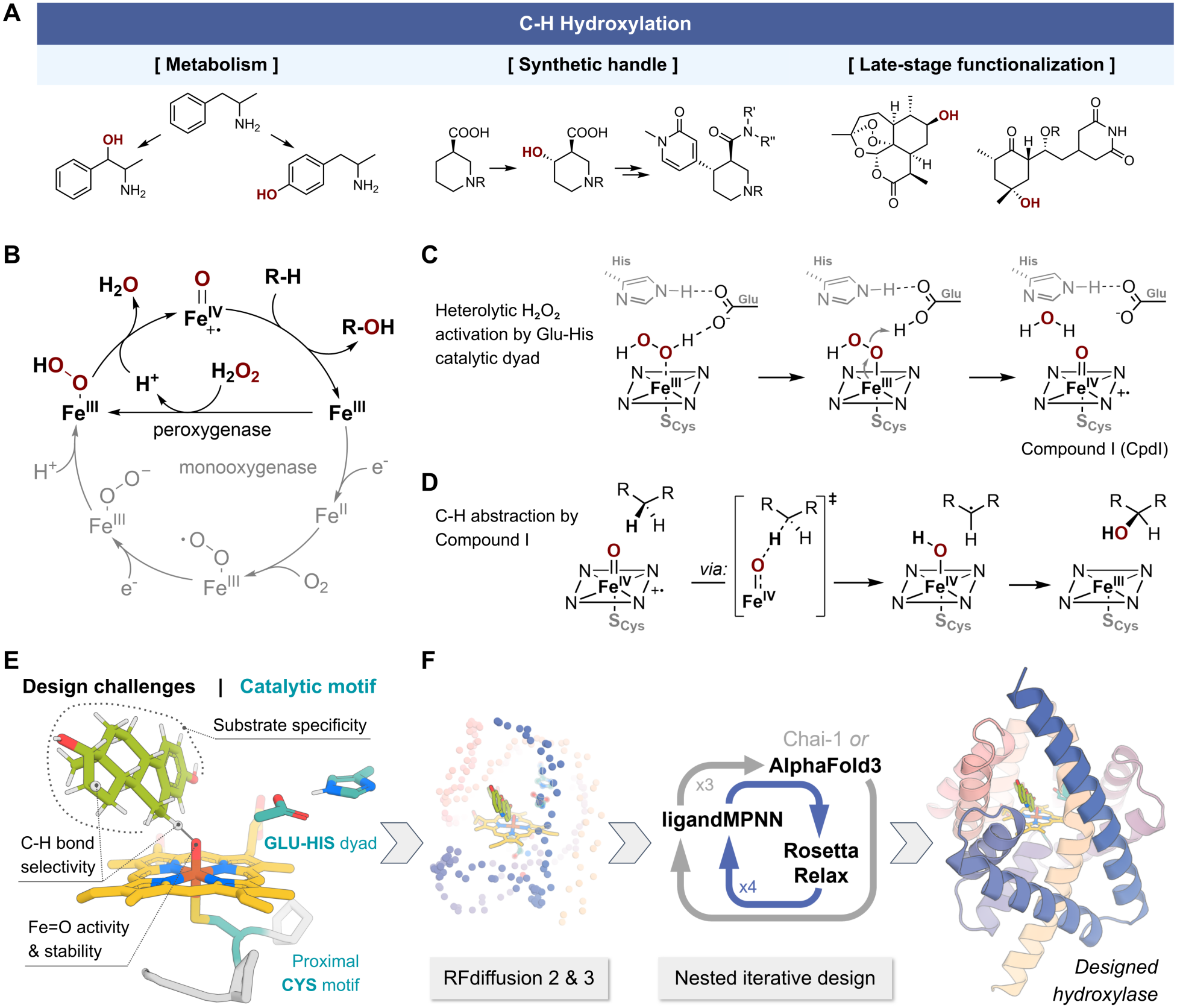
**(A)** Importance of C–H hydroxylation reactions in metabolism, as a synthetic handle (*36*), and in late-stage functionalization of drug molecules (*2*). **(B)** Mechanisms of C–H hydroxylation by activation of O_2_ or H_2_O_2_. **(C)** Glu-His catalytic dyad in peroxygenases activates H_2_O_2_ by shuffling protons and promoting heterolytic O–O cleavage to form iron-oxo Compound I (*37, 38*). **(D)** Mechanism of C–H hydroxylation by Compound I mediated C–H abstraction and radical rebound. **(E)** Challenges of biocatalytic C–H hydroxylation: precise and specific substrate binding, balancing the activity and stability of iron-oxo. Design input consists of a glutamate-histidine catalytic dyad and a cysteine PCPXX motif taken from native *Hsp*UPO, placed next to the transition state of C–H abstraction by iron-oxo. **(F)** De novo peroxygenase design workflow: backbones are generated by RFD2 and RFD3, followed by iterative sequence design, with final designs selected by stringent filtering using an all-atom structure prediction model (AlphaFold3/Chai-1).

We set out to overcome this challenge by developing a heme peroxygenase based framework for building selective C–H hydroxylases beyond the constraints of natural enzyme evolution. We reasoned that site selective hydroxylation could be achieved by de novo design of catalytic sites which organize the heme for peroxide activation while precisely positioning the substrate relative to the reactive iron-oxo intermediate in protein architectures custom built for the reaction of interest (Figure 1E). This is challenging on multiple levels because the designed enzyme must reliably bind the heme cofactor, accurately position catalytic residues that facilitate iron-oxo formation, and ensure efficient substrate access and precise positioning adjacent to CpdI while avoiding unwanted oxidative side-reactions. If successfully implemented, this strategy could provide a general route to creating selective hydroxylases on demand, expanding access to site-specific C–H functionalization across diverse molecular scaffolds and synthetic targets.

### Principles of De novo Hydroxylase Design

Our design strategy positions a minimal heme-based hydroxylation machinery adjacent to the target C–H bond of the substrate. For aliphatic C–H hydroxylation, the rate-limiting and selectivity-determining step is the abstraction of a hydrogen atom from a substrate by CpdI, forming a transient in-cage carbon radical that rapidly converts to product through hydroxyl rebound (Figure 1D) (*1*). We constructed idealized active-site models (theozymes) using DFT-derived transition-state geometries for C_sp3_–H abstraction. For aromatic hydroxylation, we instead used a transition state where the target ring carbon is positioned for electrophilic attack by CpdI.

We selected cysteine as the axial heme ligand because its strongly electron-donating thiolate enhances CpdI reactivity toward unactivated C–H bonds (*9*). To promote CpdI formation, we incorporated a Glu–His dyad (Figure 1C) in which the His-stabilized Glu facilitates proton transfer between the peroxide oxygens to facilitate heterolytic O–O cleavage and water release, but did not explicitly model this activation pathway. Coordinates for both catalytic elements were taken from native *Hsp*UPO (*39*) and positioned around the computed CpdI-substrate transition state. The Cys-containing PCPXX loop–helix motif was fixed in backbone and side-chain coordinates to preserve stabilization by neighboring prolines and a Cys sulfur–backbone NH hydrogen bond. For the Glu–His dyad, only the Glu carboxylate and C_γ_ atoms and the His imidazole ring were fixed, the minimal input necessary for catalytic tip-atom scaffolding (Figure 1E, see SI for theozyme modeling details).

We used RFdiffusion2 (*40*) and RFdiffusion3 (*41*) (RFD2 and RFD3) to build new protein structures around this minimal theozyme geometry in an unindexed fashion, allowing the model to dynamically solve the primary sequence positions and rotamers of the catalytic residues. Sequence refinement was carried out through iterative application (*42*) of LigandMPNN (*43*), Rosetta packing and minimization, and all-atom structure models including AlphaFold3 (AF3) (*44*) and Chai-1 (*45*) (Figure 1F, see SI for detailed description). Designs for which their structure predictions were in close agreement with the design models over the overall fold, catalytic residues, heme and substrate (low RMSD and high pLDDT/ipTM across five samples), and which had favorable hydrogen bonds and physics-based metrics describing the heme and substrate binding interfaces (Rosetta ddG, contact molecular surface, SASA) were subjected to a final preorganization analysis using PLACER (*46*). The high reactivity of the iron-oxo requires the substrate to be readily accessible to prevent unwanted side-reactions, and bound precisely to selectively react with the target C–H bond; we selected designs that maintain consistent placement of catalytic residues and the substrate in the PLACER ensembles for experimental characterization.

In the following sections, we first evaluate this design strategy towards aromatic and aliphatic C–H hydroxylations of model fluorogenic substrates, and then proceed to functionalize individual C-H bonds in more complex, and synthetically valuable molecules.

### Aromatic hydroxylase design

We targeted aromatic C–H hydroxylation using the fluorogenic 1,8-naphthalimide model substrates **F-8** and **NEN**, enabling sensitive activity detection at λ_ex_/λ_em_ = 450/558 nm (Figure 2A) (*47, 48*). Because the substrates share the same reactive geometry, we reasoned that a pocket designed around the larger **F-8** could also accommodate **NEN** in a productive pose (Figure 2B). Using a RFD2-MPNN-FastRelax pipeline and Chai-1 filtering (see SI section 2.1), we selected 90 designs spanning 29 unique folds for experimental characterization (Figure S3). Fluorescence screening of hemin-loaded, Strep-tag-purified proteins identified one design, **dOHar-1**, with modest but detectable activity toward **NEN**, confirmed by LC-MS, but no detectable activity toward **F-8** (Figure S4). AF3 models provided a structural rationale for the observed reactivity differences, placing **NEN** near the intended transition-state pose and **F-8** in a flipped orientation (Figure S5).

**Figure 2.**
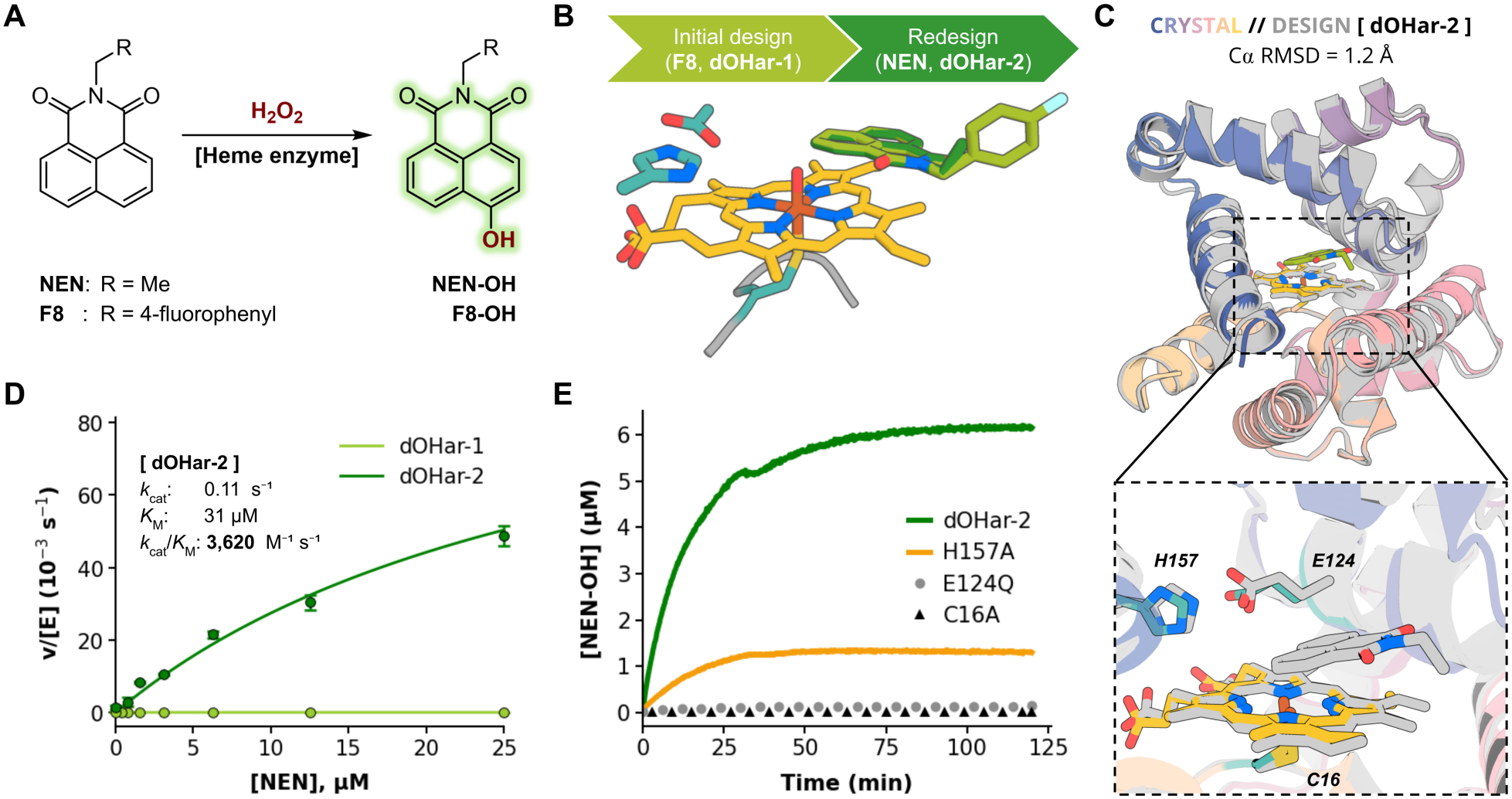
De novo design of aromatic hydroxylase. **(A)** The model aromatic hydroxylation reaction of 1,8-naphthalimide derivatives. **(B)** RFdiffusion2 input of a computed theozyme model of **F-8** (light green) and sequence redesign input of **NEN** (dark green) hydroxylation **(C)** Crystal structure of **dOHar-2**, bottom inset showing heme and catalytic dyad overlay. **(D)** Michaelis-Menten kinetics analysis of designs **dOHar-1** and **dOHar-2** ([E] = 10 nM, [H_2_O_2_] = 4 mM). **(E)** Knockout mutagenesis of catalytic residues of **dOHar-2**.

Because **dOHar-1** was initially designed around **F-8**, we sought to improve its activity by redesigning its sequence for the smaller **NEN** substrate. Using iterative enhancedMPNN (*49*) and Rosetta Relax (*42*), we generated 40 sequences on the **dOHar-1** backbone. Eleven designs showed higher **NEN**-hydroxylation activity than **dOHar-1**, with the most active variant, **dOHar-2**, exhibiting a 29-fold increase (Figure S6). Kinetic analysis of **dOHar-2** revealed a catalytic efficiency (*k*_cat_/*K*_M_) of 3,620 ± 762 M^-1^s^-1^ towards **NEN** (Figure 4D). The crystal structure of **dOHar-2**, determined at 2.03 Å resolution, closely matched the design model (C_α_ RMSD of 1.2 Å), including the placement of catalytic residues and the heme cofactor (Figure 4C). The Cys16-ligated heme formed hydrogen bonds with Arg121, the backbone amide NH of His15, and a water molecule (Figure S12). Mutagenesis supported the proposed roles of the catalytic residues, with C16A and E124Q abolishing the activity and H157A substantially reducing it (Figure 4C). The enzyme mechanism was further interrogated by performing reactions with isotopically labeled 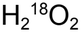, which confirmed that the oxygen atom introduced into the **NEN-OH** product is derived from hydrogen peroxide. (Figure S11). The closest structural match to **dOHar-2** in the PDB had a TM-score of 0.49 (Figure S13) (*50*).

### Aliphatic hydroxylase design

We next sought to extend our heme hydroxylase design strategy to aliphatic C–H hydroxylations using benzyloxyresorufin (**BR**), a fluorogenic substrate commonly used to assay P450 activity through *O*-dealkylation (BROD reaction, Figure 3A) (*51*). Hydroxylation of the benzylic position generates an unstable hemiacetal that fragments into fluorescent resorufin and benzaldehyde. We used our RFD2- and RFD3-based de novo design strategy with AF3 filtering to build enzymes placing the benzylic C–H bond of **BR** substrate next to the iron-oxo (see SI section 3.1.1 for details), selecting 176 designs originating from 33 unique protein folds for experimental testing (Figure S15-16). Designs were expressed in *E. coli*, and assayed as clarified lysates supplemented with 10 μM hemin (see SI section 3.2.1 for detailed description). We identified designs **F9** and **H5** (hereafter **dOHbr-1** and **dOHbr-2**) with notably increased fluorescence over background (Figure 3B). Liquid chromatography analysis of the enzymatic products verified the formation of resorufin and benzaldehyde (Figure S23).

**Figure 3.**
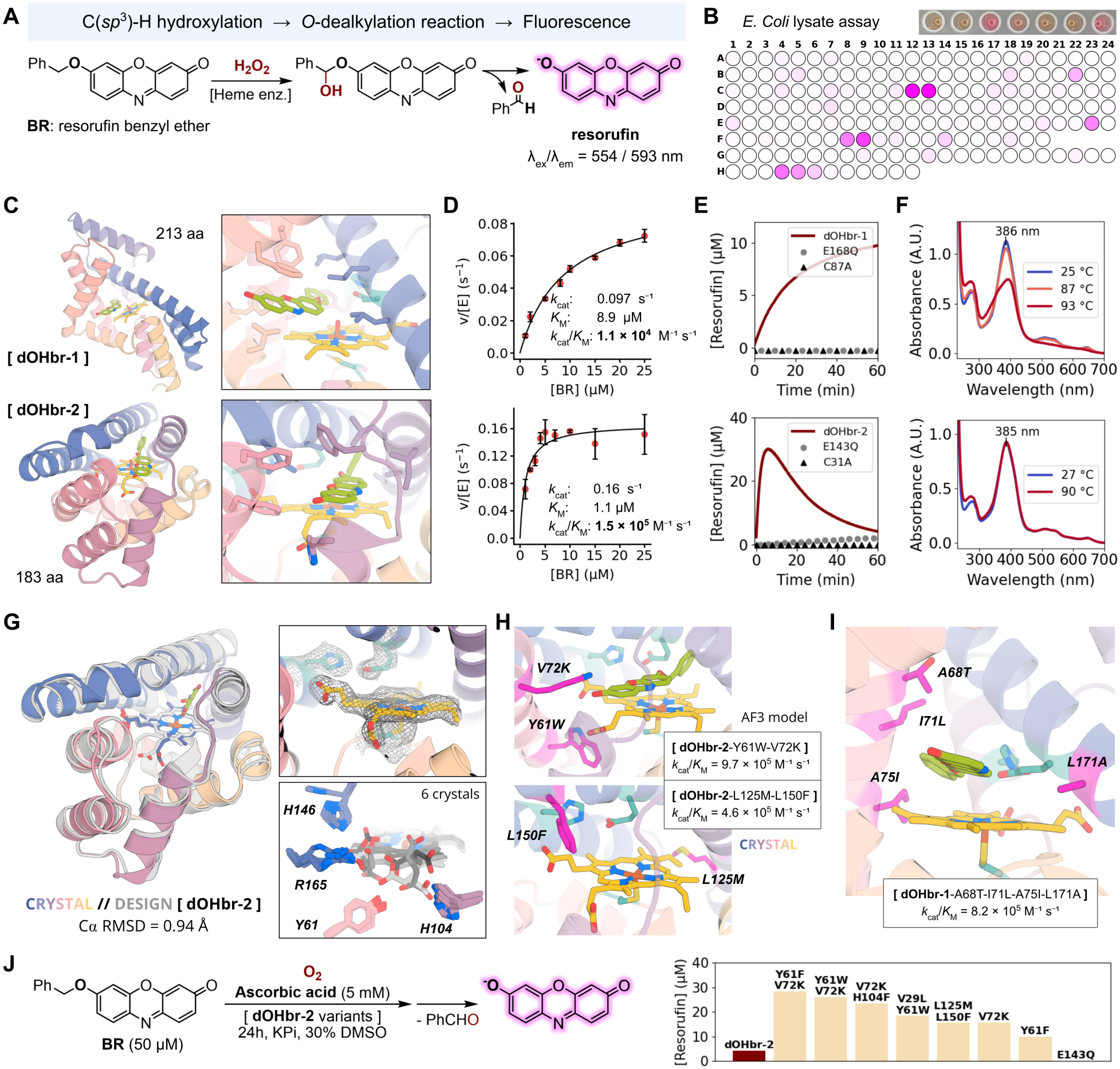
Resorufin ether C–H-hydroxylase design and characterization. **(A)** Resorufin benzyl ether *O*-dealkylation model hydroxylation reaction. **(B)** Enzymatic activity screening in clarified *E. coli* cell lysate, with visible color from the release of resorufin. The platemap is colored according to normalized activities. **(C)** Structural models of best-performing designs **dOHbr-1** and **dOHbr-2. (D)** Michaelis-Menten kinetic analysis of the designs. **dOHbr-1** was measured at 500 nM and **dOHbr-2** at 50 nM [enzyme], 4 mM H_2_O_2_. **(E)** Mutating the designed catalytic Cys and Glu residues abolishes hydroxylase activity (measured with 2 μM [enzyme], 50 μM [BR], 4 mM H_2_O_2_). Bleaching of resorufin is observed with **dOHbr-2. (F)** UV/Vis spectra collected at increasing temperatures. **(G)** Crystal structure of **dOHbr-2.**Top inset shows the density (2*F*o–*F*c map contoured at 1σ) of heme and the catalytic motif. Bottom inset shows the conformational flexibility of heme propionates across 6 crystal structures. **(H)** Top variants from the mutagenesis of **dOHbr-2**: (top) AF3 prediction of Y61W-V72K; (bottom) crystal structure of L125M-L150F. **(I)** AF3 predicted a model of the highly efficient **dOHbr-1-TLIA** variant. **(J)** Hydroxylation of resorufin benzyl ether, catalyzed by **dOHbr-2** variants (500 nM), using O_2_ and 5 mM ascorbic acid instead of H_2_O_2_.

Kinetic characterization (Figure 3D) of **dOHbr-1** and **dOHbr-2** reveals catalytic efficiencies (*k*_cat_/*K*_M_) of (1.1 ± 0.1) × 10^4^ M^-1^s^-1^ (*k*_cat_ = 0.097 ± 0.004 s^-1^, *K*_M_ = 8.9 ± 0.8 μM) and (1.6 ± 0.3) × 10^5^ M^-1^s^-1^(*k*_cat_ = 0.18 ± 0.01 s^-1^, *K*_M_ = 1.2 ± 0.2 μM), respectively. Substituting the designed axial Cys with Ala or the catalytic Glu with Gln in either enzyme abolishes hydroxylation activity (Figure 3E), supporting the essential roles of these residues in catalysis. The UV/Vis spectra of **dOHbr-1** and **dOHbr-2** are consistent with a high spin pentacoordinate Cys-ligated heme (λ_Soret_ ∼385 nm, Figure 3F) (*52*) while UV/Vis and circular dichroism (CD) spectra show that both designs are thermostable, retaining heme-binding and protein secondary structure up to 87 °C (Figures S31-32).

Design **dOHbr-1** is a RFD2-generated 213-residue protein with an open heme binding site and a semi-exposed substrate interface (Figure 3C). **dOHbr-2** has a 183-residue scaffold with a relatively deep substrate binding site, generated with RFD3 (Figure 3C). Consistent with these pocket geometries, the *K*_M_ of **dOHbr-2** is 8-fold lower than that of **dOHbr-1**, indicating tighter substrate binding, while the two designs have comparable *k*_cat_. The crystal structure of **dOHbr-2**, determined at 2.0 Å resolution, closely matches the design model with C_α_ RMSD of 0.9 Å (Figure 3G). The crystal structure confirmed the placement of the Cys31-ligated heme, with heme hydrogen-bonding contacts involving Tyr61, His104, Arg165, and His146 (Figure 3G). Catalytic side-chain placements were largely recovered, and the substrate-binding pocket is predominantly hydrophobic on the distal side of the heme. Additional structures of **dOHbr-2** showed consistent porphyrin-ring positioning but some flexibility in the heme propionates (Figure 3G, bottom inset; Figure S35).

Additional hits included **H4** (**dOHbr-3**), a sequence variant of **dOHbr-2** differing at 25 positions (Figure S33), and closely related **C12** and **C13** (**dOHbr-4** and **-5**; differing at 15 positions; Figure S34); these designs are further characterized in the SI. A 2.2 Å resolution crystal structure of RFD2-designed **dOHbr-5** matches the design model closely, both in the protein backbone (C_α_ RMSD of 0.45 Å) and in the placement of heme and catalytic residues (Figure S37). These designs are structurally distinct from known proteins on the PDB, as indicated by the TM scores of 0.53 (**dOHbr-1**), 0.50 (**dOHbr-2**), and 0.57 (**dOHbr-5**) (Figure S38-40).

Next, we sought to systematically evaluate the sequence-dependence of the hydroxylase activity. We constructed a library of 382 variants in 42 positions surrounding the active site of **dOHbr-2** predicted by AF3 to retain the designed structure (Figure S17-18) (*53*). Mutation-bearing fragments encoding one-third of the sequence were synthesized as an oligonucleotide pool, separated into individual wells by two PCR reactions, assembled into plasmids, and expressed in *E. coli* (see SI section 3.2.3 for details, Figure S24). Consistent with the flexibility observed in the crystal structure, many of the activity increasing mutations were clustered around the heme carboxylate groups, at L23, V29, Y61, F69, H104, V72 and L150, potentially stabilizing their placement through either direct hydrogen bonds or sterically occluding some propionate conformers (Figure S25). There were only a few beneficial mutations adjacent to the substrate, suggesting that computational design yielded a sufficiently optimal protein-substrate interface. Combining subsets of the beneficial mutations (Figure S28; see SI section 3.2.3 for details) led to considerable increases in *k*_cat_ over the parent **dOHbr-2**, with overall catalytic efficiencies (*k*_cat_/*K*_M_) becoming comparable to native hydroxylases (>10^5^ M^-1^s^-1^) (*39*). The most active **dOHbr-2**-Y61W-V72K reached a *k*_cat_/*K*_M_ of (9.7 ± 2.0) × 10^5^ M^-1^ s^-1^ (*k*_cat_ = 1.6 ± 0.1 s^-1^, *K*_M_ = 1.7 ± 0.3 μM; Figure 3H, top inset). Five of the top variants carried Y61F, Y61W, or H104F substitutions adjacent to the most flexible propionate. The 3 Å crystal structure of **dOHbr-2**-L125M-L150F (*k*_cat_/*K*_M_ = (4.6 ± 0.5) × 10^5^ M^-1^ s^-1^, *k*_cat_ = 1.6 ± 0.1 s^-1^, *K*_M_ = 3.6 ± 0.4 μM; Figure 3H, bottom inset; Figure S36) showed that L150F leads to improved organization of this propionate, with the same conformation observed in three of four chains in the asymmetric unit.

We further explored the evolutionary potential of de novo hydroxylases by seeking to experimentally optimize the sequence of the modestly active **dOHbr-1** design. To this end, 20 residues lining the active site pocket were individually randomized using NNK degenerate primers. Individual variants were arrayed into 96-well plates and assayed as clarified cell lysates using a spectrophotometric assay to monitor the conversion of **BR** to resorufin. Following evaluation of the most active clones as purified proteins, beneficial mutations were subsequently combined by DNA shuffling, resulting in a **dOHbr-1-TLIA** (A68T-I71L-A75I-L171A) quadruple mutant with *k*_cat_/*K*_M_ of (8.2 ± 0.8) × 10^5^ M^-1^ s^-1^ (*k*_cat_ = 5.3 ± 0.2 s^-1^, *K*_M_ = 6.4 ± 0.6 = μM), a value in line with natural hydroxylases (Figure 3I). These activity improvements correlate with substantially improved total turnover numbers prior to enzyme deactivation (TTN = 12 and 250 for **dOHbr-1** and **dOHbr-1**-**TLIA**, respectively). The mutations introduced during evolution are clustered within the distal heme pocket, with I71L and A75I positioned adjacent to the putative binding cavity accommodating the benzyl motif of **BR**, potentially improving hydrophobic packing around the substrate.

It was recently reported that unspecific peroxygenase enzymes can utilize molecular oxygen together with reductants such as ascorbic acid in place of peroxide, leading to milder reaction conditions and preventing overoxidation reactions (*54*). Motivated by this, we tested our designed hydroxylases under similar conditions (5 mM ascorbic acid, no added H_2_O_2_) and found that design **dOHbr-2** and many of its mutants are indeed active in the BROD assay (Figure 3J), with the best performing **dOHbr-2**-Y61F-V72K providing >60 TTN. These observations suggest that designed hydroxylases could be further developed to operate under milder conditions where O_2_ rather than peroxide is supplied as the exogenous oxidant.

### Regiodivergent hydroxylation of *Estradiol*

We next asked whether de novo design could direct oxidation toward distinct sites within the same complex molecule. We chose estradiol as a target substrate and pursued parallel campaigns targeting C6α and C16α hydroxylation (Figure 4A). C6 is a relatively activated benzylic position adjacent to the aromatic A-ring, whereas C16 is an unactivated aliphatic position. Both sites are accessible through native P450 metabolism, although neither is a dominant site of estradiol oxidation (*55*). As the rigid steroid scaffold contains numerous C-H bonds, these target transformations provide a rigorous test of our ability to control regio- and stereoselectivity of hydroxylation.

**Figure 4.**
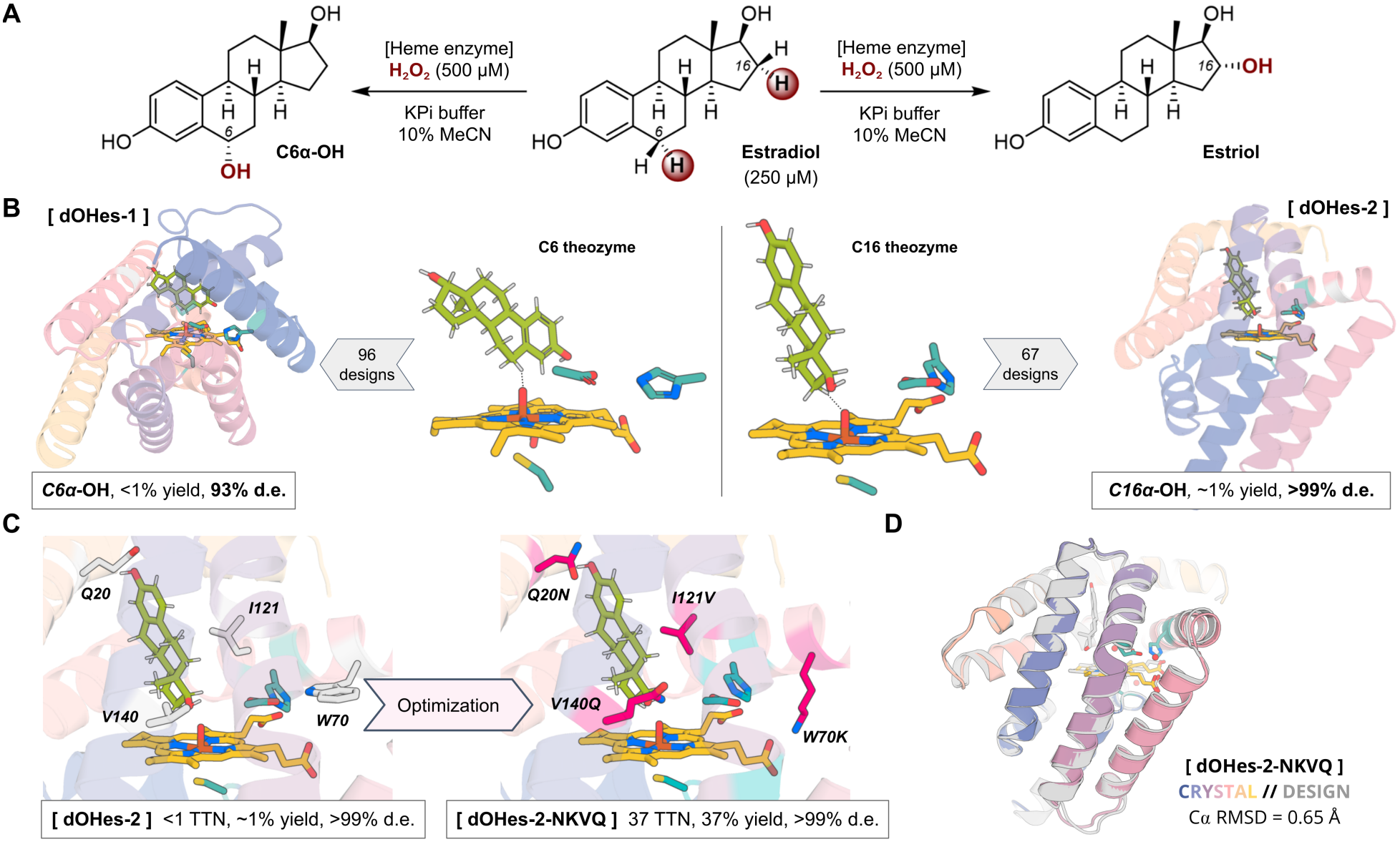
Selective C_sp3_-H hydroxylation reactions. **(A)** Regiodivergent hydroxylation of estradiol, producing 6α-hydroxyestradiol and estriol. Reactions contained 2.5 μM purified enzyme, 250 μM estradiol and 500 μM H_2_O_2_ in potassium phosphate buffer with 10% acetonitrile. **(B)** Theozyme inputs and structural models of active designs from parallel C6 and C16 hydroxylation campaigns. Screening 96 and 67 designs identified **dOHes-1** and **dOHes-2**, which yielded <1% and approximately 1% product, respectively, both with >90% diastereomeric excess (d.e.). **(C)** Active-site models of **dOHes-2** and **dOHes-2-NKVQ**, highlighting substitutions Q20N, W70K, I121V and V140Q. Combining these substitutions increased estriol yield from approximately 1% to 37% and total turnover number (TTN) from <1 to 37 while retaining >99% d.e. **(D)** Crystal structure of substrate-free, heme-bound **dOHes-2-NKVQ** at 1.9 Å resolution, closely matching the AF3 model (Cα RMSD of 0.65 Å) and confirming the predicted Lys70–heme propionate interaction.

For each target, we positioned estradiol above the heme with the desired C–H bond directed toward the iron-oxo intermediate (Figure 4B). RFdiffusion3 generated scaffolds around each catalytic motif, followed by sequence design and stringent AlphaFold3 filtering for recovery of the intended active-site geometry (see SI section 4.1 for details). For the C6 oxidation campaign (Figure S43), testing of 96 designs from 15 distinct scaffolds led to identification of **C5** (**dOHes-1**) that produced 6α-hydroxyestradiol as the major product with low (<1%) yield and 93% diastereomeric excess. From 67 designs (10 distinct scaffolds) tested in the C16 campaign (Figure S44), we identified **E5** (**dOHes-2**) that produced estriol with ca. 1% yield and 99% diastereomeric excess. Product identities were verified by LC–MS comparison with authentic standards following dansyl chloride derivatization (Figures S45-46). Despite modest initial turnover, both designs exhibited high regio- and diastereoselectivity, with little or no detectable oxidation at alternative positions.

Both designs are compact α-helical bundles with distinct substrate-binding poses above the heme (Figure 4B), with similar positioning of the Cys axial ligand, polar contacts with the heme propionates and hydrophobic packing around the heme periphery. In the 190-residue **dOHes-2**, estradiol is deeply buried, with Q20 positioned to hydrogen bond with its C3 hydroxyl group and F23 and F126 providing aromatic and hydrophobic contacts that orient C16 toward the reactive iron center (Figure S48). In the 213-residue **dOHes-1**, a shallower pose presents C6 for oxidation, with Q183 positioned to hydrogen bond with the C3 hydroxyl group (Figure S47). These distinct binding modes provide a structural rationale for the observed regiodivergence. Both **dOHes-2** and **dOHes-1** are structurally distinct from known proteins in the PDB, with both having closest-match TM scores of 0.45 (Figures S49-50).

We selected **dOHes-2** for focused mutagenesis because it combined higher initial activity with hydroxylation of the more challenging unactivated C16 position, providing a strong basis for identifying sequence features limiting catalysis. We targeted 13 positions affecting substrate access, substrate hydrogen bonding or the catalytic network, and selected 180 variants predicted by AF3 to retain the designed structure for experimental testing (Figure S51). Screening identified several beneficial substitutions (Figure S52): Q20N shortens the side chain at the tunnel-entry while preserving hydrogen-bonding capacity, while V140Q positions an amide group near the C17 hydroxyl for potential hydrogen bonding. I121V reduces steric bulk near the iron-oxo center and proton-relay network, and W70K introduces a flexible, positively charged side chain positioned to stabilize a heme propionate through electrostatic interactions. Combining these substitutions yielded **dOHes-2-NKVQ** (Q20N-W70K-I121V-V140Q), which increased estriol formation by approximately 40-fold to 37 ± 2% yield and 37 ± 2 TTN while retaining high selectivity for the target C-H bond (Figure 4C).

Crystal structures of substrate-free, heme-bound **dOHes-2** (3.6 Å resolution; Figure S54) and **dOHes-2-NKVQ** (1.9 Å resolution; Figure 4D; Figure S53) both closely match their design models (Cα RMSD of 0.72 Å and 0.65 Å, respectively), with the backbone architecture and general heme placement similar in both variants. These structures confirm that the activity gains observed in the tetramutant arise from fine-tuning of interactions within the designed active site. In the optimized variant, the catalytic residues and all four substituted side chains are positioned as observed in the model, with Lys70 contacting a heme propionate and supporting its role in cofactor binding. These substitutions suggest that refinement of substrate access and the local catalytic environment will be key to improve the activity of designed C-H hydroxylases moving forward.

## Conclusion

Our results establish a practical framework for designing a selective C–H hydroxylase around a chosen substrate and reaction geometry without relying on pre-existing enzyme scaffolds. A set of structural principles supports this chemically complex transformation, combining axial Cys coordination, heme organization, a peroxide-activating Glu–His dyad, and precise substrate positioning relative to the iron-oxo intermediate. Designing proteins focusing on these features generates site-selective catalysts without explicitly modelling every step in peroxide activation and proton transfer, which makes the design challenge more computationally tractable.

Our design approach enables aromatic and aliphatic hydroxylations across increasingly complex substrates using distinct de novo scaffolds. Crystal structures across all campaigns closely match the design models, confirming the intended backbones, heme placement, and catalytic residue positioning. On model substrates the designs achieve catalytic efficiencies reaching 10^6^ M^-1^ s^-1^, approaching the levels of natural hydroxylating enzymes. Regiodivergent and diastereoselective hydroxylation of the steroid estradiol demonstrates that oxidation sites and stereochemical outcomes can be specified at the design stage. Sequence optimization further improved activity while preserving selectivity, allowing features imperfectly captured during design to be refined experimentally. Many beneficial substitutions had interpretable roles, improving heme-propionate organization, substrate access, and second-shell interactions around the catalytic machinery. These features point to opportunities for refinement of our design principles to further improve catalytic performance and design success rates. Together, these results provide a foundation for creating hydroxylases tailored to selective C–H functionalization of chosen molecular targets.

## Data availability

Structural data supporting the findings of this study have been deposited in the Protein Data Bank (PDB). Detailed descriptions of experimental and computational procedures, Figures S1-55, Tables S1-25 are available in the supplementary information document (PDF).

## Acknowledgements

The authors would like to thank Dr. Florence J. Hardy, Dr. Brianne King, Dr. Declan Evans, Dr. Yakov Kipnis, Dr. Donghyo Kim, Avi Swartz, and Prof. Donald Hilvert for valuable discussions and insights. We are grateful for Luki Goldschmidt and Kandise Van Wormer for maintaining computational and wet lab resources at the Institute for Protein Design. We thank Dionne Vafeados, Dr. Brian Belmont, Dr. Jason Qian, and Dr. Stacey Gerben (Institute for Protein Design) for providing DNA fragments using the IPDassembly protocol. We thank Dale Whittington for assistance with mass spectrometry. We thank Dr. Jonas Wilhelm for helping improve FastMPNNdesign. We thank Dr. Jenny Cronin for assistance with project management. This work was supported by Howard Hughes Medical Institute (D.B. and I.K.); Fund for Science & Technology (J.J., S.G.); The Audacious Project at the Institute for Protein Design; The Open Philanthropy Project Improving Protein Design Fund. This work was funded by the Gates Foundation (INV-092396 and INV-092950 to A.P.G. and D.B). A.P.G. also acknowledges support from an EPSRC Centre-to-Centre Partnership with the Institute for Protein Design (EP/Z531157/1). J.J. acknowledges support by the Overseas Study Scholarship Program of Jeonnam-Gwangju Special Metropolitan City and Jeollanamdo Institute for Talent & Lifelong Education. We thank NERSC for access to the Perlmutter high-performance computing resources (Award BER-ERCAP0022018). Parts of this work was executed on Hyak, UW’s high performance computing cluster. This resource was funded by the UW student technology fee.

This work is based upon research conducted at the Northeastern Collaborative Access Team beamlines, which are funded by the National Institute of General Medical Sciences from the National Institutes of Health (P30 GM124165). The Eiger 16M detector on 24-ID-E is funded by a NIH-ORIP HEI grant (S10OD021527). This research used beamtime awards (DOI: 10.46936/APS-191329/60015048 and DOI: 10.46936/APS-195442/60017919) from the Advanced Photon Source, a U.S. Department of Energy (DOE) Office of Science User Facility operated for the DOE Office of Science by Argonne National Laboratory under Contract No. DE-AC02-06CH11357. This research used resources (FMX/AMX) of the National Synchrotron Light Source II, a US Department of Energy (DOE) Office of Science User Facility operated for the DOE Office of Science by Brookhaven National Laboratory under contract no. DE-SC0012704. The Center for BioMolecular Structure (CBMS) is primarily supported by the National Institutes of Health, National Institute of General Medical Sciences (NIGMS) through a Center Core P30 Grant (P30GM133893) and by the DOE Office of Biological and Environmental Research (KP1607011). Crystallography data were collected at NSLS II either under NECAT BAG proposal 311950.

## Author contributions

I.K. and D.B. conceptualized the study. I.K., J.J. contributed code, functions, and design pipeline development. I.K., J.J., S.G. performed computational design and analysis. I.K., J.J., S.G., A.A., B.W., R.S. performed the experiments. A.K., A.B., J.M., H.N., E.J. performed protein crystallization and data analysis. A.P.G. and D.B. supervised the research. The manuscript and supplementary information were drafted by I.K., J.J., S.G., A.A., A.P.G. and D.B. All authors reviewed and commented on the manuscript.

## Competing Interests

The authors declare no competing interests.

## Notes

### Competing Interest Statement

The authors have declared no competing interest.

