## Supplementary Information for "De Novo Design of C–H Hydroxylation Enzymes with Programmed Selectivities"

#### Contents

**1. Methods and materials** **2**
1.1. General computational methods 2
1.2. General experimental methods 4
**2. Aromatic Hydroxylase Design** **6**
2.1. Computational methods 6
2.1.1. Sequence redesign of dOHar-1 10
2.2. Experimental Methods 11
**3. Aliphatic Hydroxylase Design** **20**
3.1. Computational methods 20
3.1.1. De novo design of resorufin ether O-dealkylases 20
3.1.2. In silico optimization of dOHbr-2 34
3.2. Experimental methods 36
3.2.1. Screening and characterization of resorufin ether O-dealkylases 36
3.2.2. Optimization of dOHbr-1 40
3.2.3. Optimization of dOHbr-2 41
3.2.4. Kinetic characterization 49
3.2.5. Spectroscopic characterization 51
3.3. Supplemental figures and tables 53
**4. Estradiol hydroxylation** **60**
4.1. Computational methods 60
4.2. Experimental methods 69
**5. Crystallography** **80**
**6. References** **84**

### 1. Methods and materials

All materials were obtained from commercial suppliers and used as received, unless stated otherwise. Chemicals were sourced from Sigma-Aldrich unless stated otherwise; LB-agar, LB media and Terrific Broth II (TB-II) from MP Biomedical; *Escherichia coli* (*E. coli*) BL21 DE3 from NEB (#C2527I) or Intact Genomics (#1054-24); *E. coli* 5 alpha (DH5α), Q5 DNA polymerase, BsaI-HFv2, T4 DNA ligase, SOC medium from New England BioLabs (NEB); IPTG from Fisher Scientific; kanamycin sulfate from Thermo Fisher; resorufin benzyl ether from Ambeed or Sigma-Aldrich; N-ethyl-1,8-naphthalimide from A2Bchem; N-[(4-fluorophenyl)methyl]-1,8-naphthalimide from TargetMol, estradiol, estriol and 6α-hydroxyestradiol from MedChemExpress. Oligonucleotides and genes were synthesized by Twist Bioscience or Integrated DNA Technologies (IDT).

$^{1}H$ and $^{13}C$ NMR spectra were recorded on a Bruker Ultrashield Plus 700 ($^{1}H$ at 700 MHz, $^{13}C$ at 175 MHz) spectrometer. $^{1}H$ and $^{13}C$ spectra are referenced to residual solvent signals: DMSO-*d6* 2.50 ppm for $^{1}H$ and 39.5 ppm for $^{13}C$. Coupling constants (*J*) are reported in Hz and coupling patterns are described as d = doublet, t = triplet, q = quartet, m = multiplet, br = broad.

50/200 KPi buffer refers to a buffer containing 50 mM potassium phosphates, 200 mM NaCl at pH 7.2. At this pH the potassium phosphates composition is roughly 14.1 mM $KH_2PO_4$ and 35.9 mM $K_2HPO_4$.

#### 1.1. General computational methods

**DFT calculations**

All calculations were performed using Gaussian 16 software (*1*). Structural optimizations and frequency calculations were performed with the unrestricted U-B3LYP-D3 method along with 6-31G(d) basis set and the SDD ECP on Fe atom. D3 dispersion correction was applied using the Becke-Johnson damping function (*2*). Solvent effects of diethyl ether were included using the CPCM solvation model during optimization and single point energy calculations. This method has previously been shown to be appropriate for modeling similar systems (*3*). Frequency calculations were performed to confirm whether the structure is a minimum or a transition state. Intrinsic reaction coordinate (IRC) analysis was used to confirm that the obtained transition states connect the correct minima.

We computed the C-H abstraction transition state for aliphatic substrates (BR, IND, estradiol), and the CpdI electrophilic attack at an aromatic ring carbon transition state for aromatic substrates (NEN, F-8). We used a truncated heme model with the propionate groups cropped to ethyl groups. To mimic cysteine ligation, we used a methanethiolate proximal ligand bound to heme. We computed the oxidation transition states in all accessible heme spin states (doublet, quartet, sextet), but found no significant structural differences between them.

**Rosetta ligand parameters file creation**

The heme Compound I and the substrate molecule geometries were saved as XYZ files and subsequently converted to mol2 files using OpenBabel. The bonding information in the mol2 files was manually inspected to ensure that the entire structure is represented as a single fragment. Thereafter, mol2params.py script, available within Rosetta, was used to convert the mol2 files into Rosetta .params files. The partial charges of the carboxylate oxygen atoms of the propionate groups were adjusted in the params files from -0.74 to -1.24 to improve their treatment as good hydrogen bond acceptors during Rosetta packing and minimization. The conformational diversity of the molecules was defined in the params files using the PROTON_CHI section. For heme this was used to sample only the rotamers of the propionic acid groups. For BR substrate this was used to sample the phenyl ring and the two ether C-O bonds. For NEN substrate this was used to sample the ethyl group. For F-8 substrate this was used to sample the fluorobenzyl group.

**Theozyme construction**

The coordinates of the catalytic residues were taken from the crystal structure of a native unspecific peroxygenase, *Hsp*UPO (PDB id 7O2G) (*4*). We saved the coordinates of the Glu180-His110 catalytic dyad and the Pro38-Cys39-Pro40-Met41-Leu42 proximal heme-ligating motif into a new PDB file. M41 and L42 were mutated to alanines to scrub their sequence information while maintaining their backbone coordinates. We found that using the PCPXX motif from the UPO family served as a reliable context for scaffolding the cysteine residue, leading to high success rates in accurately recapitulating the motif in AlphaFold3 predictions. Without the backbone context (i.e., tip-atom scaffolding with RFD2/3) the cysteine was often placed in the middle of a helix with almost no general options for providing stabilizing hydrogen bond interactions. The PCPXX motif on the other hand fixes the cysteine rotamer through hydrogen bonds with the backbone amide groups of the N+2 and N+3 positions (the “XX” residues).

The distances and angles from the oxidation transition states computed with DFT were used to create a matcher/enzdes constraint file for Rosetta (*5*). The geometric constraints from the computed TS were used to define and maintain the precise relative placement of Compound I and the substrate. Additionally, the constraint files contained the definition of the heme-cysteine interaction, sampling the Fe-S bond rotation with a 90-degree period. The constraint definition and the extracted catalytic residue coordinates served as an input for building the theozyme structures. The theozyme structure PDB files were constructed using the InvRotZyme utility (*6*). In short, with this tool we placed the cysteine motif underneath the heme, the substrate next to the iron-oxo, and the Glu-His dyad near the iron-oxo, all based on the constraint definitions. Next we sampled the conformers of the heme and the substrate based on the rotable bonds defined in the corresponding params files, and saved all non-clashing combinations as separate PDB files.

#### 1.2. General experimental methods

**Size-exclusion chromatography**

Following IMAC purification, designs were further purified by SEC on ÄKTAxpress (GE Healthcare) using a Superdex Increase 75 10/300 GL column (GE Healthcare) in 50/200 KPi or TBS buffer at 0.8 mL/min flowrate. The monomeric or smallest oligomeric fractions of each run were collected. SEC retention volume to molecular weight equivalencies were calibrated with protein standards (Cytiva LMW (#28403841).

**Circular dichroism**

To determine secondary structure and thermostability of the heme proteins, far-ultraviolet circular dichroism (CD) measurements were carried out on a JASCO J-1500 instrument. The 200 to 260 nm wavelength scans were measured at every 10 °C intervals from 25 °C to 95 °C. Temperature was increased at the rate of 2 °C $min^{-1}$, and spectra were acquired after the temperature had stabilized to within 0.1 °C of target temperature for 5 seconds. Wavelength scans and temperature melts were performed using 0.40 mg $mL^{-1}$ protein in 50 mM KPi, 20 mM NaCl, pH 7.2 with a 1 mm path length cuvette. Protein concentrations were determined by absorbance at 280 nm, measured using a NanoDrop 8000 spectrophotometer (Thermo Scientific) using predicted extinction coefficients.

**Variable temperature spectrophotometric measurements**

To observe changes in the spectral properties of bound heme at increasing temperatures, UV−Vis spectra were measured of holo-proteins using the Jasco Spec V750 spectrophotometer and a 10 mm pathlength cuvette. Spectra in the 230-800 nm range were collected at every 10 °C intervals between 25 °C and 95 °C. Temperature was increased at the rate of 5 °C $min^{-1}$, and spectra were acquired after the temperature had stabilized to within 0.5 °C of target temperature for 5 seconds. Measurements were performed with 25 μM solutions of purified holoprotein in 50 mM KPi, 200 mM NaCl, pH 7.2.

**Protocol for determining hemeprotein concentration**

To quantify the protein-bounded heme concentration, UV−Vis spectra were measured of heme supplemented purified proteins using the Jasco Spec V750 spectrophotometer and a 10 mm pathlength cuvette by performing hemochrome assay (*7*). The samples were prepared by mixing 500 μL of the purified protein sample and 500 μL of solution I [0.2 M NaOH, 40% (v/v) pyridine, 0.5 mM $K_3Fe(CN)_6$]. Their UV-Vis spectra (380–620 nm) were immediately recorded for oxidized state Fe(III). To record the reduced state Fe(II), 10 μL of sodium dithionite (0.5 M solution in water) was added to the sample and their UV-Vis spectra were recorded. The heme-bounded protein concentration was calculated using the extinction coefficient and dilution factor (2× dilution in volume): $\varepsilon[557_{reduced} - 540_{oxidized}] = 23.98\ mM^{-1}\ cm^{-1}$.

**Small molecule mass spectrometry**

To validate the formation of the reaction products, the enzymatic reaction solution of **dOHar-1** was analyzed using reverse-phase hybrid quadrupole-TOF LC/MS/MS on AB SCIEX TripleTOF 5600 with ZORBAX Extend-C18, 2.1 x 50mm, 5 μm (A: $H_2O$ with 0.1% Formic Acid,

B: acetonitrile with 0.1% Formic Acid). 100 μL of enzymatic reaction solution was prepared in 5 μM of protein samples, 50 μM substrate, and 1 mM $H_2O_2$. After 1 hour of enzymatic reaction, the enzymatic reaction was stopped by adding 100 μL of acetonitrile. After centrifuging at 14,000 rpm for 20 min, the supernatant of enzyme reaction solution was filtered with 0.22 μm PES membrane filter. 50 μL of filtered reaction solution was injected to reverse-phase LC-MS.

The enzymatic reaction solution for all other samples were analyzed using reverse-phase LC-MS on Agilent 6475 LC/MS/MS with InfinityLab Poroshell 120 EC-C18, 2.1 x 100 mm, 1.9 μm (A: $H_2O$ with 0.1% Formic Acid, B: acetonitrile with 0.1% Formic Acid). 100 μL of enzymatic reaction solution was prepared in 5 μM of protein samples, 50 μM substrate, and 1 mM $H_2O_2$. After 1 hour of enzymatic reaction, the enzymatic reaction was stopped by adding 100 μL of acetonitrile. After centrifuging at 14,000 rpm for 20 min, the supernatant of enzyme reaction solution was filtered with 0.45 μm Nylon membrane filter. Estradiol and related products were derivatized by dansyl chloride (details described later). 1 μL of filtered reaction solution was injected to reverse-phase LC-MS. Chromatography was measured by a single-ion channel with the appropriate ion mass.

### 2. Aromatic Hydroxylase Design

#### 2.1. Computational methods

##### Theozyme creation

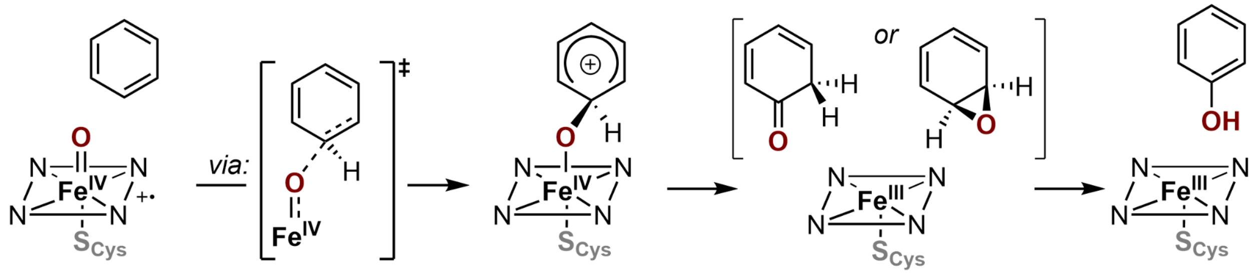


**Figure S1.** Proposed reaction mechanisms of aromatic hydroxylation. The reaction starts with an electrophilic attack of heme CpdI at the aromatic ring, forming the tetrahedral σ complex intermediate. This intermediate could then undergo either epoxide formation or the “NIH shift” by the migration of a hydrogen atom (*8*). Both intermediates ultimately lead to the formation of the phenol product either through epoxide opening or keto-enol tautomerism. We focused on the energetically more demanding first step of the reaction in our design campaign.

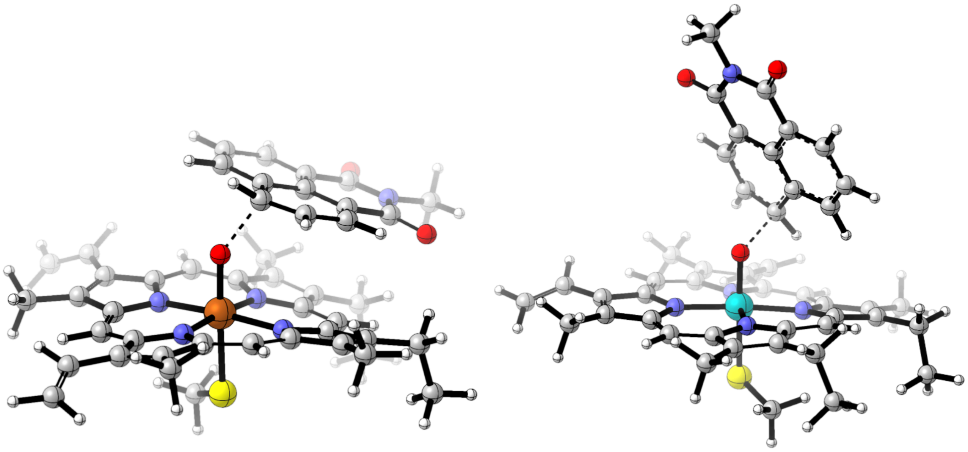

**Figure S2.** Computed transition state geometries of the initial oxidation step leading to aromatic hydroxylation of N-alkyl 1,8-naphthalimide substrate. Truncated models of heme, substrate and cysteine were used.

To design aromatic hydroxylases we used transition state geometry of the electrophilic attack of CpdI at a particular carbon atom of the aromatic substrate (Figure S1). We used DFT to compute the transition state geometries, in order to identify suitable substrate orientations relative to the heme CpdI. We identified two key substrate orientations - flat against the heme, and almost perpendicular to the heme plane (Figure S2). While the flat orientation was energetically slightly preferred, the perpendicular orientation offers more interaction opportunities with the protein environment. We built a Rosetta matcher/enzdes constraint file from the computed TS geometries and used it to construct theozyme assemblies based on the catalytic motif from *Hsp*UPO following the procedure described in Section 1.1. After filtering out catalytic motifs with clashes, 93 different motifs were selected for backbone generation.

Rosetta constraint file:

```
#block 1 for CYS coordinated to Fe
CST::BEGIN
  TEMPLATE::   ATOM_MAP: 1 atom_name: FE1 N1 C3
  TEMPLATE::   ATOM_MAP: 1 residue3:  HEO
  TEMPLATE::   ATOM_MAP: 2 atom_type: SH1
  TEMPLATE::   ATOM_MAP: 2 residue3: CYS
  CONSTRAINT:: distanceAB:    2.5   0.15  100.   1   0
  CONSTRAINT::    angle_A:   88.0   5.0  100.0  360. 0
  CONSTRAINT::    angle_B:  108.9   5.0   75.0  360. 0
  CONSTRAINT::  torsion_A:   95.0   5.0   75.0  180. 1
  CONSTRAINT:: torsion_AB:   87.9  10.0    0.0  360. 0
  CONSTRAINT::  torsion_B:  108.0  15.0   25.0  360. 0
CST::END

# block 2 for catalytic residue : Glu-His
CST::BEGIN
  TEMPLATE::   ATOM_MAP: 1 atom_type: OOC
  TEMPLATE::   ATOM_MAP: 1 residue3:  GLU
  TEMPLATE::   ATOM_MAP: 2 atom_type: Ntrp
  TEMPLATE::   ATOM_MAP: 2 residue3: HIS
  CONSTRAINT:: distanceAB:    3.0   0.2  100.   0    0
  CONSTRAINT::    angle_A:  116.0  15.0    0.0  360. 0
  CONSTRAINT::    angle_B:  126.0  15.0    0.0  360. 0
  CONSTRAINT::  torsion_A:  180.0  30.0    0.0  360. 0
  CONSTRAINT:: torsion_AB:    0.0  90.0    0.0  180. 1
  CONSTRAINT::  torsion_B:  180.0  10.0   25.0  360. 0
CST::END

################# CST_3  cofactor and substrate ##############
# block 3 for cofactor and substrate : HEO-FRE
VARIABLE_CST::BEGIN
# variable 1 : HEO-FRE
CST::BEGIN
  TEMPLATE::   ATOM_MAP: 1 atom_name: O1 FE1 N1
  TEMPLATE::   ATOM_MAP: 1 residue3:  HEO
  TEMPLATE::   ATOM_MAP: 2 atom_name: C45 C41 C35
  TEMPLATE::   ATOM_MAP: 2 residue3: FRE
  CONSTRAINT:: distanceAB:      2.2  0.2  100   0   0
CST::END
# variable 2: HEO-FRE with alternative O-C interaction
CST::BEGIN
  TEMPLATE::   ATOM_MAP: 1 atom_name: O1 FE1 N1
  TEMPLATE::   ATOM_MAP: 1 residue3:  HEO
  TEMPLATE::   ATOM_MAP: 2 atom_name: C44 C40 C34
  TEMPLATE::   ATOM_MAP: 2 residue3: FRE
  CONSTRAINT:: distanceAB:      2.2  0.2  100   0   0
CST::END
# variable 3 : HEO-NEN
CST::BEGIN
  TEMPLATE::   ATOM_MAP: 1 atom_name: O1 FE1 N1
  TEMPLATE::   ATOM_MAP: 1 residue3:  HEO
  TEMPLATE::   ATOM_MAP: 2 atom_name: C13 C9 C3
  TEMPLATE::   ATOM_MAP: 2 residue3: NEN
  CONSTRAINT:: distanceAB:      2.2  0.2  100   0   0
CST::END
# variable 4: HEO-NEN with alternative O-C interaction
CST::BEGIN
  TEMPLATE::   ATOM_MAP: 1 atom_name: O1 FE1 N1
  TEMPLATE::   ATOM_MAP: 1 residue3:  HEO
  TEMPLATE::   ATOM_MAP: 2 atom_name: C12 C8 C2
  TEMPLATE::   ATOM_MAP: 2 residue3: NEN
  CONSTRAINT:: distanceAB:      2.2  0.2  100   0   0
CST::END
VARIABLE_CST::END

# block 4 for catalytic residue : Fe-Glu
CST::BEGIN
  TEMPLATE::   ATOM_MAP: 1 atom_name: FE1
  TEMPLATE::   ATOM_MAP: 1 residue3:  HEO
  TEMPLATE::   ATOM_MAP: 2 atom_type: OOC
  TEMPLATE::   ATOM_MAP: 2 residue3: GLU
  CONSTRAINT:: distanceAB:    5.0   1.0  100.   0    0
CST::END
```

**Backbone generation**

Approximately 100,000 protein backbones were generated to scaffold the theozyme geometry using RFdiffusion2 with the following YAML input. We used unindexed motif scaffolding on the PCPX motif (positions A1-4), but found that its integrity was not always respected with RFD2. Glu (A7) and His (A6) were scaffolded with flexible primary sequence positions as well as by defining only their side-chain atoms and letting the model freely decide their full rotameric states.

RFD2 input YAML:


```
defaults:
  - base
guidepost_bonds: True
inference:
  ligand: 'HEO,FRE'
  align_motif: False
  str_self_cond: False
  state_dict_to_use: model_state_dict
  deterministic: False
  idealize_sidechain_outputs: True
idealization_metric_n_steps: 50
diffuser:
  T: 100
  trans:
    sample_schedule: linear
    exp_rate: 1
  rots:
    sample_schedule: normed_exp
    exp_rate: 10
contigmap:
  contigs: ["170-220,A1-4,A6,A7"]
  contig_atoms: "{'A1': 'CD,CG,CB,CA,N,C,O', 'A2': 'SG,CB,CA,C,O,N', 'A3': 'CD,CG,CB,CA,N,C,O', 'A4': 'CA,N,C,O',
'A6': 'CG,ND1,CD2,CE1,NE2', 'A7': 'CD,CG,OE1,OE2'}"
  length: 170-220
```


Diffused backbone filtering

All of the generated backbones were evaluated for backbone and pocket quality metrics, selecting proteins with no clashes in the backbone and with heme (excluding the propionate groups), with less than 35% loop content, no secondary structure elements pointing alone into the space, with the termini more than 15 Å away from the ligands, and with the pocket being somewhat buried (Table S1). Unideal protein backbones including clashes between the ligand and protein backbone, high loop contents, too exposed ligands, too high radius of gyration, too long helix were filtered out.

**Table S1.** Generated backbone selection criteria.

| Metric | Cutoff | Description |
|---|---|---|
| chainbreak | <= 4.5 Å | Smallest Ca-Ca distance |
| rCA_nonadj | >= 3.0 Å | Smallest non-adjacent Ca-Ca distance. Always >3.6 in the PDB. |
| lig_dist | >= 2.0 Å | Smallest distance between ligand heavyatom and protein backbone heavyatom |
| cart_bonded_avg | <= 5.0 | Average Rosetta cart_bonded score of scaffolded residues. Shows sidechain geometry quality. |
| fa_dun_avg | <= 15.0 | Average Rosetta fa_dun score of scaffolded residues. Shows rotamer quality. |
| loop_frac | <= 35% | Percentage of loops in the protein, based on Rosetta DSSP analysis. |
| longest_helix | <= 36 | Longest helix in the protein, based on Rosetta DSSP analysis. |
| rog | <= 20.0 Å | Radius of gyration of the protein. |
| term_mindist | >= 15.0 | Distance between the termini and any ligand. |
| SASA_rel | <= 0.25 | Relative SASA of the ligands ($SASA_{bound}$ / $SASA_{free}$). |

**Sequence design**

To design the protein sequence, three cycles of enhancedMPNN and Rosetta FastRelax were performed for backbones generated from RFdiffusion2. Three catalytic residues (Cys, Glu, His) were kept fixed during sequence design and their geometry was maintained by Rosetta enzyme constraints during Rosetta FastRelax. The Rosetta enzyme constraints were defined for heme binding between Cys motif and heme, the hydrogen bonding interaction between Glu and His, the substrate position relative to heme iron, catalytic Glu position relative to heme iron. The designed sequences and structures were scored using Rosetta, measuring factors describing the quality of the active site and the protein-ligand interfaces (Table S2).

<u>FastMPNNdesign protocol:</u>

```
mpnn_selector keep_hbonds_to_ligand_and_catres
mpnn 0.1 3
scale:coordinate_constraint 1.0
coord_cst_weight 1.0
scale:fa_rep 0.059
repack
scale:fa_rep 0.092
min 0.01
coord_cst_weight 0.5
scale:fa_rep 0.280
repack
scale:fa_rep 0.323
min 0.01
coord_cst_weight 0.0
scale:fa_rep 0.568
repack
scale:fa_rep 0.633
min 0.01
coord_cst_weight 0.0
scale:fa_rep 1
repack
min 0.00001
```

**Table S2.** enhancedMPNN-FastRelax selection criteria.

| **Metric** | **Cutoff** | **Description** |
|---|---|---|
| cst_score | <= 5 | Constraint relax score |
| COO_hbond | == 1.0 | Hydrogen bond between heme propionic acid side chains and protein scaffold side chains |
| HBres_not_STY | == 0 | To exclude serine, threonine, tyrosine for hydrogen bond between heme propionic acid side chains and protein scaffold |
| cms_heme | >= 420 Å | Contact molecular surface of heme |
| SASA_substrate | >= 15.0 Å | Total solvent exposure area of substrate |
| Fe_Cys_distance | <= 2.5 Å | Distance between heme iron and Cys catalytic motif |
| Fe_Glu_distance | >= 4 Å | Distance between heme iron and Glu catalytic motif |
| Fe_Glu_distance | <= 6.2 Å | Distance between heme iron and Glu catalytic motif |
| Cys_Fe_distance | <= 2.8 Å | Distance between heme iron and Cys catalytic motif |
| Glu_His_distance | <= 3.6 Å | Distance between Glu catalytic motif and HIS catalytic motif |
| cms_per_atom_substrate | >= 4.4 | Contact molecular surface of substrate |
| SASA_fluorophenyl_group | >= 5.0 $Å^2$ | Solvent exposure area of fluorophenyl group of substrate. |

**All-atom prediction model filtering**

Following the sequence design, the designs were filtered using protein-ligand co-folding model Chai-1 for the tight heme binding to protein, preorganization of the catalytic motifs, and substrate positioning. We used Chai-1 because when we began this design campaign AlphaFold3 was not yet publicly available. Designs were filtered for averaged metrics from structure prediction model: a global pLDDT > 0.85, $C_\alpha$ RMSD < 1.2 Å, catalytic residue side chain RMSD < 1.2 Å, catalytic residue pLDDT > 0.75, heme cofactor RMSD < 1.2 Å, substrate RMSD < 2.0 Å, distance between heme iron atom and carbon atom for hydroxylation < 3.5 Å, H-bond distance between Glu-His catalytic motif < 3.2 Å, minimal distance between heme iron atom and carboxylate group of Glu catalytic motif < 5.0 Å.

##### 2.1.1. Sequence redesign of dOHar-1

As the initial hit **dOHar-1** was initially designed toward **F-8**, we sought to optimize its sequence toward **NEN** using 5 cycles of enhancedMPNN-FastRelax. For enhancedMPNN input preparation, we swapped **F-8** to **NEN**, while keeping the oxidation transition state geometry. Each cycle of sequence design consists of enhancedMPNN sequence sampling, sequence mutations, and FastRelax. For enhancedMPNN sequence sampling, Cys and Met was excluded for sequence redesign except the catalytic motif to avoid unintended heme coordination. The residues forming hydrogen bond interaction with substrate was fixed for the next sequence design cycle with probability of 0.75. Apolar bias was applied for binding pocket residue (8 Å distance from iron) by adding residue-wise bias -1.0 on residues KREQYDNWH. The designed sequences and structures were scored using Rosetta, measuring factors describing the quality of the active site and the protein-ligand interfaces (Table S2).

#### 2.2. Experimental Methods

**Protein expression and purification for aromatic hydroxylase initial screening**

DNA oligos encoding the designed protein sequences were obtained from IDT as eBlocks. The overall initial screening protocols followed Semi-Automated Protein Production (SAPP) protocol (*9*). Linear DNA fragments encoding the designs were cloned into a vector containing a C-terminal Strep-tag, giving the final expressed sequence as `MSG-<design>-GSAWSHPQFEK`. After cloning, the plasmids were transformed into BL21(DE3) cells, incubated in SOC medium for 2 hours at 37 °C, and grown overnight at 37 °C in 1 mL of LB media containing 50 µg/ml of kanamycin. For protein expression, 4x100 µL of overnight LB culture was used to inoculate 4x1 mL of TB-II media and grown for 2 hours at 37 °C while shaking at 1300 rpm, followed by induction of protein expression by the addition of 1 mM IPTG. The bacterial cell cultures were incubated at 37 °C with shaking for 4 hours. The cells were collected by centrifugation at 4100g for 20 min, and the supernatant was removed. The cell pellets were stored frozen at -80 °C until cell lysis. For cell lysis, the frozen cell pellet was thawed and resuspended in 500 µL of buffer containing 50 mM KPi (pH 7.2), 200 mM NaCl, 0.1 mg/mL lysozyme, 0.01 mg/mL deoxyribonuclease I (DNase I), 0.5 mg/mL polymyxin B and Pierce protease inhibitor (1 tablet per 100 mL). After 10 minutes of shaking at 37 °C, 100 µM of hemin was added to the lysis reaction, and it was shaken at 37 °C for another 50 minutes. The plates were centrifuged at 4100g for 30 min and the supernatant was collected for further protein purification. The cleared cell lysate was purified by StrepTag affinity chromatography with 200 µL of 50% Strep-Tactin® resin suspension in a 96-well 25 µm fritted plate. The wash buffer was prepared with 50 mM KPi and 200 mM NaCl. The elution buffer was prepared with 50 mM KPi, 200 mM NaCl, and 50 mM biotin, at pH 7.2. The resin was equilibrated with 3x500 µL of wash buffer using a vacuum manifold. The clarified lysate was gently added to each well in the fritted plate. The fritted plate was covered with an aluminum seal and inverted several times to resuspend the resin. The resuspended resin was allowed to settle at room temperature for 15 minutes. The flowthrough was collected by centrifugation (1500g, 2 min). The protein-loaded resin was washed 3 times with 500 µL of wash buffer, using centrifugation. Finally, the protein was eluted off the resin using 200 µL of elution buffer using centrifugation (1500g, 2 min). The eluted protein plate was covered with an aluminum seal and stored at 4 °C until further use.

**Protein expression and purification for redesign**

Synthetic DNA fragments encoding the redesigned **dOHar-1** sequences were obtained from Twist Bioscience as gene fragments and cloned into the pDTstrep1 vector containing a C-terminal Strep-tag, using Gold Gate Assembly. The same protein expression (IPTG induction) and purification (StrepTactin affinity chromatography) procedures were used as described in the initial screening protocol.

**Fluorescent-based activity assay**

To characterize the peroxygenase activities of the designs, the purified protein samples were incubated with fluorogenic substrate **NEN** and **F-8**. Substrates and product standards were prepared in N,N-Dimethylformamide (DMF). Enzyme activity was screened in 50 µL reaction volumes, 5 µM designed protein, 50 µM substrate dissolved in DMF, and 4mM $H_2O_2$ in 96-well half area plates. The fluorescence signal for **NEN** and **F-8** was recorded on a Synergy Neo2 plate reader under excitation 450 nm, emission 558 nm. Half area 96-well flat bottom assay plates with a non-binding surface made from black polystyrene, were purchased from Corning (#3686).

Three sets of serial dilutions of an authentic sample of the **NEN-OH** product (0.8 - 50 µM) were made in 95% reaction buffer and 10% DMF to construct a fluorescence calibration curve, where each point was measured at 25 °C in a 96-well half area plate using 50 µL. Every point was measured in triplicate and averaged over 3 minutes of measurements. The calibration curve was fitted with a linear regression equation to convert fluorescence into the product concentration of **NEN-OH**.

**Kinetics analysis**

Protein samples used in kinetic measurements were obtained from 40 mL scale *E. coli* expression cultures, followed by purification using Ni-NTA affinity chromatography size-exclusion chromatography. The concentration of hemoprotein was determined using the pyridine hemochrome assay.

50 µl of enzymatic reaction solutions were prepared in KPi buffer (40 mM potassium phosphate, 140 mM NaCl, pH 7.2) and varying concentrations of **NEN** (0.8-50 µM) in 96-well half area plates. Kinetic reactions were performed by premixing the 10 nM designed protein and **NEN** (10% final DMF), and adding them to a solution of 4 mM $H_2O_2$. Each reaction was monitored for 60 minutes at 25 °C and performed in triplicate. Fluorescence measurements were performed with Synergy Neo2 plate reader. The enzymatic activity progression curves were analyzed by first subtracting the fluorescence of the no-enzyme background reaction. The resulting fluorescence curve was manually set to start at y=0 fluorescence units such that a linear regression (`y = mx + b`) could be calculated for the linear initial velocity with `b=0` (i.e., only the slope, `m`, needed to be determined), using *scipy.stats.linregress* function in the Python *SciPy* library. The calculated slopes for the initial velocities at each substrate concentration point were transformed to be expressed in terms of [product]/[enzyme] using the calibration curve (Figure S9) and known enzyme concentration. The steady-state parameters $k_{cat}$ and $K_M$ were determined by nonlinear regression fitting all data points to the Michaelis-Menten equation using *scipy.optimize.curve_fit* : $v_o/[E]_o = k_{cat}[S]/(K_M + [S])$, where $v_o$ denotes the initial velocity, $[E]_o$ the enzyme concentration, and [S] the substrate concentration. Standard deviations were calculated from the standard error of the fit considering each triplicate value as an individual data point.

**Small molecule mass spectrometry**

For **dOHar-1,** enzymatic reaction solutions were analyzed using reverse-phase hybrid quadrupole-TOF LC/MS/MS on AB SCIEX TripleTOF 5600 with ZORBAX Extend-C18, 2.1x50 mm, 5 µm) and a 50 µL injection volume. Mobile phases A and B were water with 0.1% formic acid and acetonitrile with 0.1% formic acid, respectively, with a flow rate of 0.3 mL $min^{-1}$. The gradient was held at 5% B for 1 min. The gradient was linearly increased from 5% B to 100% B for 4.5 min, and held at B 100% for 2.5 min.

For **dOHar-2,** enzymatic reaction solutions were analyzed using reverse-phase LC-MS on Agilent 6475 LC/MS/MS with an InfinityLab Poroshell 120 EC-C18 column (2.1 × 100 mm, 1.9 µm) and a 1 µL injection volume. Mobile phases A and B were water with 0.1% formic acid and acetonitrile with 0.1% formic acid, respectively, with a flow rate of 0.5 mL min$^{-1}$. The gradient was linearly increased from 10% B to 70% B for initial 4.7 min, linearly increased to 95 % B until 5.7 min, holded at 95% over 2 min.

##### Synthesis of *N*-ethyl 4-hydroxy-1,8-naphthalimide (NEN-OH)

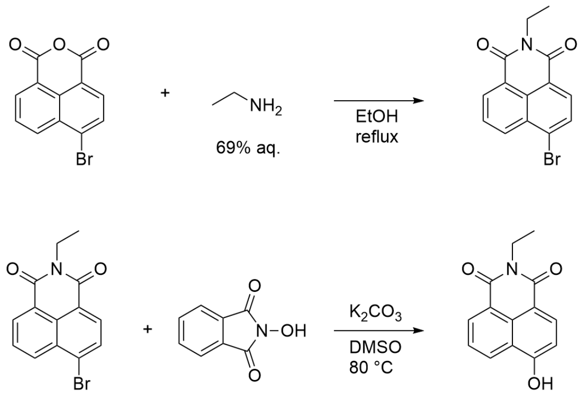


*crude*

Prepared following published protocols (*10*, *11*). In short, the intermediate *N*-ethyl 4-bromo-1,8-naphthalimide was synthesized by refluxing 4-bromo-1,8-naphthalic anhydride (277 mg, 1 mmol) with ethylamine (1.2 mmol, as 69% aq. solution) in ethanol (20 mL) for 3 hours. After cooling to room temperature, the precipitate was isolated by filtration and washed with cold ethanol (20 mL) to obtain the crude product that was used directly in the next step without further purification.

Next, the dried crude product was mixed with *N*-hydroxyphthalimide (392 mg, 2.4 mmol) and $K_2CO_3$ (415 mg, 3 mmol), in DMSO (40 mL) and stirred at 80 °C for 5 hours. After cooling, the mixture was poured into cold water (200 mL) and the pH was adjusted to 5-6. The precipitate was filtered and washed with ice-cold water. The crude product was purified using the Biotage flash chromatography system using DCM/MeOH (10:1) as the mobile phase. Poor solubility of NEN-OH in the solvent system caused significant tailing and co-elution with impurities, and as a result only 21 mg of product (7% yield) with sufficient purity was isolated. $^1$H NMR (700 MHz, DMSO-*d6*) δ 11.85 (br, 1H), 8.53 (dd, *J* = 8.3, 1.3 Hz, 1H), 8.47 (dd, *J* = 7.2, 1.3 Hz, 1H), 8.35 (d, *J* = 8.2 Hz, 1H), 7.75 (dd, *J* = 8.3, 7.3 Hz, 1H), 7.15 (d, *J* = 8.2 Hz, 1H), 4.05 (q, *J* = 7.1 Hz, 2H), 1.19 (t, *J* = 7.1 Hz, 3H). $^{13}$C NMR (176 MHz, DMSO-*d6*) δ 163.5, 162.8, 160.3, 133.5, 131.1, 129.2, 128.9, 125.6, 122.4, 121.9, 112.6, 110.0, 34.4, 13.2. These data are in agreement with those reported previously in the literature (*10*).

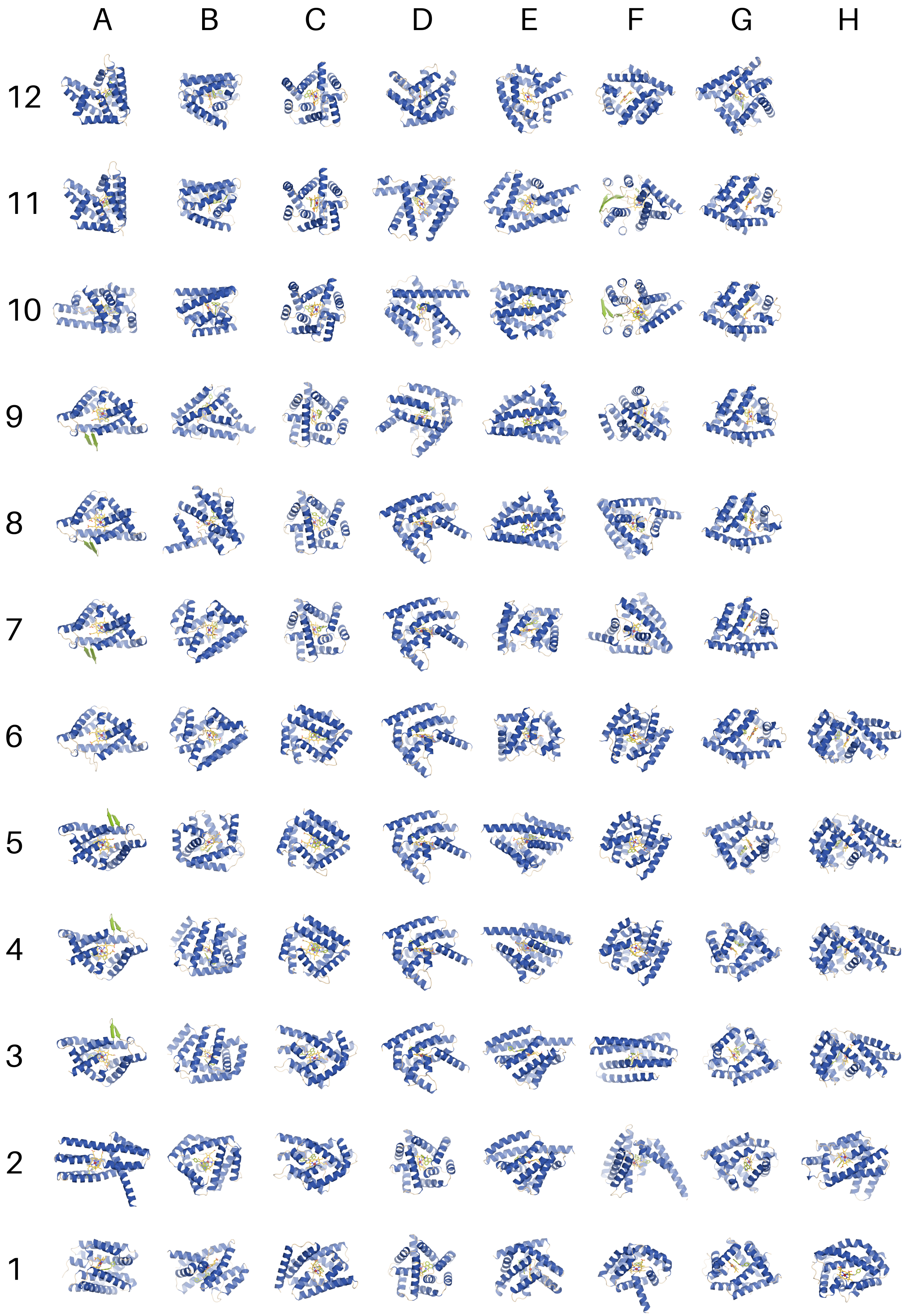


**Figure S3.** Panel of the 90 experimentally characterized designs from 29 unique scaffolds.

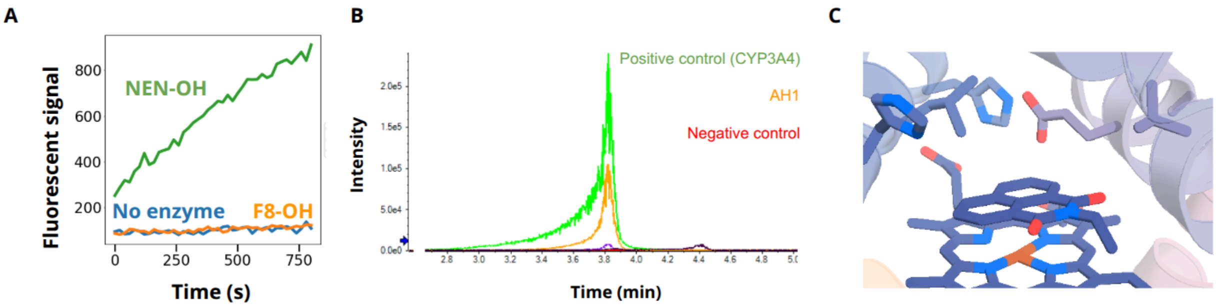


**Figure S4. (A)** Initial aromatic hydroxylase activity screen with the heme-loaded StrepTag purified design **dOHar-1**. The aromatic hydroxylation reactions of 50 µM **NEN** (green) and 50 µM **F-8** (yellow) were monitored by the fluorescence signal. The blue progress curve displays the reaction without designed protein, serving as negative control. **(B)** Mass spectrometric analysis (Q-TOF) of enzymatic reaction solution of design **dOHar-1**. 5 µM **dOHar-1** was incubated with 50 µM **NEN** for 1 hour at room temperature, after which the enzymatic reaction was quenched with 100 µL acetonitrile. A buffer with 50 µM **NEN** was used as negative control. **(C)** Chai-1 prediction of **dOHar-1** with **NEN** substrate.

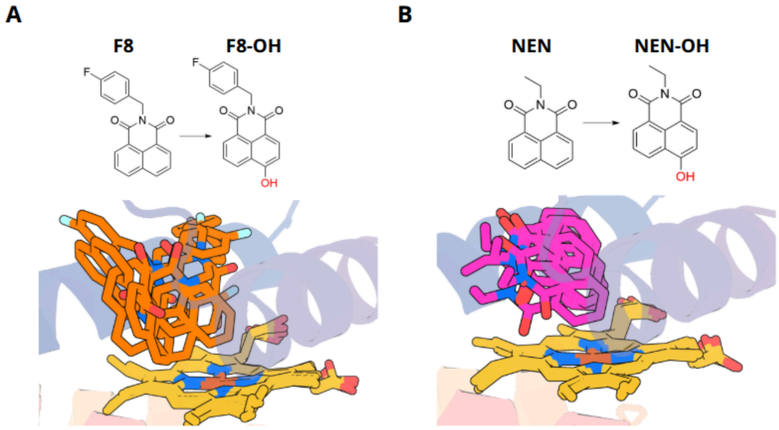


**Figure S5.** AlphaFold3 predictions of **dOHar-1** together with (A) **F-8**, and (B) **NEN**. Predictions with the larger fluorophenyl-substituted **F-8** show significant fluctuations in the substrate positioning, and rare placement of the desired carbon atom near the heme. Predictions with **NEN** are on the other hand more consistent, and show the desired carbon atom being close to heme.

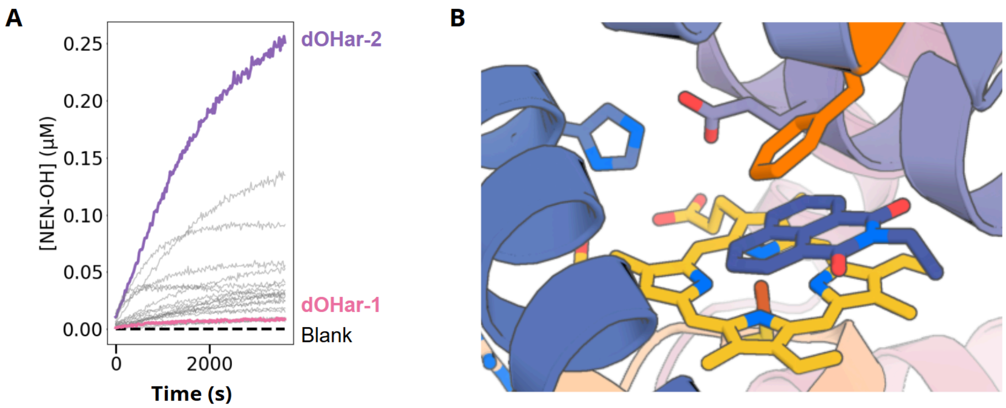


**Figure S6. (A)** Aromatic hydroxylase activity screen of redesigned variants of **dOHar-1**, showing the significantly increased activity of **dOHar-2** under screening conditions. **(B)** Active site of dOHar-2 showing Cys-ligated heme (yellow), catalytic residues (blue), and Phe residue (Orange) interacting with the aromatic ring of **NEN**.

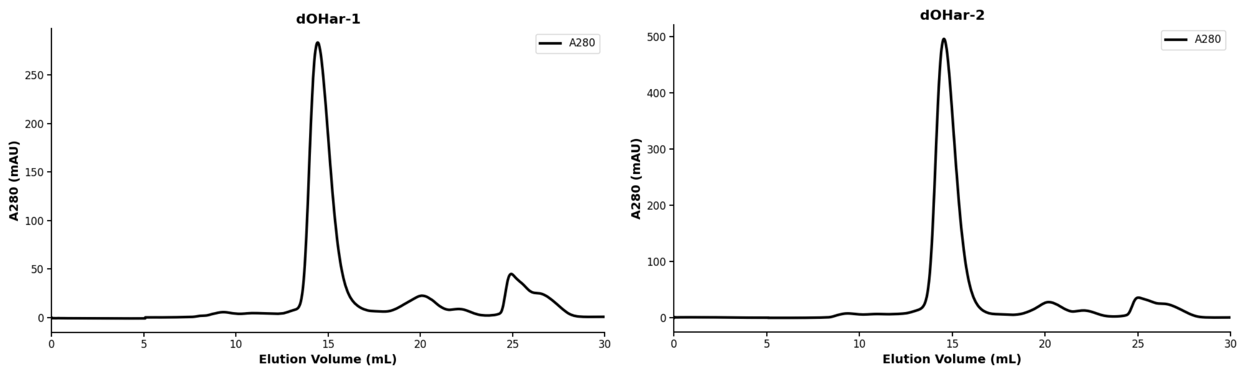


**Figure S7.** Size-exclusion chromatograms of purified **dOHar-1** and **dOHar-2**, using Superdex Increase 75 10/300 GL column and 50 mM KPi 200 mM NaCl pH 7.2 running buffer. The black trace corresponds to a chromatogram based on absorption at 280 nm.

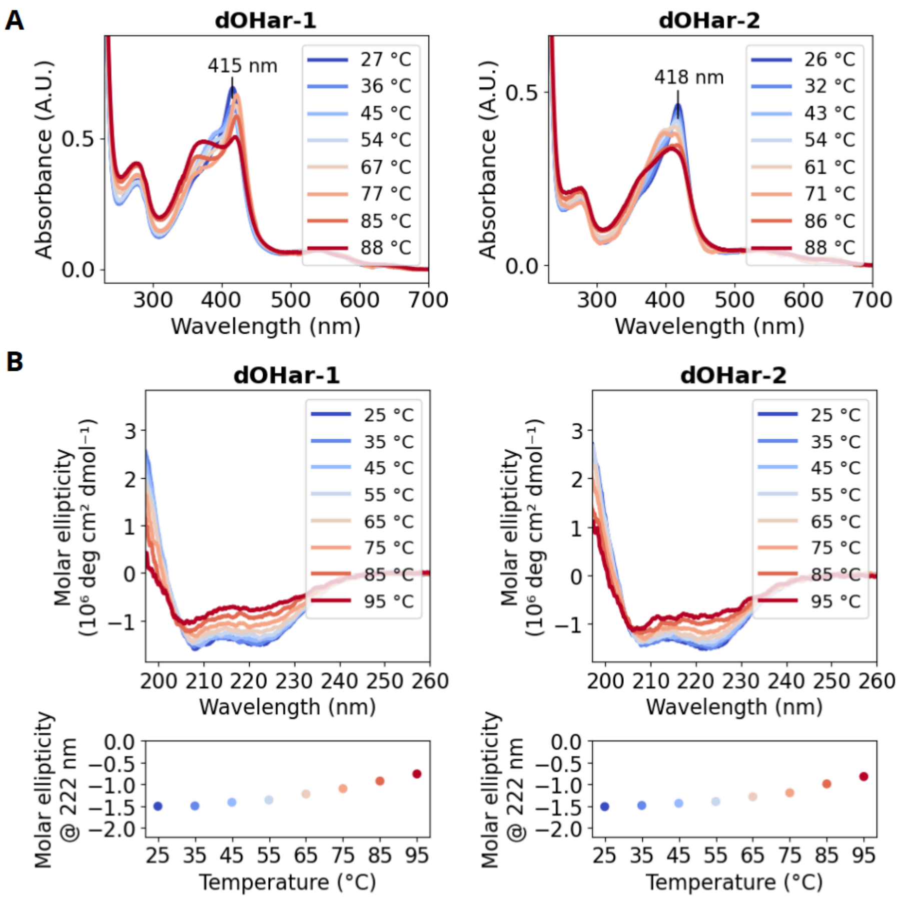


**Figure S8.** Thermal stability characterization of design **dOHar-1** and **dOHar-2**. (A) UV-Vis spectrum across different temperatures from 24 °C to 88 °C (B) Circular dichroism (CD) spectroscopy across different temperatures from 25 °C to 95 °C.

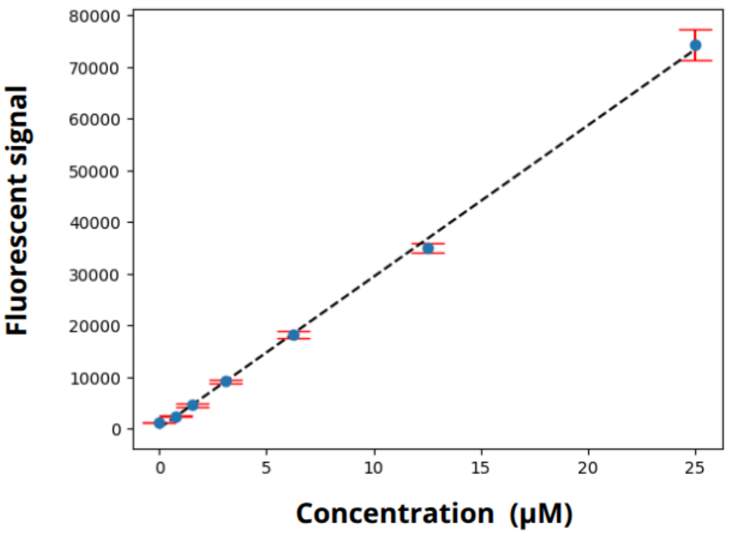


**Figure S9.** Calibration curve of **NEN-OH** fluorescence at concentrations 0.8-25 μM. ($r^2$= 0.9994)

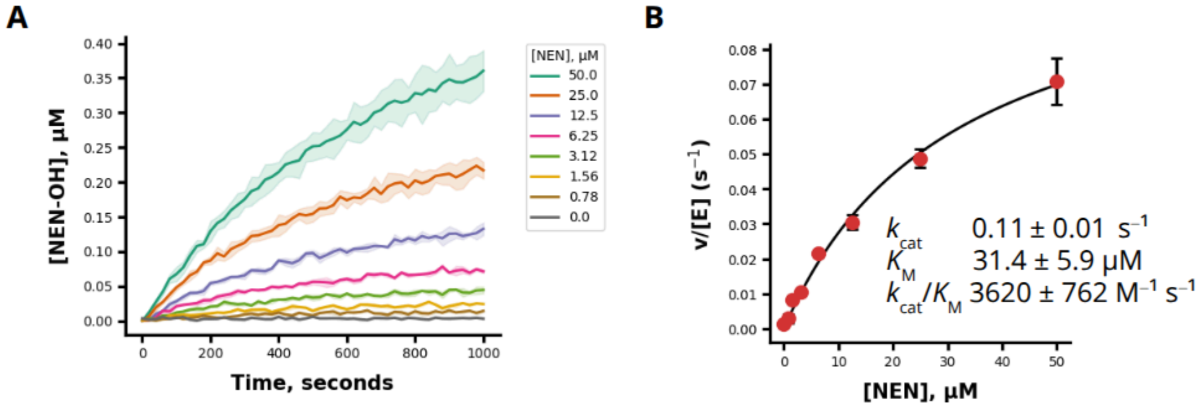


**Figure S10.** Aromatic hydroxylase activity of **dOHar-2. (A)** Enzymatic reaction progression curves for 1000 s at 0.78 μM - 50 μM of **NEN**. **(B)** Michaelis-Menten kinetic analysis plots of **dOHar-2**.

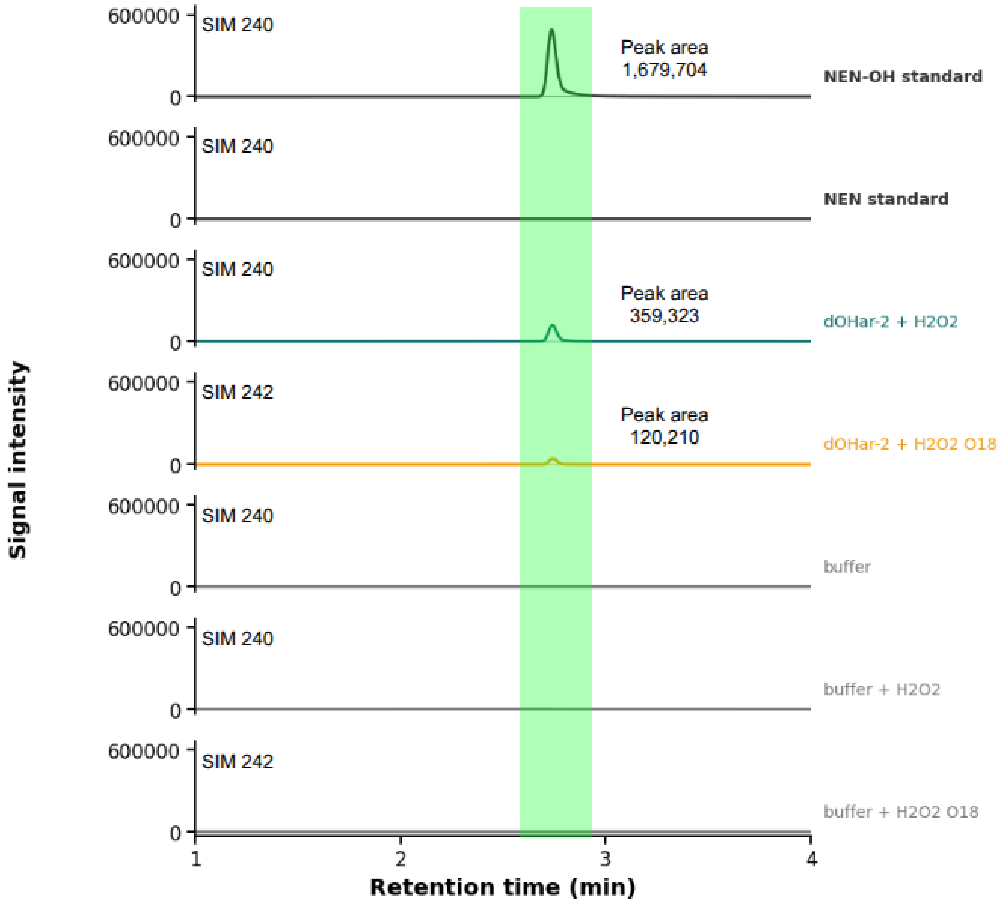


**Figure S11.** Mass spectrometric analysis (selected ion monitoring; SIM) of enzymatic reaction solution of design **dOHar-2** with $H_2O_2$ or $^{18}O$ isotope-labeled $H_2O_2$. 100 μL of 5 μM **dOHar-2** was incubated with 50 μM **NEN** and 4 mM $H_2O_2$ for 1 hour incubation at room temperature. After 1 hour incubation, the enzymatic reaction was quenched with 100 μL acetonitrile.

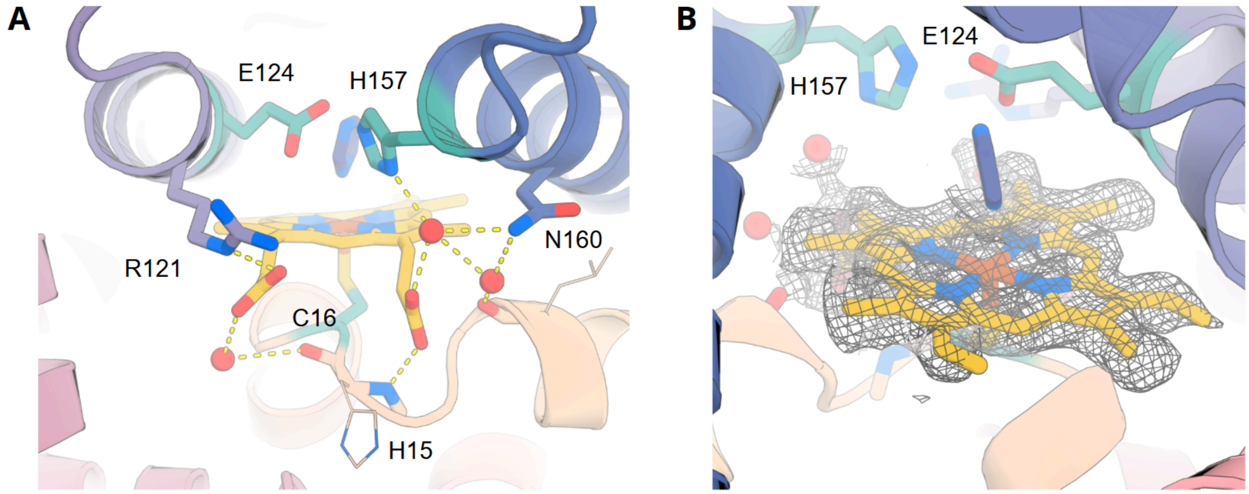


**Figure S12.** Crystal structure of **dOHar-2**. **(A)** Hydrogen bond network (yellow dashed lines) surrounding the propionate groups of heme. Crystal structure revealed many water-mediated H-bonds. **(B)** Density map of heme in the crystal structure (2*F*o–*F*c map contoured at 1σ).

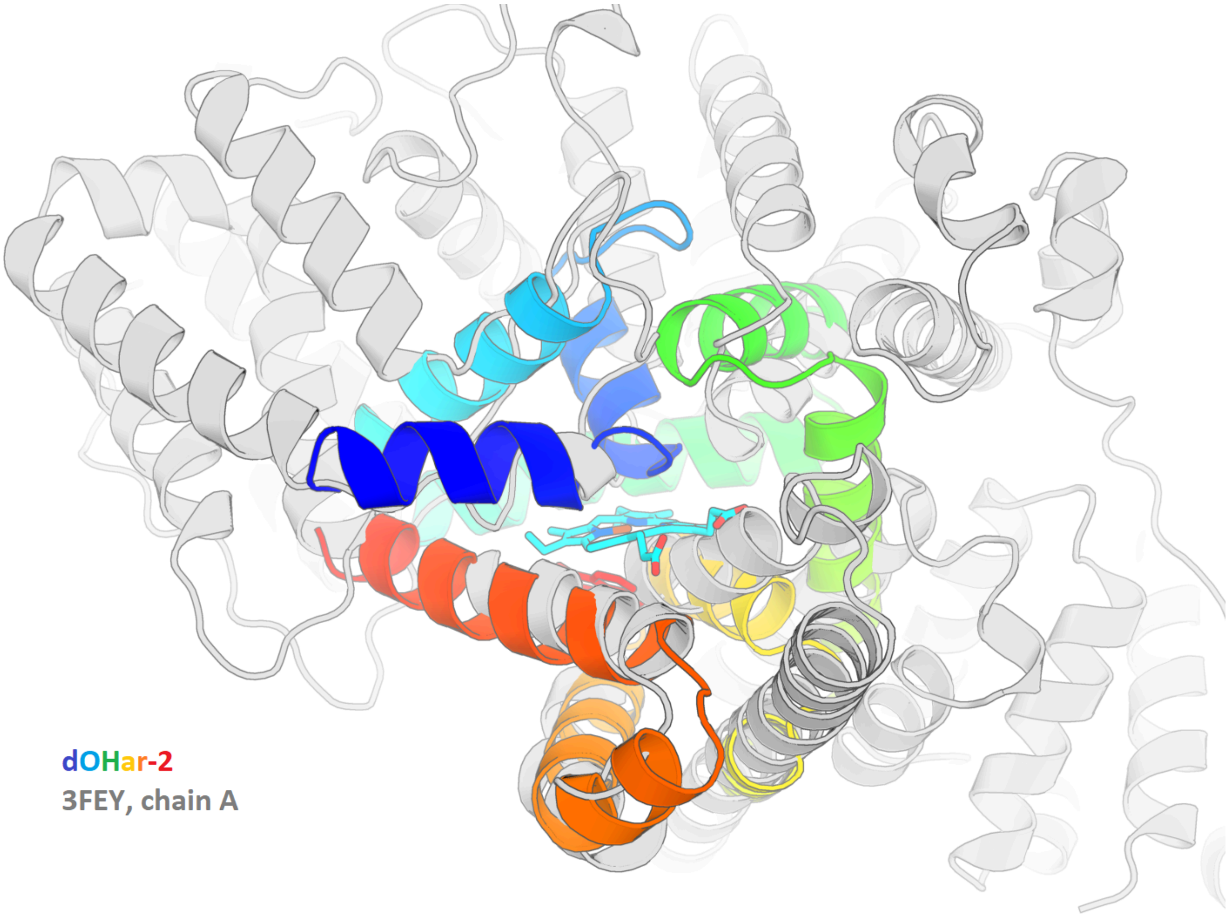


**Figure S13.** Alignment of **dOHar-2** (rainbow) with its closest match in the PDB (3FEY, chain A; gray). TM-score of 0.49 indicates low similarity between the structures, and the alignment shows that only a few individual secondary structure elements match.

### 3. Aliphatic Hydroxylase Design

#### 3.1. Computational methods

##### 3.1.1. De novo design of resorufin ether *O*-dealkylases

**Theozyme construction**

To design enzymes for the benzylic hydroxylation of resorufin benzyl ether (referred to as **BR** in the manuscript, and `BRO` in the computational workflow), we first computed the C-H abstraction transition state using DFT (Figure S14). We computed multiple distinct TS geometries based on the conformation of the substrate, but found that the key geometric parameters of the C-H abstraction within the Fe=O···H-C atoms remained largely unchanged. The distances and angles from the TS were used to create a constraint file for Rosetta relaxation, packing and minimization steps. The design input theozyme model was built by placing the substrate as a separate molecule next to Compound I, and using geometric constraints from the computed TS to define and maintain their precise relative placement. We used both enantiomeric transition states as an input for design - while the chirality does not matter for reaction outcome, it does give more options to find suitable active sites.

Designs were built around the same ferryl heme-thiolate (Compound I) theozyme used for our other peroxygenase campaigns, with the proximal cysteine and a distal Glu-His dyad taken from a native *Hsp*UPO (PDB: 7o2g). Using the InvRotZyme utility (*6*) we created a total of 251 unique assemblies, arising from different heme and substrate conformers, and 90-degree sampling of the PCPXX motif around the Fe-S bond. We used three ORI token placements for each input, sampled along the Fe=O bond vector, giving a total of 753 inputs.

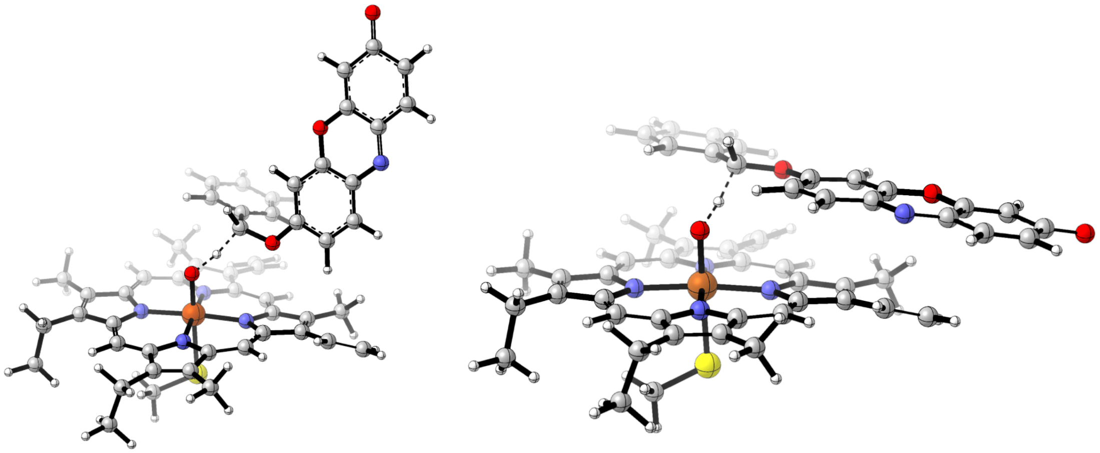

**Figure S14.** Computed transition state geometries of hydrogen atom abstraction by iron-oxo of a truncated heme model from the benzylic C-H bond of the **BR** substrate. The breaking C-H and forming O-H bonds are shown with dashed lines.

Constraint file definition for BR hydroxylase design

```
#block 1 for CYS coordinated to Fe
CST::BEGIN
  TEMPLATE::   ATOM_MAP: 1 atom_name: FE1 N4 C19
  TEMPLATE::   ATOM_MAP: 1 residue3:  HOM
  TEMPLATE::   ATOM_MAP: 2 atom_type: SH1
  TEMPLATE::   ATOM_MAP: 2 residue3: CYS
  CONSTRAINT:: distanceAB:    2.4   0.15  100.   1   0
  CONSTRAINT::    angle_A:   87.8   5.0  100.0  360. 0
  CONSTRAINT::    angle_B:  108.4   5.0   75.0  360. 0
  CONSTRAINT::  torsion_A:   98.6   5.0   25.0  180. 0
  CONSTRAINT:: torsion_AB:   86.8  30.0    0.0   90. 0
  CONSTRAINT::  torsion_B:  108.4  15.0  100.0  360. 0
CST::END

# block 2 for cofactor and substrate : HOM-BRO
VARIABLE_CST::BEGIN
CST::BEGIN
  TEMPLATE::   ATOM_MAP: 1 atom_name: O5 FE1 N4
  TEMPLATE::   ATOM_MAP: 1 residue3:  HOM
  TEMPLATE::   ATOM_MAP: 2 atom_name: C13 O9 C14
  TEMPLATE::   ATOM_MAP: 2 residue3: BRO
  CONSTRAINT:: distanceAB:     2.5  0.2  100   1   0
  CONSTRAINT::    angle_A:   130.0  5.0   75  360. 0
  CONSTRAINT::    angle_B:   115.0  5.0   75  360. 0
  CONSTRAINT::  torsion_A:   132.0 45.0    0   90. 0
  CONSTRAINT:: torsion_AB:   130.0 15.0   10  360. 0
  CONSTRAINT::  torsion_B:   125.0 10.0   25  360. 0
  ALGORITHM_INFO:: match
     SECONDARY_MATCH: DOWNSTREAM_CST 1
  ALGORITHM_INFO::END
CST::END
CST::BEGIN
  TEMPLATE::   ATOM_MAP: 1 atom_name: O5 FE1 N4
  TEMPLATE::   ATOM_MAP: 1 residue3:  HOM
  TEMPLATE::   ATOM_MAP: 2 atom_name: C13 O9 C14
  TEMPLATE::   ATOM_MAP: 2 residue3: BRO
  CONSTRAINT:: distanceAB:     2.5  0.2  100   1   0
  CONSTRAINT::    angle_A:   130.0  5.0   75  360. 0
  CONSTRAINT::    angle_B:   115.0  5.0   75  360. 0
  CONSTRAINT::  torsion_A:   132.0 45.0    0   90. 0
  CONSTRAINT:: torsion_AB:  -130.0 15.0   10  360. 0
  CONSTRAINT::  torsion_B:  -125.0 10.0   25  360. 0
  ALGORITHM_INFO:: match
     SECONDARY_MATCH: DOWNSTREAM_CST 1
  ALGORITHM_INFO::END
CST::END
VARIABLE_CST::END

# block 3 for catalytic residue : Heme-Glu
CST::BEGIN
  TEMPLATE::   ATOM_MAP: 1 atom_name: O5 FE1 N1
  TEMPLATE::   ATOM_MAP: 1 residue3:  HOM
  TEMPLATE::   ATOM_MAP: 2 atom_type: OOC
  TEMPLATE::   ATOM_MAP: 2 residue3: GLU
  CONSTRAINT:: distanceAB:    5.0   0.6  100.   0    0
  CONSTRAINT::    angle_A:  115.0  10.0   50.0  360. 0
  CONSTRAINT::    angle_B:   95.0  10.0   50.0  360. 0
  CONSTRAINT::  torsion_A:  -30.0  20.0    0.0   90. 0
  CONSTRAINT:: torsion_AB:  -75.0  15.0   25.0  360. 0
  CONSTRAINT::  torsion_B:  -30.0  15.0   25.0  360. 0
  ALGORITHM_INFO:: match
     SECONDARY_MATCH: DOWNSTREAM_CST 1
  ALGORITHM_INFO::END
CST::END

# block 4 for catalytic residue : Glu-His
CST::BEGIN
  TEMPLATE::   ATOM_MAP: 1 atom_type: OOC
  TEMPLATE::   ATOM_MAP: 1 residue3:  GLU
  TEMPLATE::   ATOM_MAP: 2 atom_type: Ntrp
  TEMPLATE::   ATOM_MAP: 2 residue3: HIS
  CONSTRAINT:: distanceAB:    3.0   0.2  100.   0    0
  CONSTRAINT::    angle_A:   97.5  20.0   25.0  360. 0
  CONSTRAINT::    angle_B:  136.0  15.0   25.0  360. 0
  CONSTRAINT::  torsion_A:  180.0  10.0   25.0  360. 0
  CONSTRAINT:: torsion_AB:    0.0  30.0    0.0  180. 0
  CONSTRAINT::  torsion_B:  180.0  10.0   25.0  360. 0
  ALGORITHM_INFO:: match
     SECONDARY_MATCH: UPSTREAM_CST 3
  ALGORITHM_INFO::END
CST::END
```

**Backbone generation**

RFdiffusion2

Using RFdiffusion2 we generated a total of 20000 backbones of 170-230 amino acids in length, with the following YAML input. We did not use unindexed motif scaffolding on the PCPXX motif (positions A1-5) because we found that its integrity was not always respected with RFD2. Only Glu (A8) and His (A6) were scaffolded with flexible primary sequence positions. The position of PCPXX in the primary sequence was therefore determined randomly in the beginning of each diffusion trajectory.

RFD2 input YAML:


```
defaults:
  - base
diffuser:
  T: 75
inference:
  num_designs: 12
  model_runner: NRBStyleSelfCond
  ligand: 'HOM,BRO'
  guidepost_xyz_as_design_bb: [True]
  contig_as_guidepost: True
  recenter_xt: True
  only_guidepost_positions: "A6-6,A8-8"
model:
  freeze_track_motif: True
contigmap:
  contigs: ["30-230,A1-5,30-230,A6-6,A8-8"]
      contig_atoms: "{'A1': 'CG,CD,CB,CA,C,O,N', 'A2': 'SG,CB,CA,C,O,N', 'A3':
'CG,CD,CB,CA,C,O,N', 'A4': 'CA,C,O,N', 'A5': 'CA,C,O,N', 'A6': 'NE2,CE1,ND1,CD2,CG,CB', 'A8':
'OE2,OE1,CD,CG'}"
  length: 170-230
```


RFdiffusion3

Using RFdiffusion3 we generated a total of 480 backbones of 175-230 amino acids in length, starting from 25 theozyme inputs. With RFD3, unindexed scaffolding of the PCPXX motif was also successful. We only used one ORI token position, centered at the iron atom. We generated outputs with a batch size of 4, in 5 batches.

RFD3 input JSON:


```
{"input": "input.pdb",
"ligand": "HOM,BRO",
"unindex": "A1-5,A6,A8",
"length": "175-230",
"Ori_token":[0.0,0.0,0.0],
"dialect": 2,
"select_unfixed_sequence": "A4-5",
"select_fixed_atoms":{
               "A1":"N,CA,C,O,CB,CG,CD",      # PRO
               "A2":"N,CA,C,O,CB,SG",         # CYS
               "A3":"N,CA,C,O,CB,CG,CD",      # PRO
               "A4":"N,CA,C,O",
               "A5":"N,CA,C,O",
               "A6":"ND1,CG,NE2,CD2,CB,CE1",  # HIS
               "A8":"OE2,OE1,CD,CG,CB",       # GLU
              "B10": "ALL",                  # heme
               "C11": "ALL"}}                 # BR substrate
```

Diffused backbone filtering

All of the generated backbones were evaluated for backbone and pocket quality metrics, selecting proteins with no clashes in the backbone and with heme (excluding the propionate groups), with less than 35% loop content, no secondary structure elements pointing alone into the space, with the termini more than 15 Å away from the ligands, and with the pocket being somewhat buried (Table S3). 1442 and 111 backbones passed the selection criteria in RFD2 and RFD3 campaigns, respectively.

**Table S3.** Generated backbone selection criteria.

| **Metric** | **Cutoff** | **Description** |
|---|---|---|
| chainbreak | <= 4.5 Å | Smallest Ca-Ca distance |
| rCA_nonadj | >= 3.0 Å | Smallest non-adjacent Ca-Ca distance. Always >3.6 in the PDB. |
| lig_dist | >= 2.5 Å | Smallest distance between ligand heavyatom and protein backbone heavyatom |
| cart_bonded_avg | <= 5.0 | Average Rosetta cart_bonded score of scaffolded residues. Shows sidechain geometry quality. |
| fa_dun_avg | <= 15.0 | Average Rosetta fa_dun score of scaffolded residues. Shows rotamer quality. |
| loop_frac | <= 35% | Percentage of loops in the protein, based on Rosetta DSSP analysis. |
| longest_helix | <= 36 | Longest helix in the protein, based on Rosetta DSSP analysis. |
| rog | <= 20.0 Å | Radius of gyration of the protein. |
| term_mindist | >= 15.0 | Distance between the termini and any ligand. |
| SASA_rel | <= 0.25 | Relative SASA of the ligands ($SASA_{bound}$ / $SASA_{free}$). |

Backbone diversification with partial diffusion

We used RFD2 and RFD3 partial diffusion to further sample the backbone space around each of the created protein scaffolds. At relatively low noise/timesteps partial diffusion generates perturbed variants of the backbones where the structural elements have shifted slightly. In the RFD2 campaign we used `partial_T` of 25 timesteps, and in the RFD3 campaign we used a `partial_T` noise level of 4.0. We fixed the positions of all atoms of the scaffolded residues and the ligands. Below are the partial diffusion YAML and JSON inputs.

RFD2 partial diffusion YAML:


```
defaults:
  - aa
diffuser:
  partial_T: 25
inference:
  num_designs: 3
  ligand: 'HOM,BRO'
  guidepost_xyz_as_design_bb: [False]
  contig_as_guidepost: True
  infer_guidepost_positions: True
  guidepost_xyz_as_design: True
  align_motif: True
model:
  freeze_track_motif: True
contigmap:
  contigs: ["41,A42-42,25,A68-68,54,A123-123,77"]  # example contigs
  length=200-200
guidepost_bonds: True
```

RFD3 partial diffusion JSON:

```
{"design_name":{"input":"diffusion_output.pdb",
"ligand":"HOM,BRO",
"contigs":"102,A103-103,49,A153-153,42,A196-196,19",  # example contigs
"length":"215-215",
"ori_token":[0.0,0.0,0.0]}}
```

The partial diffusion outputs were analyzed for the same backbone quality metrics as before (Table S3), with approximately 75% of the backbones passing the filters.

#### Iterative sequence design and structure prediction

To turn the generated backbones into proteins with a sequence that folds into the intended shape and retains the precise positioning of the heme, substrate and all catalytic residues (cysteine, glutamate and histidine), we used a four-stage design strategy. The goal is to go increasingly refined with the sequence design, and increasingly stringent with the selection strategies, to narrow in on viable design candidates.

In all stages, we used structural constraints during Rosetta repacking, minimization and relaxation. These constraints were applied using the Rosetta AddOrRemoveMatchCsts mover, and provided in the enzdes constraint format (*5*), defining the heme-Cys, heme-BR, heme-Glu, and Glu-His interaction geometries.

##### #1 Predesign

In the very first stage, we used ligandMPNN (T=0.1) to generate a single sequence for each of the diffused and partially diffused backbones. We then repacked, minimized the protein in Cartesian space, and coarsely relaxed it again in torsional space, all while applying enzdes constraints. Cartesian minimization was necessary to resolve any bond length and angle issues in the catalytic residue side-chains that were caused by the tip-atom-based scaffolding. In case of RFD3 outputs, applying the constraints also resolved any small deviations from the ideal geometry as RFD3 can slightly move the motif atoms. The structures were thereafter loosely analyzed for active site quality metrics: constraint score, ligand SASA and contact molecular surface, and ddG. The selection cutoffs were loose to only exclude the worst of the worst backbones. Overall this step retained 90% of the backbones.

At this point, the remaining unique protein backbones were manually inspected to exclude any subjectively unviable protein scaffolds. The main inspection criteria was the closedness/openness of the active site - whether the pocket is so closed that the substrate would not be able to get in, or whether the protein offers too few contacts for the substrate to be able to reliably bind. We also paid attention to undesirable scaffold features such as only loopy regions covering the substrate or the area under the heme being too unstructured or exposed - features that would be very hard to precisely control and design in later stages. The goal was to catch any unviable backbones that managed to sneak through earlier filters, and prevent spending any further computational resources on them (spending 10 seconds per backbone here can save thousands of GPU hours later on).

#2 Backbone sequence design and folding

In the second stage the goal was to identify the scaffolds that generally can be folded as designed, and loosely put the catalytic side chains into the right spot. This design stage sets the majority of the sequence for each backbone, with later stages refining only the catalytic core and its immediate surroundings. For that we generated new sequences for each of the structures from the previous step using enhancedMPNN (T=0.1,0.2,0.3; 5 sequences each), while omitting Cys and Met, applying a small bias (-0.5) against Ala and Arg, and fixing the identities of the catalytic residues. Sequences with long repeats of 5 or more of the same amino acid in a row were excluded.

The structures resulting from the generated sequences were then predicted using AlphaFold3. We did not include the ligands in the prediction at this stage, focusing purely on the folding of the *apo*-protein. The intention was to find scaffolds and sequences that are able to loosely fold into the correct shape without the aid of heme and/or the substrate. The predictions were performed in single-sequence mode (no MSA, no template), using 6 recycles and 1 diffusion sample. To find good sequence/scaffold candidates we used selection criteria listed in Table S4, focusing on protein confidence (pLDDT, pTM) and accuracy (RMSD).

Overall, in the RFD2 campaign around 1% of all sequences passed the filtering criteria, representing 29% of unique diffused backbones. In the RFD3 campaign around 7% of all sequences passed the filtering criteria, representing 50% of unique diffused backbones. Most backbones were represented by a single sequence, but a small handful had 20-35 sequences passing (from the combination of the diffused + 5 partially diffused backbones, with 15 sequences each from MPNN design). These results also illustrate the improvements in the designability of the backbones coming from RFD3, compared to RFD2. While both methods yield scaffolds for which a viable sequence can be found, then RFD3 does it with much higher success rates and at reduced computational cost.

**Table S4.** AlphaFold3 *apo* prediction selection criteria.

| **Metric** | **Cutoff** | **Description** |
|---|---|---|
| chain_ptm_ch0 | >= 0.75 | Global structural accuracy of protein chain A (predicted TM score) |
| rmsd | <= 1.5 Å | C$\alpha$ RMSD between the design model and the prediction. |
| plddt | >= 75.0 | All-atom local structural accuracy of protein chains. |
| plddt_BB | >= 80.0 | Local structural accuracy of protein chains, averaged over backbone atoms (C,N,CA,O). |
| rmsd_SR1 | <= 1.5 Å | All-atom RMSD of CYS |
| plddt_SR1 | >= 75.0 | All-atom confidence of CYS |
| rmsd_SR3 | <= 2.0 Å | All-atom RMSD of GLU |
| plddt_SR3 | >= 75.0 | All-atom confidence of GLU |
| rmsd_SR4 | <= 2.0 Å | All-atom RMSD of HIS |
| plddt_SR4 | >= 75.0 | All-atom confidence of HIS |

Finally, the successful AF3 predicted structures were aligned to the corresponding design models (enhancedMPNN inputs), the heme and the substrate directly transplanted to the AF3 model, and this structure saved as a new PDB file.

#3 Active site sequence design

In the third stage the focus is now set on optimizing the sequence of the active site. For each structure from the previous step, positions with Cα within 11 Å of any ligand heavy atom were selected as designable, excluding the catalytic residues. For design we used an iterative ligandMPNN and Rosetta repack/minimize protocol, implemented through the FastMPNNdesign package (*12*). With the protocol shown below we performed sequence design in four steps while ramping down the ligandMPNN temperature, and increasing the Rosetta fa_rep scoreterm applied to packing and minimization. Enzdes constraints were applied during all packing and minimization steps. After step 1 the top two sequences based on ranking by contact molecular surface (cms) were kept, after step 2 the top sequences ranked by cms and ddG were kept, and after step 3 the top 2 ranked by cms were kept. Using the MPNNselector feature of FastMPNNdesign we kept the identities of designed positions fixed if they introduced desired hydrogen bonds to the heme carboxylate groups. This was done after each MPNN step. This protocol yielded 10 sequences from one input. Biases were applied to LigandMPNN sequence generation against Tyr, Ala and Arg residues, and Cys and Met were omitted from design. On positions that are near the tail end of the BR substrate (close to the phenolic oxygen) we applied an additional bias (-1.0) against polar residues (KREQDNHY) to avoid the introduction of H-bond networks that would close the pocket. This decision is inspired by native P450 and UPO enzymes that have largely hydrophobic substrate access tunnels.

FastMPNNdesign protocol:

```
scale:coordinate_constraint 1.0
scale:fa_rep 0.150
mpnn 1.0 10
repack
scale:fa_rep 0.200
min 0.01
keep_best 2 cms
mpnn_selector keep_hbonds_to_ligand_and_catres
scale:coordinate_constraint 0.2
scale:fa_rep 0.8
mpnn 0.4 3
repack
min 0.01
keep_best 5 cms
keep_best 3 ddg
mpnn_selector keep_hbonds_to_ligand_and_catres
scale:coordinate_constraint 0.0
scale:fa_rep 1.0
mpnn 0.2 1
repack
min 0.01
keep_best 2 cms
mpnn_selector keep_hbonds_to_ligand_and_catres
mpnn 0.1 5
repack
min 0.00001
```

The created design models were thereafter evaluated based on various Rosetta-based metrics describing the active site (Table S5). Additionally, the design models were also relaxed after deleting the substrate, and they were then analyzed for the stability of the backbone and active site. The goal of this step was to avoid designs where the substrate is overly critical for the stability of the active site. We observed that these selection criteria retained 93% of the unique protein scaffolds (original diffusion outputs) that were used as inputs, meaning that some scaffolds are just unable to support an active site that would meet the described quality metrics.

**Table S5.** Stage #3 Rosetta filtering criteria.

| Metric | Cutoff | Description |
|---|---|---|
| all_cst | <= 4.0 | Weighted constraint score - how well are constraints respected. |
| cms_per_atom_HOM | >= 5.75 | Contact molecular surface of heme |
| cms_per_atom_BRO | >= 5.6 | Contact molecular surface of BR substrate |
| L_SASA_HOM | <= 0.05 | Relative SASA of heme |
| L_SASA_BRO | <= 0.10 | Relative SASA of BR substrate |
| substrate_SASA | >= 0.1 | Absolute SASA of BR substrate. Selecting those with at least some exposed area. |
| corrected_ddg | <= -75.0 | Rosetta ddG between the protein and all ligands (two-body energies). |
| COO_hbond | = True | Do both COO groups have at least one H-bond partner each. |
| HBres_not_STY | = True | Not all H-bonds to COO groups come from Ser, Thr, Tyr |
| YW_dist | >= 6.0 | Distance between iron-oxo and the closest Tyr/Trp sidechain |
| YW_count | <= 1.0 | Number of Tyr/Trp residues within 7 Å of iron-oxo. |
| KRE_count | <= 1.0 | Number of Lys, Arg, Glu residues touching the back of the BR substrate. Avoiding salt-bridges blocking substrate entry. |
| rmsd_CA_NoSub | <= 0.8 | Protein C$\alpha$ RMSD after relaxing without the substrate. |
| rmsd_sc_NoSub | <= 1.0 | Average pocket residue all-atom RMSD after relaxing without the substrate. |
| rmsd_HOM_NoSub | <= 0.8 | Heme RMSD after relaxing without the substrate. |
| ddg_NoSub | <= -65.0 | Rosetta ddG between the protein and heme after relaxing without the substrate. |
| cms_per_atom_NoSub | >= 5.6 | Contact molecular surface of heme after relaxing without the substrate. |

LigandMPNN was then used to generate additional sequence diversity for the designs passing the above filters. For this, we selected positions that form the second layer of residues around the active site, with C$\alpha$ within 8-12 Å from any ligand heavyatom, and its sidechains not within 5.0 Å of the ligands. We generated additional sequences (T=0.1,0.2, 5 each) with biases against Ala (-0.5) and Arg (-1.0), and omitting Cys and Met.

We then evaluated the folding of all of the sequences corresponding to passing designs (1 original + 10 diversified) using AlphaFold3. In this stage we co-folded the proteins with heme and the BR substrate, using 10 recycles and saving 5 diffusion samples. Heme was represented as the CCD_HEM model, instead of Compound I used for design. Substrate was provided as a userCCD mmCIF input with predefined atom names and bonding information. To select passing sequences we focused on the precise and confident recapitulation of the catalytic residue and heme binding geometries, but did not yet filter the predictions for substrate binding (Table S6). The intention was to identify high quality heme-binding sites first, and then further refine the substrate binding site, given an updated AF3-predicted protein structure. The selection criteria for protein and side chain quality are now much more stringent than before. Most metrics are analyzed as an average across the five diffusion samples from AF3, and the RMSDs are also filtered based on the worst value among the five models, ensuring that the sequence can be folded reliably in the same geometry. Overall, these filtering criteria retained 18% of input sequences, and 89% of unique scaffolds. The latter is notable, indicating that once the foldability of a backbone is established in an earlier stage, it is possible to turn most of them into good *in silico* heme binders.

**Table S6.** AlphaFold3 *holo* prediction selection criteria for heme-binding in stage #3.

| Metric | Cutoff | Description |
|---|---|---|
| chain_ptm_ch0 | >= 0.80 | Global structural accuracy of protein chain A (predicted TM score), average of 5 models |
| chain_iptm_ch0 | >= 0.75 | Interface structural accuracy of protein chain A, average of 5 models |
| chain_ptm_ch1 | >= 0.75 | Global structural accuracy of heme (predicted TM score), average of 5 models |
| chain_iptm_ch1 | >= 0.65 | Interface structural accuracy of heme, average of 5 models |
| chain_pair_pae_min_inter_max | <= 5.0 | Largest interchain minPAE value (average of off-diagonal elements in PAE matrix) from 5 models. |
| rmsd | <= 1.2 Å | Cα RMSD between the design model and the prediction, average of 5 models |
| plddt | >= 82.0 | All-atom local structural accuracy of protein chains. |
| plddt_BB | >= 87.0 | Local structural accuracy of protein chains, averaged over backbone atoms (C,N,CA,O). |
| lig0_rmsd | <= 2.0 Å | Heme RMSD, average of 5 models |
| lig0_rmsd_max | <= 2.5 Å | Worst heme RMSD among 5 models. |
| lig0_plddt | >= 80.0 | Heme pLDDT, average of 5 models |
| rmsd_SR1 | <= 1.0 Å | All-atom RMSD of CYS, average of 5 models |
| rmsd_SR1_max | <= 1.2 Å | Worst CYS RMSD among 5 models. |
| plddt_SR1 | >= 85.0 | All-atom confidence of CYS |
| rmsd_SR3 | <= 1.6 Å | All-atom RMSD of GLU |
| rmsd_SR3_max | <= 2.0 Å | Worst GLU RMSD among 5 models. |
| plddt_SR3 | >= 85.0 | All-atom confidence of GLU |
| rmsd_SR4 | <= 1.6 Å | All-atom RMSD of HIS |
| rmsd_SR4_max | <= 2.2 Å | Worst HIS RMSD among 5 models. |
| plddt_SR4 | >= 85.0 | All-atom confidence of HIS |

Finally, the successful AF3 predicted structures (all five samples) were aligned to the corresponding design models, the substrate directly transplanted from the design to the AF3 model, and this structure saved as a new PDB file.

<u>#4 Refinement of the substrate binding site</u>

In the fourth stage we focused exclusively on more deeply sampling the sequence space around the substrate. For this we selected the positions within 9.0 Å of the ligands, and excluded those that are closer to CYS than to the substrate (basically, underneath the heme). The design was performed using the same protocol as highlighted in stage #3, including the second layer diversification with ligandMPNN. Slightly more stringent Rosetta selection criteria were applied to the design models (Table S7).

**Table S7.** Stage #4 Rosetta filtering criteria.

| Metric | Cutoff | Description |
|---|---|---|
| all_cst | <= 3.0 | Weighted constraint score - how well are constraints respected. |
| cms_per_atom_HOM | >= 6.0 | Contact molecular surface of heme |
| cms_per_atom_BRO | >= 6.0 | Contact molecular surface of BR substrate |
| L_SASA_HOM | <= 0.05 | Relative SASA of heme |
| L_SASA_BRO | <= 0.10 | Relative SASA of BR substrate |
| substrate_SASA | >= 0.01 | Absolute SASA of BR substrate. Selecting those with at least some exposed area. |

| | | |
|---|---|---|
| corrected_ddg | <= -80.0 | Rosetta ddG between the protein and all ligands (two-body energies). |
| COO_hbond | = True | Do both COO groups have at least one H-bond partner each. |
| HBres_not_STY | = True | Not all H-bonds to COO groups come from Ser, Thr, Tyr |
| YW_dist | >= 6.0 | Distance between iron-oxo and the closest Tyr/Trp sidechain |
| YW_count | <= 1.0 | Number of Tyr/Trp residues within 7 Å of iron-oxo. |
| KRE_count | <= 1.0 | Number of Lys, Arg, Glu residues touching the back of the BR substrate. Avoiding salt-bridges blocking substrate entry. |
| rmsd_CA_NoSub | <= 0.8 | Protein C$\alpha$ RMSD after relaxing without the substrate. |
| rmsd_sc_NoSub | <= 1.0 | Average pocket residue all-atom RMSD after relaxing without the substrate. |
| rmsd_HOM_NoSub | <= 0.8 | Heme RMSD after relaxing without the substrate. |
| ddg_NoSub | <= -68.0 | Rosetta ddG between the protein and heme after relaxing without the substrate. |
| cms_per_atom_NoSub | >= 5.8 | Contact molecular surface of heme after relaxing without the substrate. |

We then evaluated the folding of all of the sequences corresponding to passing designs (1 original + 10 diversified) using AlphaFold3. We co-folded the proteins with heme and the BR substrate, using 10 recycles and saving 5 diffusion samples. To select passing sequences we focused on the precise and confident recapitulation of the entire active site, including catalytic residues, heme and the substrate (Table S8). Most metrics are analyzed as an average across the five diffusion samples from AF3, and the RMSDs are also filtered based on the worst value among the five models, ensuring that the sequence can be folded reliably in the same geometry. Overall, these filtering criteria retained 0.6% of input sequences, but still 75% of unique scaffolds. Filtering for substrate RMSD removed by far the most sequences, with a pass rate of 2%. This points to the significant challenge in finding sequences that can support precise positioning of the substrate within the active site, and why an iterative and increasingly stringent approach is justified.

**Table S8.** AlphaFold3 *holo* prediction selection criteria in stage #4.

| Metric | Cutoff | Description |
|---|---|---|
| chain_ptm_ch0 | >= 0.85 | Global structural accuracy of protein chain A (predicted TM score), average of 5 models |
| chain_iptm_ch0 | >= 0.80 | Interface structural accuracy of protein chain A, average of 5 models |
| chain_ptm_ch1 | >= 0.75 | Global structural accuracy of heme (predicted TM score), average of 5 models |
| chain_iptm_ch1 | >= 0.70 | Interface structural accuracy of heme, average of 5 models |
| chain_iptm_ch2 | >= 0.70 | Interface structural accuracy of substrate, average of 5 models |
| chain_pair_pae_min_inter_max | <= 5.0 | Largest average interchain minPAE value (average of off-diagonal elements in PAE matrix) from 5 models. |
| rmsd | <= 1.2 Å | C$\alpha$ RMSD between the design model and the prediction, average of 5 models |
| plddt | >= 85.0 | All-atom local structural accuracy of protein chains. |
| plddt_BB | >= 87.0 | Local structural accuracy of protein chains, averaged over backbone atoms (C,N,CA,O). |
| lig0_rmsd | <= 2.0 Å | Heme RMSD, average of 5 models |
| lig0_rmsd_max | <= 2.5 Å | Worst heme RMSD among 5 models. |
| lig0_plddt | >= 80.0 | Heme pLDDT, average of 5 models |
| lig1_rmsd | <= 2.0 Å | Substrate RMSD, average of 5 models |
| lig1_rmsd_max | <= 2.5 Å | Worst substrate RMSD among 5 models. |
| lig1_plddt | >= 70.0 | Substrate pLDDT, average of 5 models |

| | | |
|---|---|---|
| rmsd_SR1 | <= 1.0 Å | All-atom RMSD of CYS, average of 5 models |
| rmsd_SR1_max | <= 1.2 Å | Worst CYS RMSD among 5 models. |
| plddt_SR1 | >= 85.0 | All-atom confidence of CYS, average of 5 models |
| rmsd_SR3 | <= 1.6 Å | All-atom RMSD of GLU, average of 5 models |
| rmsd_SR3_max | <= 2.0 Å | Worst GLU RMSD among 5 models. |
| plddt_SR3 | >= 85.0 | All-atom confidence of GLU, average of 5 models |
| rmsd_SR4 | <= 1.6 Å | All-atom RMSD of HIS, average of 5 models |
| rmsd_SR4_max | <= 2.2 Å | Worst HIS RMSD among 5 models. |
| plddt_SR4 | >= 85.0 | All-atom confidence of HIS, average of 5 models |

**Final analysis of designs**

Constrained relax

All models that passed the AF3 selection were subsequently relaxed with Rosetta FastRelax. Heavy backbone constraints were applied to minimize any backbone movement and retain the AF3-predicted backbone geometry. Enzdes constraints were applied to nudge the catalytic residues, heme and the substrate closer to idealized orientation (they were already quite close coming from AF3 due to relatively strict RMSD filters). The relaxed structures were again evaluated for scores describing the active site and interactions with the ligands. The purpose of this analysis was to exclude designs where the AF3 prediction would not let the active site adopt the intended geometry (i.e. those having a poor constraint score), or would change some designed interactions with the heme (loss of H-bonds to COO) or the substrate (poor contact molecular surface). This was done for all five AF3 samples, giving each sequence multiple shots to provide a good structural model. The relaxed models were filtered based on the same criteria highlighted in Table S7, excluding the no-substrate-relax scores. Overall 47% of the designs passed this step, with the biggest failure mode being the constraint score.

Heme-only AlphaFold3 predictions

To further ensure that the active site remains preorganized without the substrate, and protein folding isn't substrate-dependent we performed additional AlphaFold3 predictions of passing designs, omitting the BR ligand. The predictions were performed with 10 recycles and 5 diffusion samples, representing heme as CCD_HEM, and without using an MSA or templating. The designs were selected based on the same filtering criteria as in stage #4 (Table S8), except for metrics related to BR substrate ("ch2" and "lig1" scores). 45% of the designs passed this stage, with the biggest failure mode being the pLDDT of heme, suggesting that co-folding with the substrate may have increased the model's confidence in placing the heme.

PLACER analysis

We used PLACER to analyze the designed active sites for conformational stability. For this we docked the substrate into a cropped 600-atom active site, while keeping the heme "fixed" (heme structure and docking pose were predicted from not noised coordinates). We generated 100 samples, using the relaxed AF3 models as input. The predictions were analyzed for the accuracy (RMSD) and confidence (pRMSD) of the catalytic side chains and the substrate. For the substrate we focused on the five highest confidence predictions, since a lot of the lower confidence samples had poorly predicted geometries. We also looked at the $p_{near}$ metric of the substrate docking funnel (RMSD vs pRMSD), measuring its "funnelness" (*13*). Table S9 lists the filtering criteria that were applied to select 61% of the designs that passed. The biggest failure mode was actually the `rmsd_pocket_worst` metric which measures the worst average sidechain RMSD of some pocket residue, indicating that the pocket contains a conformationally floppy residue.

**Table S9.** PLACER analysis selection criteria.

| Metric | Cutoff | Description |
|---|---|---|
| rmsd_prmsd_lig_BRO_top5 | <= 2.0 Å | Average RMSD of the substrate, among the top 5 models ranked by pRMSD |
| plddt_top5 | >= 0.85 | Average 1D track confidence (pLDDT) of the top5 models, ranked by pLDDT |
| plddt_pde_top5 | >= 0.75 | Average 2D track confidence (pLDDT-pde) of the top5 models, ranked by pLDDT-pde |
| prmsd_top5 | <= 2.5 | Average substrate pRMSD of the top5 models, ranked by pRMSD |
| prmsd_pnear_BRO | >= 0.3 | Funnelness of the substrate RMSD-pRMSD relationship |
| rmsd_catres1 | <= 0.4 Å | Average RMSD of Cys, across 100 models |
| rmsd_catres2 | <= 1.3 Å | Average RMSD of Glu, across 100 models |
| rmsd_catres3 | <= 1.3 Å | Average RMSD of His, across 100 models |
| u_catres1 | <= 0.4 | Average uncertainty (pRMSD) of Cys, across 100 models |
| u_catres2 | <= 1.0 | Average uncertainty (pRMSD) of Glu, across 100 models |
| u_catres3 | <= 1.0 | Average uncertainty (pRMSD) of His, across 100 models |
| u_pocket | <= 0.5 | Average uncertainty (pRMSD) of all pocket residues (within 5 Å of the substrate), across 100 models |
| rmsd_pocket_worst | <= 3.0 Å | Worst average sidechain RMSD of some pocket residue |
| u_pocket_worst | <= 2.0 | Worst average pRMSD of some pocket residue |

<u>Similarity analysis</u>

The design campaigns yielded hundreds of designs from tens of unique scaffolds that had to be narrowed down. Sequence similarly and clustering analyses were used to select representative designs for experimental testing. Design pairs from the same scaffold that had fewer than 12 different residues were manually inspected to ensure that the differences are meaningful (e.g. mutations in close vicinity of the active site). For backbones with large numbers of passing sequences we first used sequence clustering (at 85% similarity), and then randomly picked 3-6 representative examples from each cluster for manual inspection and further similarity analysis. Finally we selected 140 designs (24 scaffolds) from the RFD2 campaign and 36 designs (9 scaffolds) from the RFD3 campaign for experimental testing. Figures S15-16 show the diversity of scaffolds in the test set.

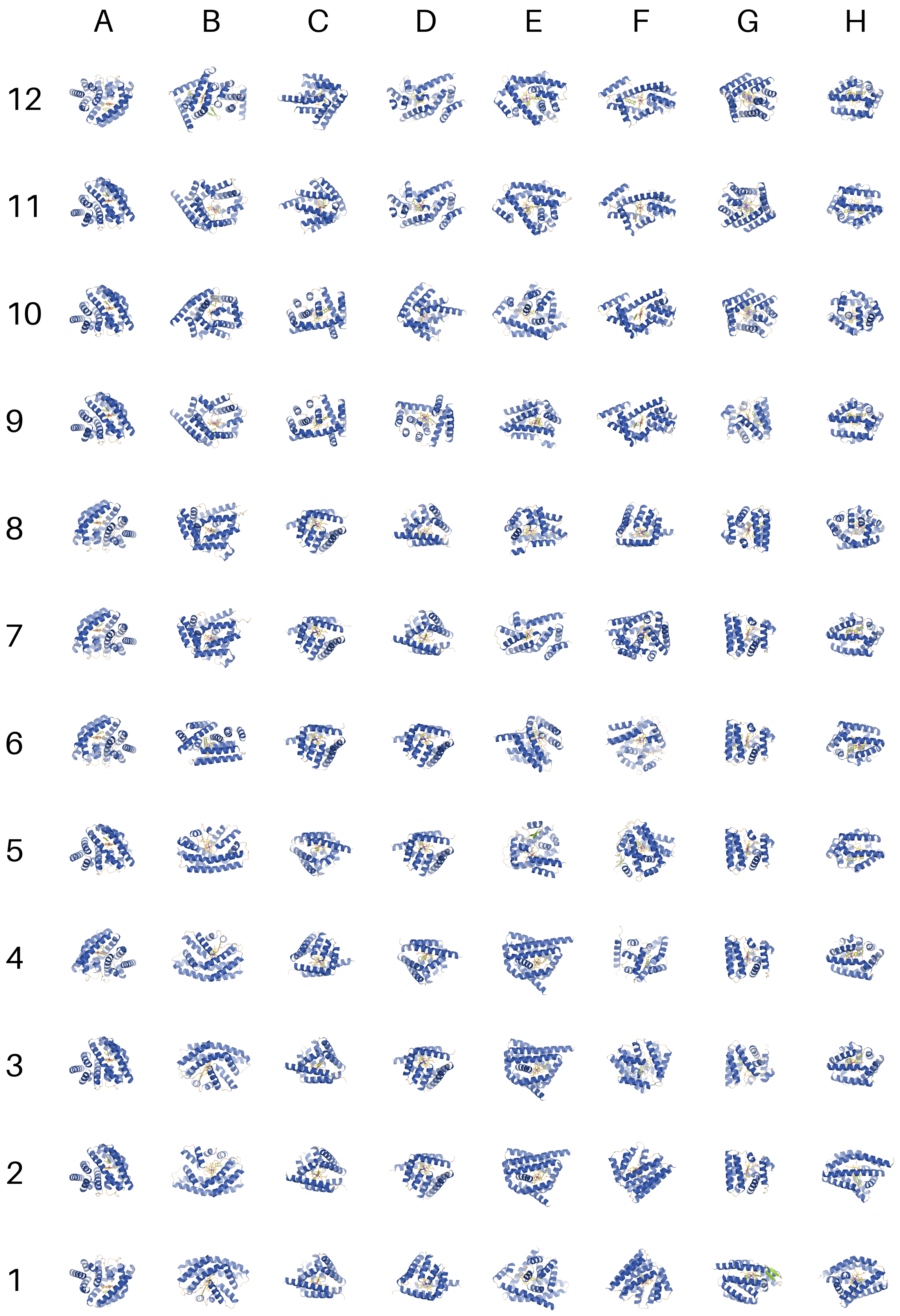


**Figure S15.** Designed **BR** hydroxylases selected for experimental testing (set A1-H12).

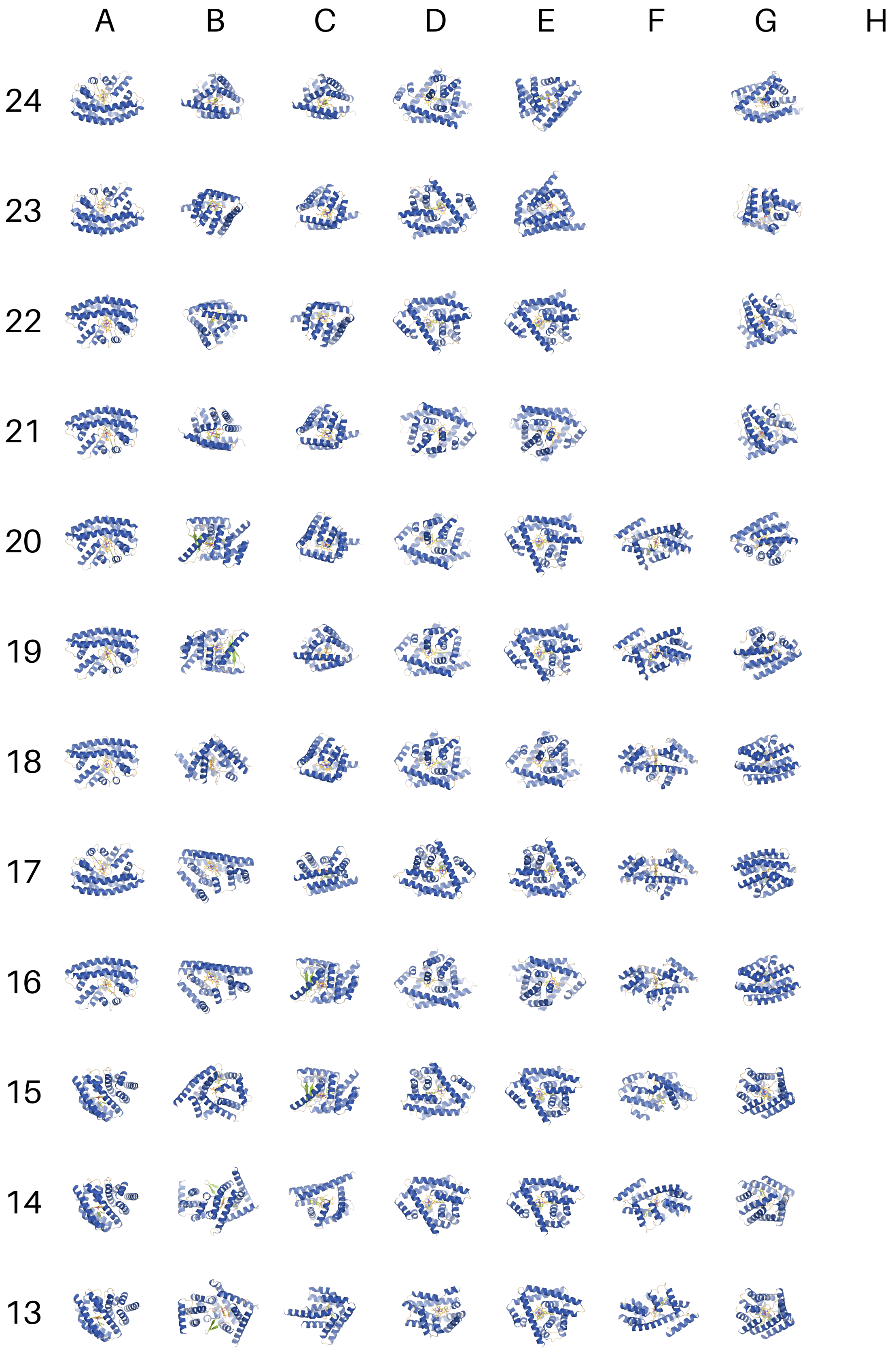


**Figure S16.** Designed **BR** hydroxylases selected for experimental testing (set A13-H24).

##### 3.1.2. In silico optimization of dOHbr-2

We used AlphaFold3 predictions to identify viable mutations in the **dOHbr-2** backbone for experimental characterization. In our previous work we learned that AF3 predictions are effective at removing certain negative mutations from the set - proteins that did not fold according to AF3 were also not functional (*14*). We selected 42 positions surrounding the entire active site (Figure S17), excluding the catalytic residues Cys-31, Glu-143 and His-168. The existing amino acid was swapped for each of the 18 other amino acids (excluding cysteine), and the resulting 828 sequences were co-folded with heme and the substrate **BR** using AlphaFold3 (10 recycles, 5 diffusion samples, no MSA, no templating). The models were analyzed for how accurately and confidently the protein backbone and active site structures were predicted (Table S10). The selection cutoffs for AF3 metrics were slightly adjusted compared to the design campaign, tightening the protein accuracy and confidence selection, but loosening the substrate accuracy (only looking at the lowest **BR** RMSD among five models, instead of their average). Surprisingly, >95% of the mutants folded correctly, with only the substrate RMSD being a more stringent filter and keeping 60% of the variants for a total of 432 passing sequences. To expand the selection at positions with fewer than five passing mutants, we applied loosened filtering criteria there (slightly higher substrate RMSD and lower heme and substrate confidences) and kept additional 25 variants for a total of 455 sequences. To reduce the set of mutations down to 382 (so that it could fit onto a single 384-well plate) we manually inspected the variants and excluded those that seemed redundant at more remote positions (Table S18, shown in orange).

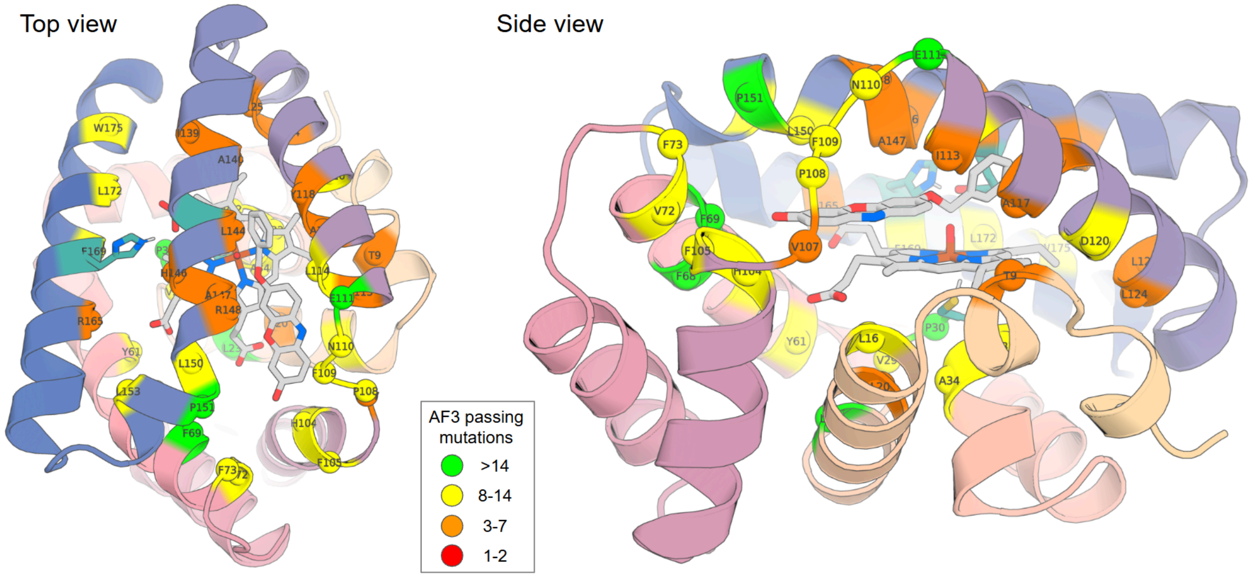


**Figure S17.** Positions selected for *in silico* mutagenesis of **dOHbr-2** are highlighted as colored spheres. Sphere color corresponds to how many mutants passed the AF3 selection.

**Table S10.** AlphaFold3 *holo* prediction selection criteria for *in silico* SSM of **dOHbr-2**. Metrics that differ from the cutoff values used in the design campaign (Table S8) are shown in bold.

| **Metric** | **Cutoff** | **Description** |
|---|---|---|
| chain_ptm_ch0 | >= 0.85 | Global structural accuracy of protein chain A (predicted TM score), average of 5 models |
| chain_iptm_ch0 | >= 0.80 | Interface structural accuracy of protein chain A, average of 5 models |
| chain_ptm_ch1 | >= 0.75 | Global structural accuracy of heme (predicted TM score), average of 5 models |
| chain_iptm_ch1 | >= 0.70 | Interface structural accuracy of heme, average of 5 models |
| chain_iptm_ch2 | >= 0.70 | Interface structural accuracy of substrate, average of 5 models |
| chain_pair_pae_min_inter_max | **<= 2.5** | Largest average interchain minPAE value (average of off-diagonal elements in PAE matrix) from 5 models. |
| rmsd | **<= 1.0 Å** | Cα RMSD between the design model and the prediction, average of 5 models |
| plddt | **>= 90.0** | All-atom local structural accuracy of protein chains. |
| plddt_BB | **>= 92.0** | Local structural accuracy of protein chains, averaged over backbone atoms (C,N,CA,O). |
| lig0_rmsd | <= 2.0 Å | Heme RMSD, average of 5 models |
| lig0_rmsd_max | <= 2.5 Å | Worst heme RMSD among 5 models. |
| lig0_plddt | >= 80.0 | Heme pLDDT, average of 5 models |
| **lig1_rmsd_min** | **<= 1.6 Å** | Substrate RMSD, lowest among 5 models |
| lig1_plddt | >= 70.0 | Substrate pLDDT, average of 5 models |
| rmsd_SR1 | **<= 0.8 Å** | All-atom RMSD of CYS, average of 5 models |
| rmsd_SR1_max | **<= 1.0 Å** | Worst CYS RMSD among 5 models. |
| plddt_SR1 | >= 85.0 | All-atom confidence of CYS, average of 5 models |
| rmsd_SR3 | **<= 1.2 Å** | All-atom RMSD of GLU, average of 5 models |
| rmsd_SR3_max | **<= 1.5 Å** | Worst GLU RMSD among 5 models. |
| plddt_SR3 | >= 85.0 | All-atom confidence of GLU, average of 5 models |
| rmsd_SR4 | **<= 1.4 Å** | All-atom RMSD of HIS, average of 5 models |
| rmsd_SR4_max | **<= 1.6 Å** | Worst HIS RMSD among 5 models. |
| plddt_SR4 | >= 85.0 | All-atom confidence of HIS, average of 5 models |

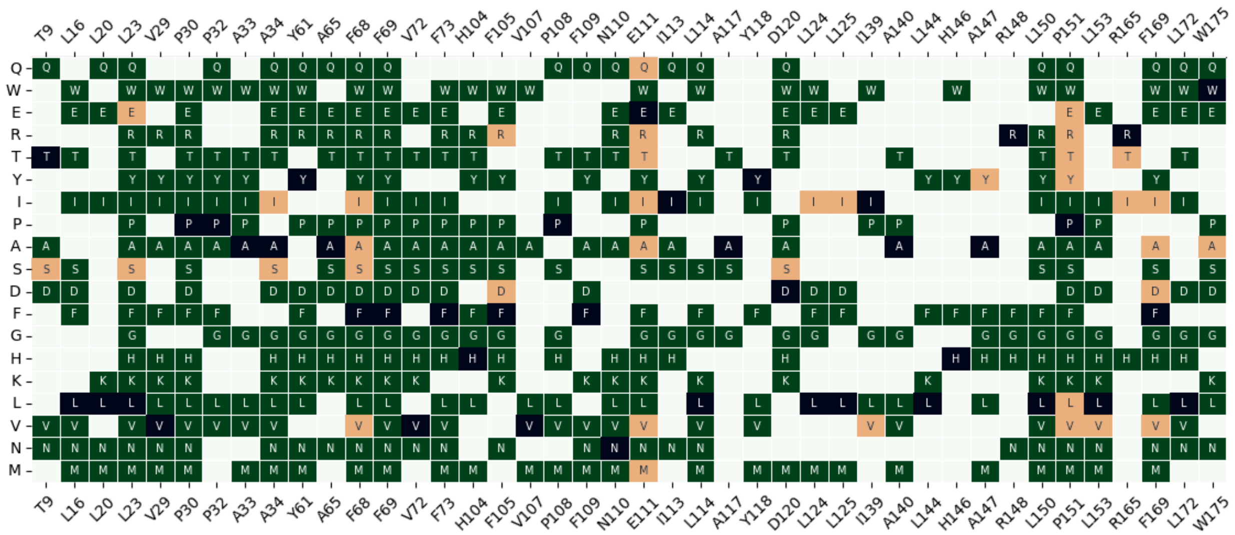

**Figure S18.** Map of mutations that passed AlphaFold3 selection at each of the 42 positions (dark green & orange). Parent amino acid is marked with a black square. Variants that passed, but were omitted from experimental testing are shown in orange.

#### 3.2. Experimental methods

##### 3.2.1. Screening and characterization of resorufin ether *O*-dealkylases

**Design library cloning and small-scale expression**

The synthetic DNA fragments encoding the designed protein sequences were obtained from Twist as Gene Fragments (RFD2 designs A1-F23) or were generated from oligonucleotide pools using the IPDassembly process (*15*) (RFD3 designs G1-H12). Linear DNA fragments encoding the designs were cloned using Golden Gate Assembly (GGA) (*16*) into FP0017 vector containing an N-terminal hexahistidine tag and a TEV cleavage site (designs A1-F23); or into LM627 vector (AddGene 191551) containing C-terminal SNAC and hexahistidine tags (designs G1-H12). The cloned plasmids were transformed into BL21(DE3) *E. coli* chemically competent cells (Intact Genomics) by incubating a mixture of 5 µL of cells and 1 µL of GGA mixture on ice for 30 minutes, followed by 30 second heat shock at 42 °C and a 2 minute rest on ice. The transformed cells were gently mixed with 100 µL of SOC outgrowth media and incubated at 37 °C for 1 hour. For selective cell growth, the outgrowth cultures were added to deep-well round-bottom 96w plates containing 900 µL of LB media, supplemented with kanamycin (50 µg/mL), and incubated at 37 °C with 1200 r.p.m. shaking for 16 hours. Thereafter, 100 µL of the starter culture was saved as a glycerol stock. For protein expression, 100 µL of the LB starter culture was used to inoculate in 900 µL of TB-II media, supplemented with kanamycin (50 µg/mL), and incubated for 4 hours at 37 °C while shaking at 1200 r.p.m. Expression was induced by the addition of 1 mM of IPTG and the cultures incubated for 2 hours at 37 °C while shaking at 1200 r.p.m. The cells were collected by centrifugation at 4150g for 8 minutes, with the discarded media removed. At this point many cell pellets had a slight pink color, indicating the presence of heme-containing proteins. The cell pellets were stored at -80 °C until needed.

For cell lysis, the frozen cell pellets were thawed at room temperature and resuspended in 250 µL of KPi buffer (50 mM potassium phosphates, 200 mM NaCl, pH 7.2) containing 1.0 mg/mL lysozyme, 0.01 mg/mL deoxyribonuclease I (DNase I), 0.5 mg/mL Polymyxin B sulfate and 1 mM PMSF. The resuspended cell lysate was incubated for 10 minutes at 37 °C with shaking at 1200 r.p.m., after which 10 µM of hemin was added (as a 200 µM solution in DMSO), and the mixture shaken at 37 °C for another 45 minutes. The plates were centrifuged at 4100g for 30 min and supernatant was collected by careful pipetting.

It should be noted that at this stage the expressed proteins are polyclonal, meaning that they may contain multiple variants resulting from mistakes in DNA synthesis. However in our experience for the vast majority of designs the largest population is still the intended designed protein.

**BROD cell lysate assay**

The resorufin benzyl ether *O*-dealkylase (BROD) assays for initial discovery were performed by mixing 50 µM **BR** (in 500 µM DMSO stock), KPi buffer and clarified cell lysates containing the designed proteins (20% of the final volume). This mixture was transferred to a black flat-bottom half area 96-well plate containing a solution of hydrogen peroxide (final conc. 500 µM), and the fluorescence signal from the release of resorufin ($\lambda_{ex}/\lambda_{em}$ 554/593 nm) was collected using a Synergy Neo 2 platereader over the period of 1 hour. Figure S19 presents the normalized final fluorescence signal levels.

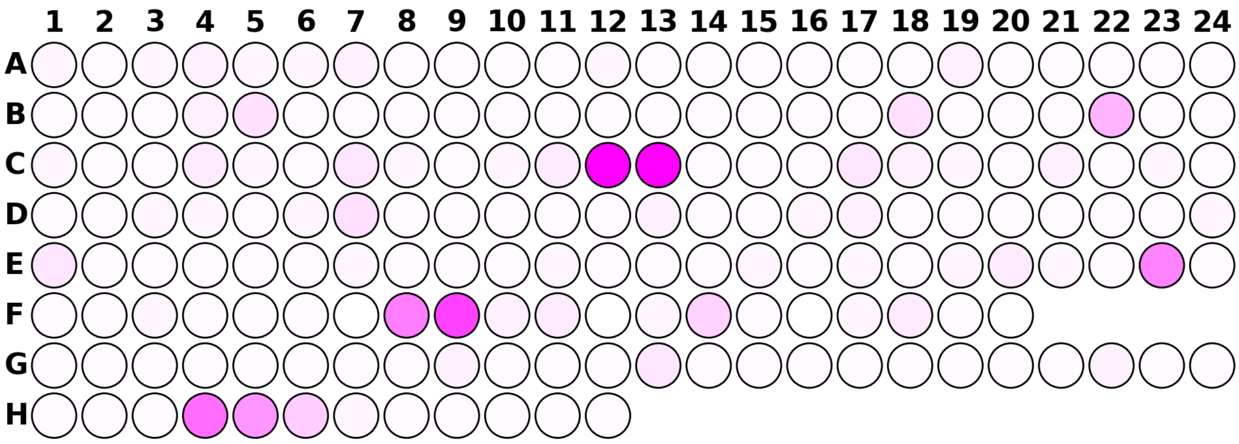


**Figure S19.** Enzymatic activity screening results in clarified *E. coli* cell lysate. The platemap is colored based on normalized fluorescence levels after 1 hour.

**Validation of design hits**

Production of clonal variants at larger scale

Designs with the strongest final fluorescence signal (C12, C13, F9, H4, H5) were validated as clonal purified proteins. For this we diluted a streak from the previously saved glycerol stock in $ddH_2O$, and spread it onto kanamycin-supplemented LB-agar plate. Individual colonies were picked on the next day, diluted in 10 µL of $ddH_2O$ - half of which was used to start 3 mL LB-kanamycin overnight cultures (in 14 mL round-bottom tube, shaken at 250 r.p.m at 37 °C), and half was used for sequencing. The DNA of the encoding region was amplified using colonyPCR with T7 and T7-term primers. The unpurified PCR products were sent to GeneWiz/Azenta for Sanger sequencing. We proceeded with the clones where the DNA sequence matched the sequence of the original DNA fragment. New clonal glycerol stocks were prepared from the overnight cultures, 2 mL of the culture was centrifuged and the cell pellet saved for DNA MiniPrep, and 900 µL was used to inoculate 40 mL TB-II autoinduction media (containing 50 µg/mL kanamycin, 50x 5052, 1 mM $MgSO_4$ and 1000x Studier trace metals). The cultures were grown and proteins expressed by shaking (250 r.p.m) at 37 °C for 16 hours. The cells were collected by centrifugation at 4100g for 8 minutes and frozen at -20 °C until needed. The cells were thawed and resuspended in 30 mL of lysis buffer (50 mM potassium phosphates, 200 mM NaCl, 25 mM imidazole, 0.1 mg/mL lysozyme, 10 µg/mL DNAse I, pH 7.4) and stored on ice. Hemin (0.5 mL of 6 mM hemin in 0.5 M NaOH) was added to the resuspended cells and vigorously shaken. The cells were lysed by ultrasonication (0.75" probe, 10 s on 15 s off, total 2.5 min on time) while cooled in an ice bucket. Cell debris was removed by centrifugation (30 minutes at 15,000 g, 12 °C). The clarified lysates were mixed with Ni-NTA resin, equilibrated for 15 minutes by rocking at 4 °C, and the lysate removed by gravity filtering. The protein-loaded resin was washed with 45 mL wash buffer (50 mM KPi, 200 mM NaCl, 25 mM imidazole, pH 7.4) and the bound protein was eluted off the resin with 1.3 mL of elution buffer (50 mM KPi, 200 mM NaCl, 300 mM imidazole, pH 8). The eluted proteins were filtered through a 0.22 µm filter, centrifuged at 18,000g for 5 minutes, and purified using size exclusion chromatography (AKTA, Superdex Increase 75 10/300 GL column) with KPi (50 mM Tris, 200 mM NaCl, pH 7.2) running buffer. The fractions corresponding to monomeric or lowest oligomeric fractions were collected. The chromatograms shown in Figure S20 were collected by monitoring the absorbance at 280 nm (protein) and 408 nm (heme).

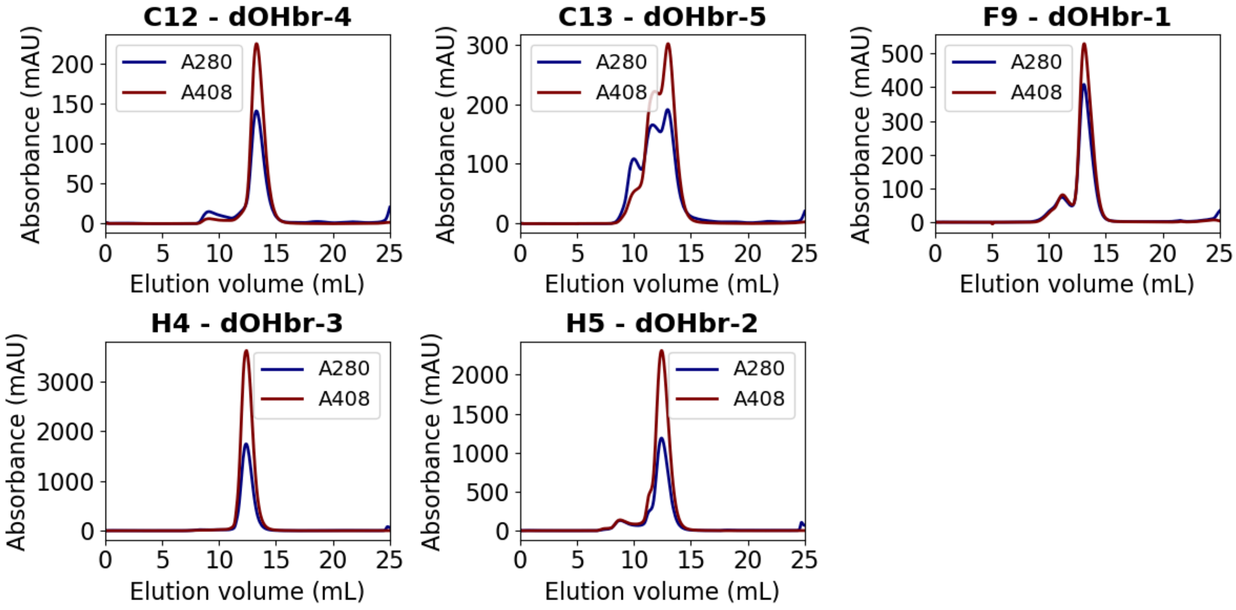


**Figure S20.** Size-exclusion chromatograms of purified heme-binding proteins, using Superdex Increase 75 10/300 GL column and 50 mM KPi 200 mM NaCl pH 7.2 running buffer. Blue trace corresponds to a chromatogram based on absorption at 280 nm, red trace is collected at 408 nm and is indicative of heme-containing species.

BROD activity under normalized conditions

The resorufin benzyl ether O-dealkylase (BROD) assays for hit validation were performed at two $H_2O_2$ concentrations by first mixing 50 µM **BR** (in 500 µM DMSO stock), KPi buffer and 2 µM purified hemoprotein. This mixture was transferred to a black flat-bottom half area 96-well plate containing solutions of hydrogen peroxide (final conc. 500 µM and 4 mM), and the fluorescence signal from the release of resorufin ($\lambda_{ex}/\lambda_{em}$ 554/593 nm) was collected using a Synergy Neo 2 platereader over the period of 1 hour. Figure S21 presents the collected reaction progress curves.

Varying the $H_2O_2$ concentration revealed that two of the designs (**C12** and **C13**) have increased turnover numbers (TON) at lower [$H_2O_2$], suggesting some level of peroxide sensitivity. Designs **F9**, **H4** and **H5** on the other hand display both higher TON and activity as [$H_2O_2$] increases. **H5** displays a decay of resorufin after 10 minutes, likely because of bleaching through peroxidase activity. This is mitigated by reducing [$H_2O_2$] or [enzyme]. These results confirm that the lysate screening hits are indeed functional enzymes under controlled conditions.

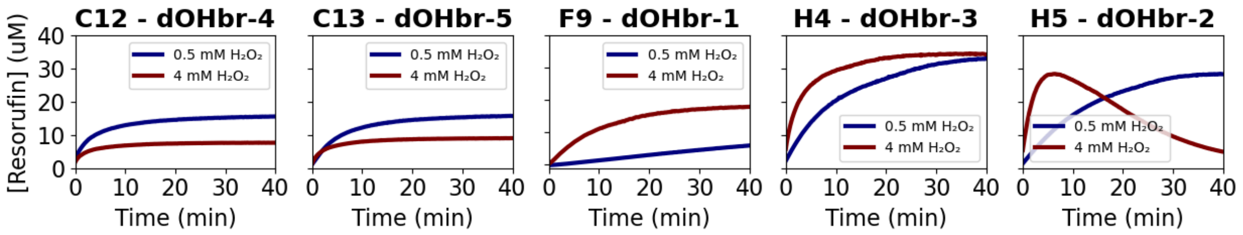


**Figure S21.** Resorufin benzyl ether assay performed with purified enzymes (2 μM), 50 μM **BR** and 0.5 or 4 mM $H_2O_2$. Reaction progress was monitored by resorufin fluorescence ($\lambda_{ex}/\lambda_{em}$ 554/593 nm).

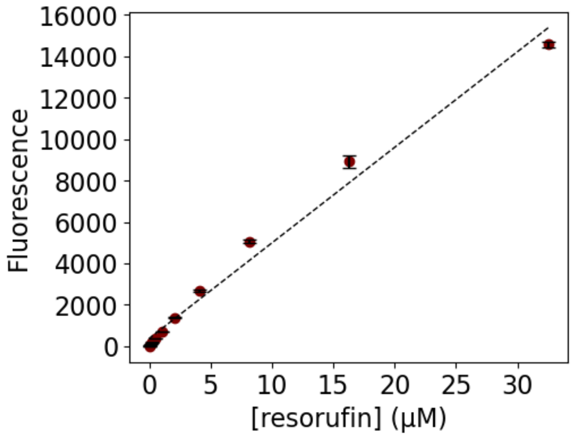


**Figure S22.** Calibration curve for resorufin fluorescence concentration dependence ($r^2$ = 0.985).

Detection of BROD reaction products

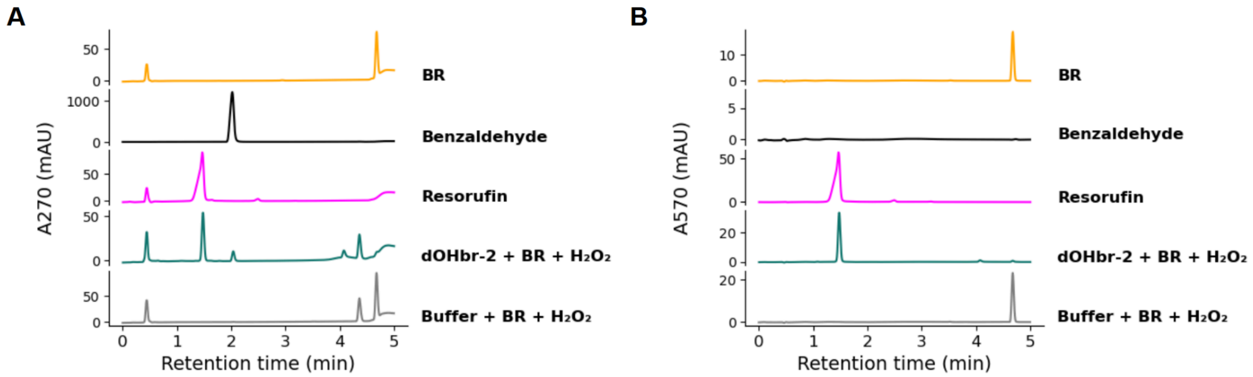


**Figure S23.** UPLC analysis of the BROD reaction, catalyzed by **dOHbr-2**, confirming the enzymatic reaction product formation. The enzymatic reaction was performed with 2 μM enzyme, 50 μM **BR**, and 4 mM $H_2O_2$. After 1 hour, the enzymatic reaction was quenched by the addition of equal volume of acetonitrile. 5 μL of each sample was injected for UPLC analysis. (A) Chromatogram collected based on absorption at 270 nm. (B) Chromatogram collected based on absorption at 570 nm.

##### 3.2.2. Optimization of dOHbr-1

Library construction

In Round 1 of evolution, 20 positions were individually randomized using degenerate NNK codons. DNA libraries were constructed by overlap extension polymerase chain reaction (PCR) with end primers designed for Golden Gate assembly. Constructs were subcloned into an in-house made pET28-ccdB vector using BsaI restriction enzyme.

Shuffling by overlap extension PCR

After evolution, beneficial diversity was combined by DNA shuffling of fragments generated by overlap extension PCR. Primers were designed that encoded either the parent amino acid or the identified mutation. These primers were used to generate short fragments that were gel-purified and mixed for assembly of the full-length gene by overlap extension PCR. Final full-length genes contain all possible combinations of mutations at specified positions. Genes were cloned as described above.

Library Screening

For protein expression and screening, all transfer and aliquoting steps were performed using a Hamilton liquid-handling robot. Chemically competent *E. coli* 5α cells were transformed with the appropriate library plasmids. Freshly transformed colonies were used to inoculate 2YT medium (150 µL) supplemented with kanamycin sulfate (50 µg/mL) in Corning Costar 96-well microtitre round-bottom plates. Each plate also contained six freshly transformed clones of the parent template and two clones of pBbE8k_RFP as an internal reference. Plates were incubated overnight (30 °C, 80% humidity, 850 r.p.m.), then an aliquot of overnight culture (20 µL) was used to inoculate 2YT medium (480 µL) supplemented with kanamycin sulfate (50 µg/mL). The cultures were incubated (30 °C, 80% humidity, 850 r.p.m.) until an $OD_{600}$ of 0.6 was reached, temperature reduced to 25 °C and IPTG added (100 µM final concentration) and grown for a further 20 hours. Cells were collected by centrifugation (2,900 × g for 10 min). The supernatant was discarded, and the pelleted cells were resuspended in 400 µL lysis buffer (50 mM potassium phosphate, 200 mM NaCl pH 7.2) supplemented with lysozyme (1.0 mg/mL), polymyxin B (0.5 mg/mL) and DNase I (10 µg/mL)) and incubated (1 h, 30 °C, 80% humidity) with shaking (850 r.p.m.), followed by a 60 °C heat shock for 1 hr (at 850 r.p.m.). Precipitates were removed by centrifugation at 2,900 × g for 20 mins. 30 µL of clarified lysate was transferred to a 96-well microtitre plate followed by the addition of 30 µL hemin (final assay concentration of 1 µM) and incubated at room temperature for 20 mins. Subsequently, the addition of buffer (50 mM potassium phosphate, 200 mM NaCl pH 7.2) and **BR** (final concentration 25 µM) in DMSO (10% v/v final concentration). Reactions were initiated by the addition of $H_2O_2$ (final concentration 4 mM, from a 30 mM stock). Resorufin formation was monitored by fluorescence excitation at 571 nm over 40 mins using a CLARIOstar plate reader (BMG Labtech).

The most active clones were rescreened in lysate in triplicate. Expression and screening were performed as described above, with cultures inoculated from glycerol stocks prepared from the original library cultures. The most active variants were rescreened as purified proteins with starter cultures inoculated from glycerol stocks prepared from the original library plate overnight cultures.

Protein expression and purification

For expression of **dOHbr-1** and variants, single colonies of freshly transformed chemically competent *E. coli* BL21 (DE3) cells were cultured (18 hr at 37 °C, 180 r.p.m.) in 5 mL 2YT supplemented with kanamycin (50 µg/mL). These starter cultures were used to inoculate 2YT medium (50 mL) containing kanamycin (50 µg/mL). Cultures were grown at 37 °C (180 r.p.m.) until $OD_{600}$ of 0.8. Temperature reduced to 25 °C and protein expression was initiated with 100 µM of IPTG, and grown for a further 18 hrs. Cells were collected by centrifugation (6,880 x g for 10 min) and resuspended in lysis buffer (50 mM potassium phosphate, 200 mM NaCl pH 7.2 supplemented with 20 mM imidazole, 10 µg/mL DNase I, 10 µg/mL lysozyme). Cells were disrupted by sonication (13 mm probe, 1 s on 1 s off – total duration 20 mins). The lysate was clarified by centrifugation at 10,000 × g for 20 mins. The supernatant was then incubated with hemin (final concentration 40 µM) for 30 mins on a rolling platform at 4 °C. The heme loaded clarified lysates were subjected to affinity chromatography using Ni-NTA Agarose (Qiagen). His-tagged proteins were eluted with a high imidazole buffer (50 mM potassium phosphate, 200 mM NaCl, 250 mM imidazole, pH 7.2). The eluent containing purified protein was buffer exchanged into assay buffer (50 mM potassium phosphate, 200 mM NaCl, pH 7.2) using a 10DG column (Bio-Rad). Subsequently, the designs were further purified by SEC ÄKTA pure system using a Superdex 75 16/600 column (GE Healthcare) preequilibrated with buffer (50 mM potassium phosphate, 200 mM NaCl pH 7.2). Proteins were aliquoted, flash frozen in liquid nitrogen and stored at -80 °C. The concentration of heme loaded proteins were determined using the extinction coefficient of the Soret peak (as determined using a pyridine hemochromogen assay (Table S12).

##### 3.2.3. Optimization of dOHbr-2

Fragmentation strategy and DNA library construction

The DNA strategy chosen for this mutagenesis campaign is based on assembling the mutant variants from three DNA fragments - two constant ones and one containing the mutation. Only the mutant-fragments would be individually synthesized. This reduces the cost of DNA synthesis and enables efficient combinatorial library testing. The sequence of **dOHbr-2** was split into three sections (Figure S24, positions 1-57, 58-128, 129-181) with the cut sites selected so that there would be at least two constant residues on either side.

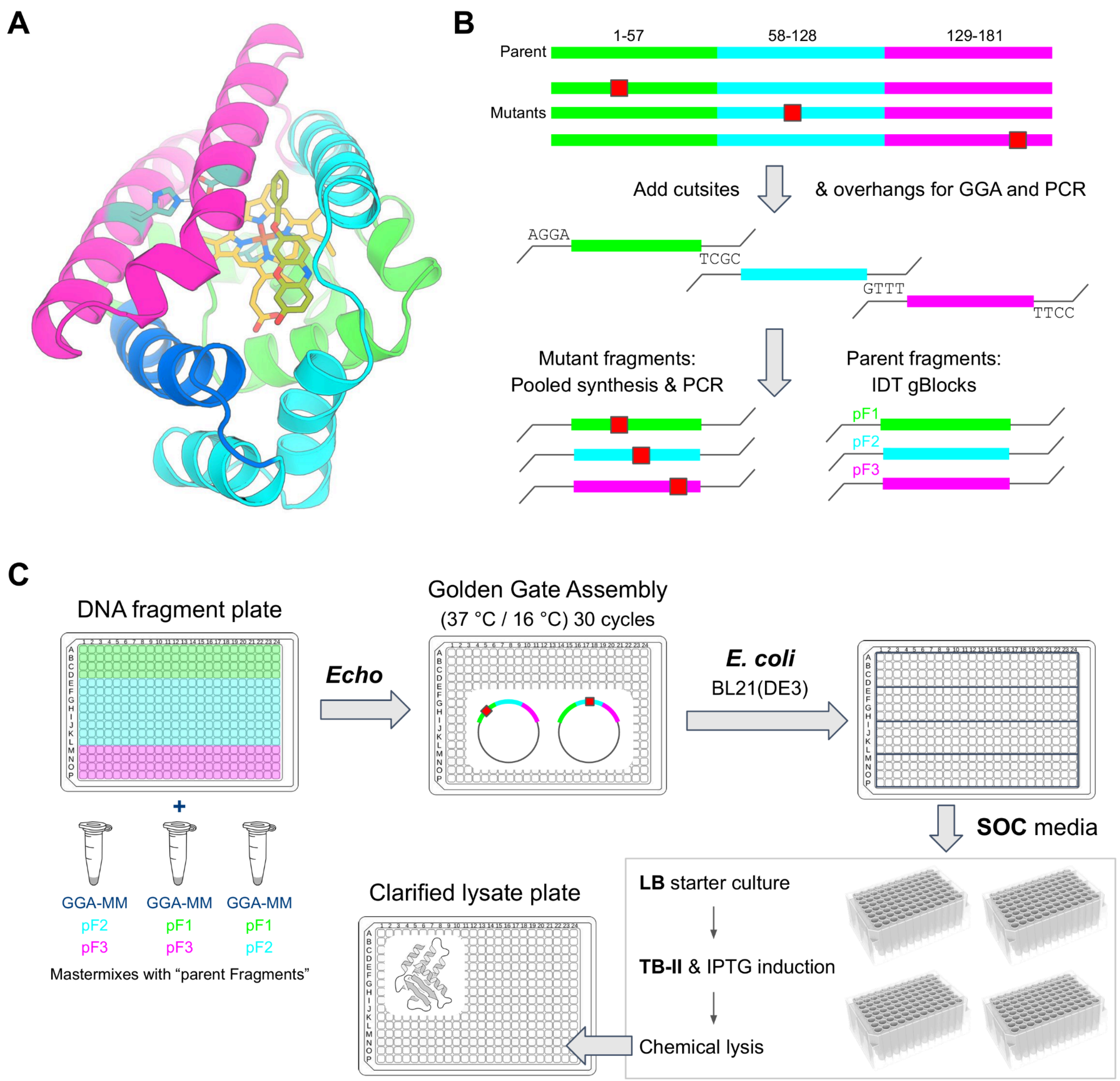


**Figure S24.** Fragmentation strategy and DNA library construction. **(A)** Fragmentation of **dOHbr-2** into three segments. Dark blue and cyan both belong to the second segment - the darker blue color distinguishes a helix with numerous mutated positions referenced in later sections. **(B)** Fragmentation of the encoding DNA for synthesis. **(C)** Assembly and protein expression workflow that produces a library of expressed proteins in 24 hours.

DNA sequences encoding the AF3-selected mutants of **dOHbr-2** (Figure S18) were created by swapping only the codon at a given position to a new one. The DNA sequences were split at the cutsites, and suitable 4 bp overhangs that would enable efficient Golden Gate assembly were selected within the 4 aa/12 bp cut region. BsaI restriction enzyme recognition sites were added to the 5' and 3' ends of the split fragments, along with orthogonal amplification primer sequences (*15*), and the fragments padded to 350 nucleotides. The synthetic DNA of mutant fragments was obtained using the IPDassembly process. These sequences were synthesized within a 350 nucleotide oligonucleotide pool at Twist Bioscience. Two nested PCR stages were conducted to dial-out design-specific fragments within each well of a microwell plate, yielding a 384-well plate where each well contains a unique mutant DNA fragment. The synthetic DNA fragments encoding the constant regions of **dOHbr-2** were obtained from IDT as gBlocks and PCR-amplified to produce more material for cloning reactions.

Mutant library production

The mutant fragments were assembled into full sequences in LM627 vector (with C-terminal SNAC and His tags) by Golden Gate assembly. Using the Echo acoustic liquid handling robot we combined in individual wells of a 384-well plate the amplified mutant fragments with the corresponding master mix solution (containing BsaI restriction enzyme, T4 DNA ligase, T4 buffer, LM627 cloning vector, and the two constant DNA fragments). The **dOHbr-2** parent design was assembled in two replicates as a control within the same experiment from the three individual constant fragments. The assembly reaction (1 µL total volume) was performed using 30 cycles of 1 minute at 37 °C and 1 minute at 16 °C, followed by 5 minute deactivation at 60 °C. The assembled plasmids were transformed into BL21(DE3) *E. coli* (Intact Genomics) by the addition of 5 µL of competent cells to the 384-well plate containing the cloning reaction mixtures. After incubation on ice for 30 minutes, heat shock at 42 °C for 30 second, and resting on ice for 2 minutes, the cells were transferred to deep round-bottom 96-well plates by the addition of 100 µL of SOC outgrowth media. The cultures were incubated at 37 °C for 1 hour, after which 900 µL of LB media containing kanamycin (50 µg/mL) was added and the cultures incubated at 37 °C for 16 hours while shaking at 1200 r.p.m. Thereafter, glycerol stocks were prepared from the starter cultures, and expression cultures were prepared by transferring 100 µL of the starter culture to 900 µL TB-II supplemented with kanamycin (50 µg/mL). The cultures were allowed to grow for 4 hours at 37 °C, shaking at 1200 r.p.m., after which 1 mM IPTG was added to induce protein expression, and the cultures shaken for another 2 hours at 37 °C. The cells were collected by centrifugation at 4100g for 8 minutes, the leftover media discarded, and the pellets frozen at -20 °C overnight.

For cell lysis, the frozen cell pellets were thawed at room temperature and resuspended in 250 µL of KPi buffer (50 mM potassium phosphates, 200 mM NaCl, pH 7.2) containing 1.0 mg/mL lysozyme, 0.01 mg/mL DNase I, 0.5 mg/mL Polymyxin B sulfate and 1 mM PMSF. The resuspended cell lysate was incubated for 10 minutes at 37 °C with shaking at 1200 r.p.m., after which 10 µM of hemin was added (as a 200 µM solution in DMSO), and the mixture shaken at 37 °C for another 45 minutes. The plates were centrifuged at 4100g for 30 min and supernatant was collected by careful pipetting. The clarified lysates were transferred to a 384-well plate and diluted to 2.5% v/v.

It should be noted that at this stage the expressed proteins are polyclonal, meaning that they may contain multiple variants resulting from mistakes in DNA synthesis. However in our experience for the vast majority of designs the largest population is still the intended designed protein.

Library screening with BROD assay

The resorufin benzyl ether *O*-dealkylase (BROD) assays were performed by first mixing 50 µM BR (in 500 µM DMSO stock), KPi buffer and clarified cell lysates containing the designed proteins (0.5% final lysate loading). This mixture was transferred to a black 384-well plate containing a solution of hydrogen peroxide (final conc. 4 mM, final reaction volume 15 µL), and the fluorescence signal from the release of resorufin ($\lambda_{ex}/\lambda_{em}$ 554/593 nm) was collected using a Synergy Neo 2 platereader over the period of 1 hour. The screening assay was performed in duplicate. The activity improvements were calculated as a combination of initial rate improvement and yield improvement using the equation $\texttt{score} = \texttt{rate}_{\texttt{Mut}}/\texttt{rate}_{\texttt{WT}} \times \texttt{Yield}_{\texttt{Mut}}/\texttt{Yield}_{\texttt{WT}}$, where `rate` is the slope of fluorescence increase in the first five minutes, and `Yield` is the highest fluorescence signal reached at 1 hour. We chose this combined evaluation approach to capture variants that display either significant rate increase, and/or turnover numbers, while penalizing those that show significant decrease in either. Figure S25 presents the activity improvements of each mutant. To validate the reproducibility of our approach we assembled and tested biological replicates of selected positive and negative hits from the initial screen and found the results to be overall consistent (Figure S26).

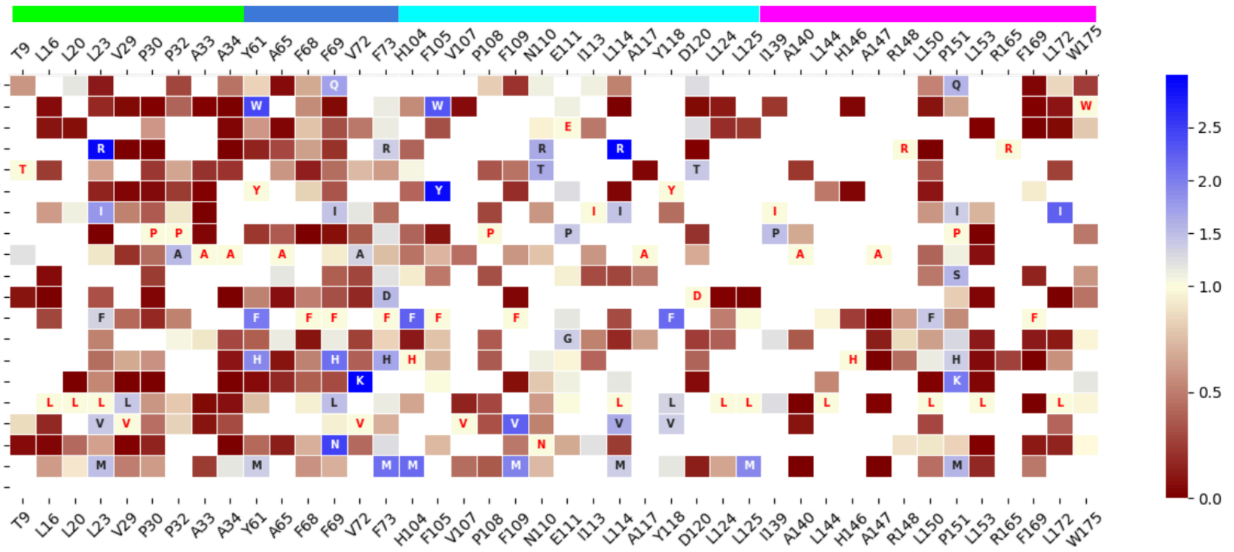


**Figure S25.** Activity improvements of **dOHbr-2** mutants of the parent design in BROD assay. Horizontal colored bar (green, blue, magenta) visualizes in which section of the protein (Figure S24A) a given position lies. For each position the identity of the parent residue is written red. Mutants with >1.3 fold improvement over parent are displayed with gray or white text. Gray to blue cell colors indicate improvement in activity, and orange to red colors indicate decreased activity.

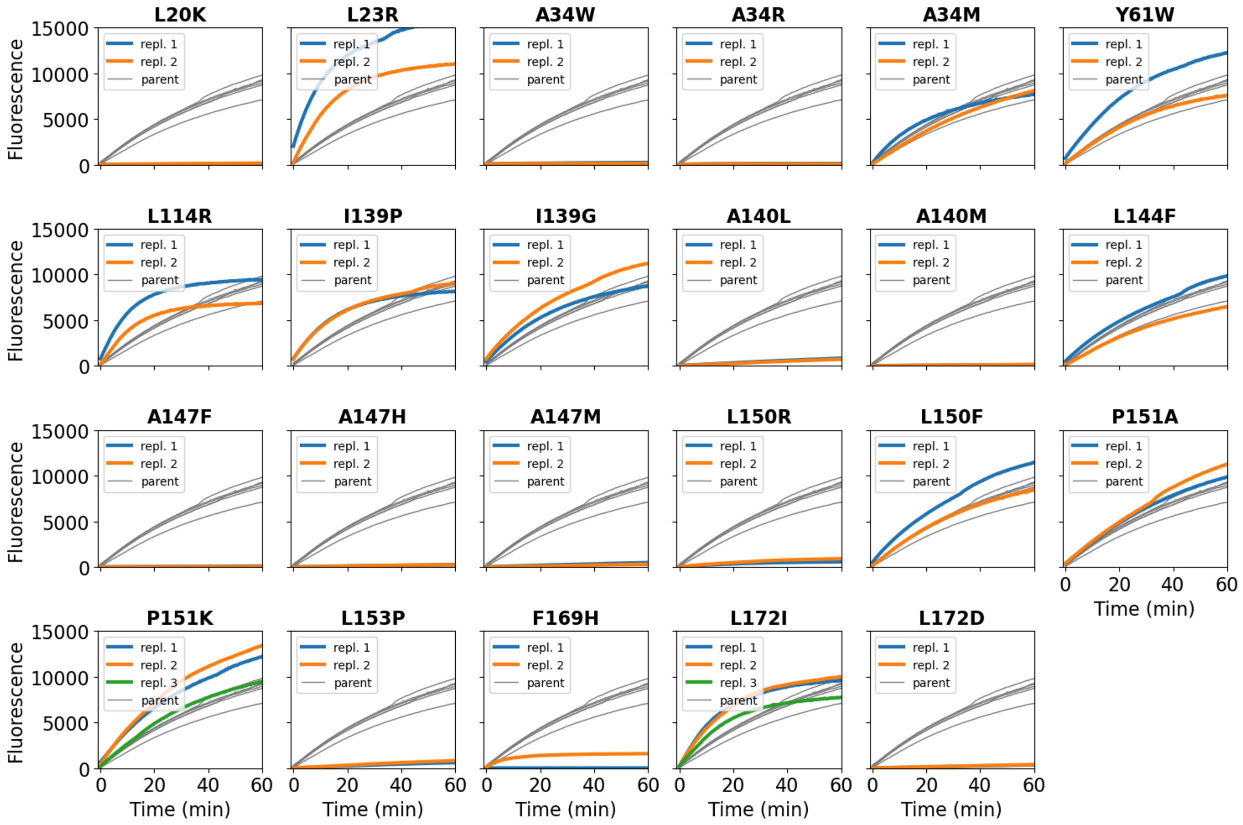


**Figure S26.** BROD assays performed with biological replicates (colored curves) of selected mutants of **dOHbr-2**. Gray curves belong to six biological replicates of **dOHbr-2**. Reactions were performed with 0.5% cell lysate, 50 µM **BR**, and 4 mM $H_2O_2$.

To confirm that the observed activity in the polyclonal mixtures corresponds to the intended mutant, a few selected variants were isolated as single clones by streaking the glycerol stock on LB-agar plate, isolating and sequencing the colonies and expressing them on 40 mL scale, followed by SEC purification as heme-loaded proteins. BROD assay was performed with 50 µM **BR**, 100 nM enzyme, in KPi buffer with 25% DMSO as co-solvent. The results shown in Figure S27 indicate that the positive hits from the polyclonal lysate screen are in fact more active than the parent.

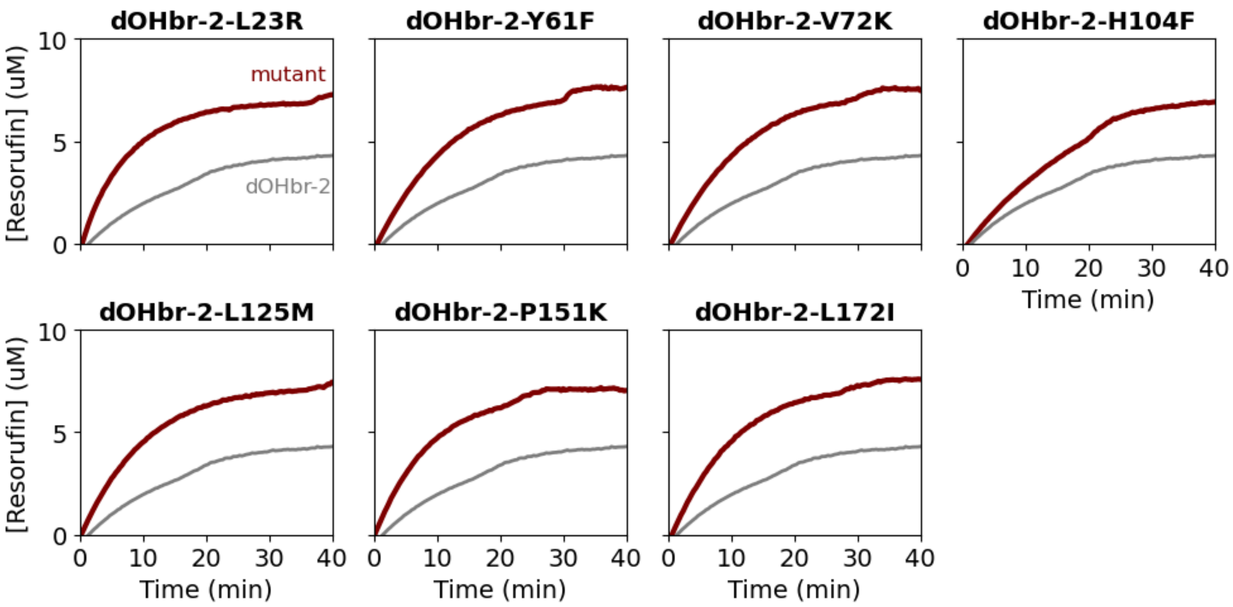


**Figure S27.** BROD assay validation of selected point mutants (with purified clonal proteins) from the mutagenesis of **dOHbr-2**. Red progress curves correspond to the mutants, and the gray curve represents the parent **dOHbr-2**. BROD assay was performed with 50 µM **BR**, 100 nM enzyme, in KPi buffer with 25% DMSO as co-solvent.

<u>Combinatorial library construction</u>

Leveraging the DNA fragment strategy we used the existing physical inventory of mutant fragments to combine them into double-mutant variants and explore the combinatorial and potentially epistatic effects of previously identified beneficial mutations. The beneficial mutations were selected based on slightly different fold-increase cutoffs for the three protein segments: 1.2-fold for segment 1 and 3, and 1.5-fold for segment 2. This was done to balance the library because the 2nd fragment pool was the largest, and also offered the most beneficial mutations, allowing a more stringent selection. Since the existing fragments only allow the testing of interfragment double-mutants, we also obtained an additional intrafragment double-mutant library of DNA fragments (using IPDassembly) that combine two mutations belonging to the same protein section. Taken together, we constructed a library of 723 double-mutants.

The DNA fragments were assembled using a similar approach as described for the first round of mutagenesis, using Echo liquid handling system to set up the Golden Gate Assembly reactions. The library was transformed into BL21(DE3) *E. coli* (Intact Genomics) and the proteins expressed and lysed using the same approach as described for the first round - 1 mL cultures, expressed using IPTG induction, 10 µM hemin added during lysis. The clarified lysates were diluted to 1% (v/v) and stored in 384-well plates.

<u>Combinatorial library screening</u>

The resorufin benzyl ether *O*-dealkylase (BROD) assays were performed by first mixing 50 µM BR (in 200 µM DMSO stock, 25% final DMSO), KPi buffer, and clarified cell lysates containing the designed proteins (0.2% final lysate loading). We reduced the lysate loading to increase the sensitivity of the assay in detecting improvements in activity. This mixture was transferred to a black 384-well plate containing a solution of hydrogen peroxide (final conc. 4 mM, final reaction volume 15 µL), and the fluorescence signal from the release of resorufin ($\lambda_{ex}/\lambda_{em}$ 554/593 nm) was collected using a Synergy Neo 2 platereader over the period of 1 hour. The screening assay was performed in triplicate.

The library contained 22 variants with 4-fold apparent activity improvement, and 7 variants with 5-fold improvement. This analysis revealed that certain mutations (Y61{F,W}, P151{K,M}, L125M, L150F, L172I) can be successfully combined with many other mutations, whereas some others (L20Q, L23{I,R}) seem to be more sensitive. Some mutations did not productively combine with any other mutation - T9A, L23M, V29H, P32A, A34M, N110R - the first five of these are situated underneath the heme. We selected 18 double mutants for validation and further characterization.

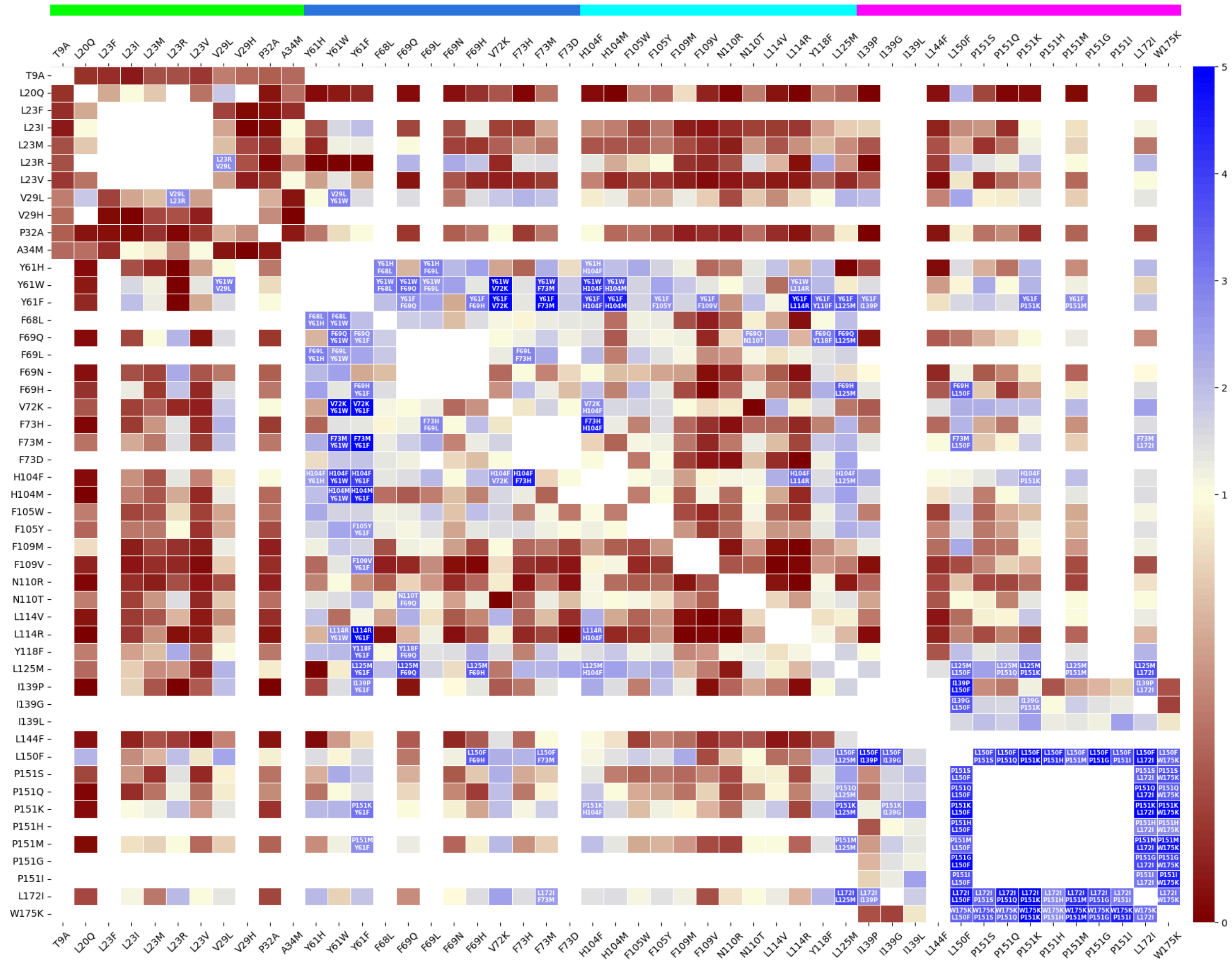


**Figure S28.** Combinatorial double-mutation landscape of **dOHbr-2** in the BROD reaction. Activity improvements are shown as a combination of initial rate and maximum fluorescence. Mutants with >2.5 fold improvement over parent are displayed with white text. Gray to blue cell colors indicate improvement in activity, and orange to red colors indicate decreased activity. Horizontal colored bar (green, blue, magenta) visualizes in which section of the protein (Figure S24A) a given position lies.

Validation and characterization of double-mutants

Clonal purified protein samples were prepared following the validation procedure described in section 3.2.1. BROD assay was performed with 50 μM **BR**, 4 mM $H_2O_2$, 100 nM enzyme, in KPi buffer with 25% DMSO as co-solvent (Figure S29). We found that most of the double-mutants were significantly more active than the parent, as well as more active than the previously characterized point mutants. The Y61W-V72K mutant displayed the highest activity, providing 150 TTN under these conditions. Position 61 is located underneath the heme carboxylates, and the Tyr forms a hydrogen bond with one of them, with Y61W and Y61F mutations potentially resolving the heme propionate conformational frustration that we observed in the crystal structure of **dOHbr-2**. Position 72 is located near the substrate entry, and the V72K mutation may create an additional hydrogen bond interaction with the phenolic oxygen of the resorufin moiety of the substrate. Similarly, P151K is located near the substrate entry and could offer an additional hydrogen bond to the substrate. Of other notable mutations, L23R is situated underneath the heme, and the introduced Arg may also reorient the heme propionates through hydrogen bonds. The roles of L125M and L172I are not immediately apparent - both are behind the heme and away from the substrate and could potentially subtly optimize the positioning of heme through hydrophobic interactions. A 3 Å resolution crystal structure of **dOHbr-2**-L125M-L150F revealed that the L150F mutation likely reduces the conformational flexibility of the frustrated heme propionate (Figure S36).

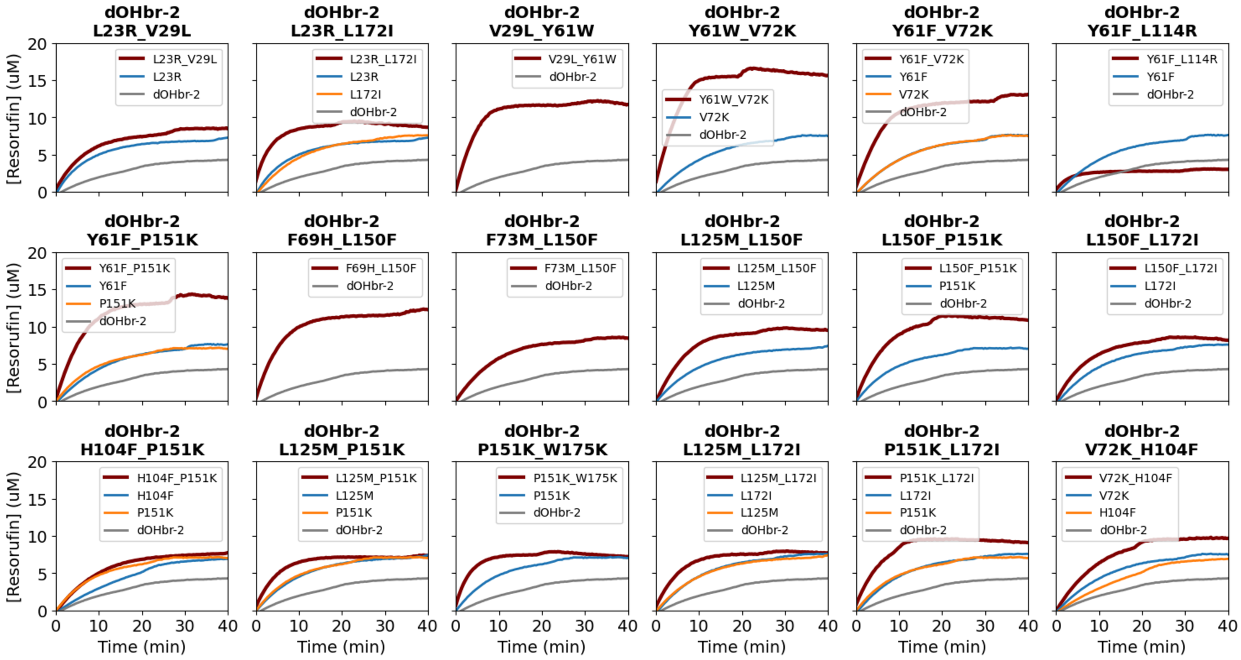


**Figure S29.** Time-courses of the BROD reaction, validating selected double mutants (with purified clonal proteins) from the combinatorial mutagenesis of **dOHbr-2**. Red progress curves correspond to the mutants, and the gray curve represents the parent **dOHbr-2**. Blue and orange progress curves correspond to the single mutations that make up a given variant (from Figure S27). BROD assay was performed with 50 μM **BR**, 4 mM $H_2O_2$, 100 nM enzyme, in KPi buffer with 25% DMSO as co-solvent.

##### 3.2.4. Kinetic characterization

Kinetic assays for **dOHbr-1**, **dOHbr-2**, and their variants were performed on a Cary UV-50 spectrophotometer (Varian) with a 1 cm pathlength cuvette. Steady-state kinetics assays to determine the total turnover numbers were carried out using **BR** (10 µM) and enzymes (concentrations of 50 nM and 5 nM, for **dOHbr-1** and **dOHbr-1-TLIA**, respectively). The reaction was initiated by the addition of 4 mM $H_2O_2$, and the UV−Vis spectra at 572 nm was recorded immediately as a function of time. The absorbance of the product was converted to concentration using the extinction coefficient ($\varepsilon_{572}$) of 56,000 $M^{-1}$ $cm^{-1}$. Assays were performed in triplicate.

Initial velocity ($v_0$) versus [BR] were measured using His-tagged purified **dOHbr-1** (500 nM), **dOHbr-1-TLIA** (50 nM) and **dOHbr-2** and its variants (50 nM). Reactions were performed in KPi buffer (40 mM potassium phosphate, 140 mM NaCl, pH 7.2) with 25% v/v DMSO co-solvent and varying concentrations of **BR** (1-25 µM). The reaction was initiated by the addition of 4 mM $H_2O_2$, and the UV−Vis spectra at 572 nm was recorded immediately as a function of time. The absorbance of the product was converted to concentration using the extinction coefficient ($\varepsilon_{572}$) of 56,000 $M^{-1}$ $cm^{-1}$. Assays were performed in triplicate. The observed initial rates were fitted to the Michaelis−Menten equation (Table S11, Figure S30).

Kinetic assays for **dOHbr-3**, **dOHbr-4**, and **dOHbr-5** were performed on BioTek Synergy Neo2 with fluorescence measurements by monitoring an excitation/emission wavelength of 554 nm and 593 nm, respectively. Initial velocity ($v_0$) versus [BR] were measured using His-tagged purified protein samples. The concentration of hemoprotein was determined using the pyridine hemochrome assay. 50 µl of enzymatic reactions were performed in KPi buffer (40 mM potassium phosphate, 140 mM NaCl, pH 7.2) with 25% v/v DMSO co-solvent and varying concentrations of **BR** (0.4-25 µM). The reaction solutions were prepared by premixing the purified protein sample and the serially diluted substrate with 25% v/v DMSO co-solvent, and adding them to a solution of 4 mM $H_2O_2$ in a 96-well half area plate to start the reaction. The concentrations of 10 nM, 100 nM, and 100 nM were used for **dOHbr-3, dOHbr-4,** and **dOHbr-5**, respectively. Half area 96-well flat bottom assay plates with a non-binding surface made from black polystyrene, were purchased from Corning. The fluorescence signal at 554/593 nm was recorded immediately as a function of time for 10 minutes at 25 °C. Assays were performed in triplicate. To determine the Michaelis−Menten parameters, we used the same data processing workflow as described for aromatic hydroxylase kinetics in section 2.2, using the calibration curve shown in Figure S22 to convert resorufin fluorescence into concentration. The obtained kinetic parameters and plots are shown in Table S11 and Figure S30.

**Table S11**. Michaelis-Menten kinetic parameters of peroxygenase reactions at saturating conditions of **BR** for designed proteins and their variants. Assay conditions: **dOHbr-1** (500 nM), **dOHbr-1-TLIA** (50 nM), **dOHbr-2** and its variants (50 nM), **dOHbr-3** (10 nM), **dOHbr-4** (100 nM), **dOHbr-5** (100 nM), 4 mM $H_2O_2$, varying **BR** concentrations (1 – 25 μM) with 25% v/v DMSO co-solvent. In 40 mM potassium phosphate, 140 mM NaCl pH 7.2.

| | $k_{cat}$ ($s^{-1}$) | $K_M$ (μM) | $k_{cat}/K_M$ ($s^{-1}$ $M^{-1}$) |
|---|---|---|---|
| **dOHbr-1** | 0.097 ± 0.0036 | 8.91± 0.807 | (1.09 ± 0.11) × $10^4$ |
| **dOHbr-1-TLIA** | 5.29 ± 0.19 | 6.43 ± 0.64 | (8.23 ± 0.87) × $10^5$ |
| **dOHbr-2** | 0.18 ± 0.01 | 1.16 ± 0.23 | (1.55 ± 0.32) × $10^5$ |
| **dOHbr-2**-V29L_Y61W | 1.64 ± 0.05 | 2.10 ± 0.29 | (7.82 ± 1.10) × $10^5$ |
| **dOHbr-2**-Y61W_V72K | 1.61 ± 0.08 | 1.66 ± 0.34 | (9.70 ± 2.04) × $10^5$ |
| **dOHbr-2**-L125M-L150F | 1.65 ± 0.06 | 3.62 ± 0.41 | (4.56 ± 0.54) × $10^5$ |
| **dOHbr-3** | 0.23 ± 0.01 | 1.83 ± 0.20 | (1.24 ± 0.14) × $10^5$ |
| **dOHbr-4** | 0.015 ± 0.001 | 0.95 ± 0.33 | (1.68 ± 0.60) × $10^4$ |
| **dOHbr-5** | 0.035 ± 0.001 | 2.68 ± 0.48 | (1.29 ± 0.24) × $10^4$ |

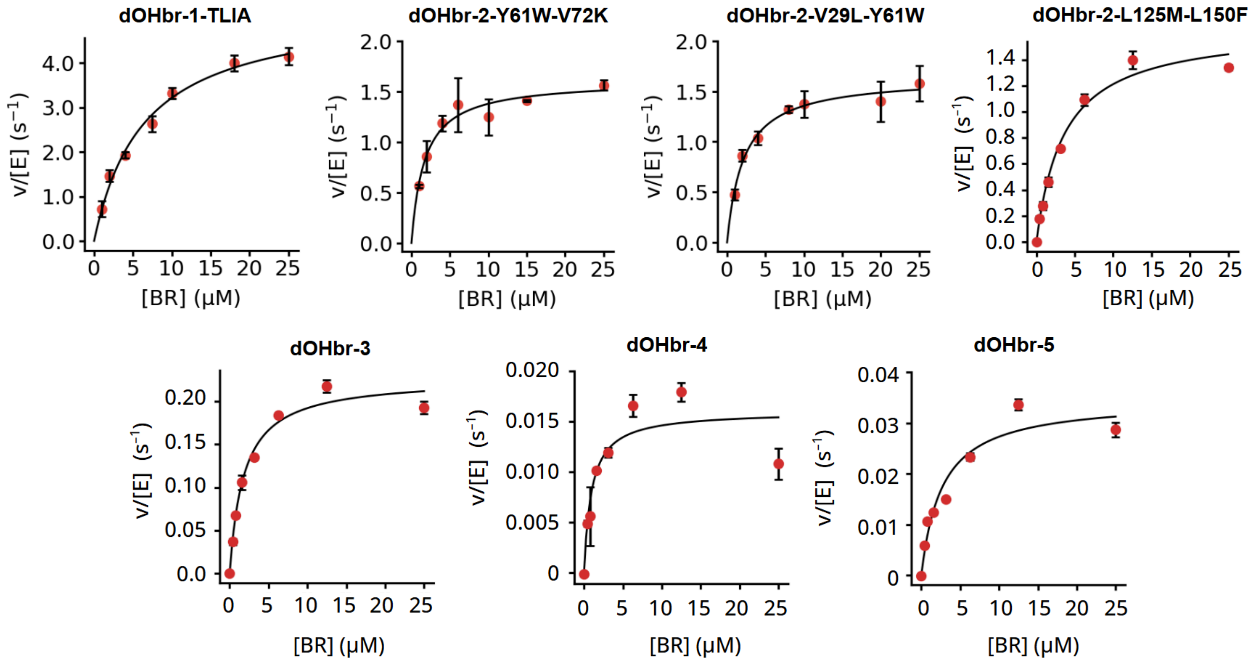


**Figure S30.** Michaelis-Menten kinetic analysis plots of **dOHbr-1-TLIA**, **dOHbr-2**-Y61W-V72K, **dOHbr-2**-V29L-Y61W, **dOHbr-2**-L125M-L150F, **dOHbr-3**, **dOHbr-4** and **dOHbr-5**.

##### 3.2.5. Spectroscopic characterization

We used temperature-dependent UV/Vis and circular dichroism spectroscopy to study the thermostability of the design hits and a few selected mutants of **dOHbr-2**. The spectra indicate that **dOHbr-2** and **dOHbr-3** are highly thermostable, with **dOHbr-1** being only slightly less stable at the highest measured temperature. The design **dOHbr-4** on the other hand is much less stable, showing changes in the secondary structure as well as in heme binding starting from 65 °C. The activity-increasing double-mutations of **dOHbr-2** have different effects on the protein thermostability. While V72K-H104F has fully retained the high stability of the parent scaffold, then most others start to show slightly lower stability at the highest temperatures. L23R-L172I on the other hand has become more destabilized and starts showing gradual changes already at 75 °C. This is possibly due to the introduction of the charged Arg-23 residue into the core of the protein.

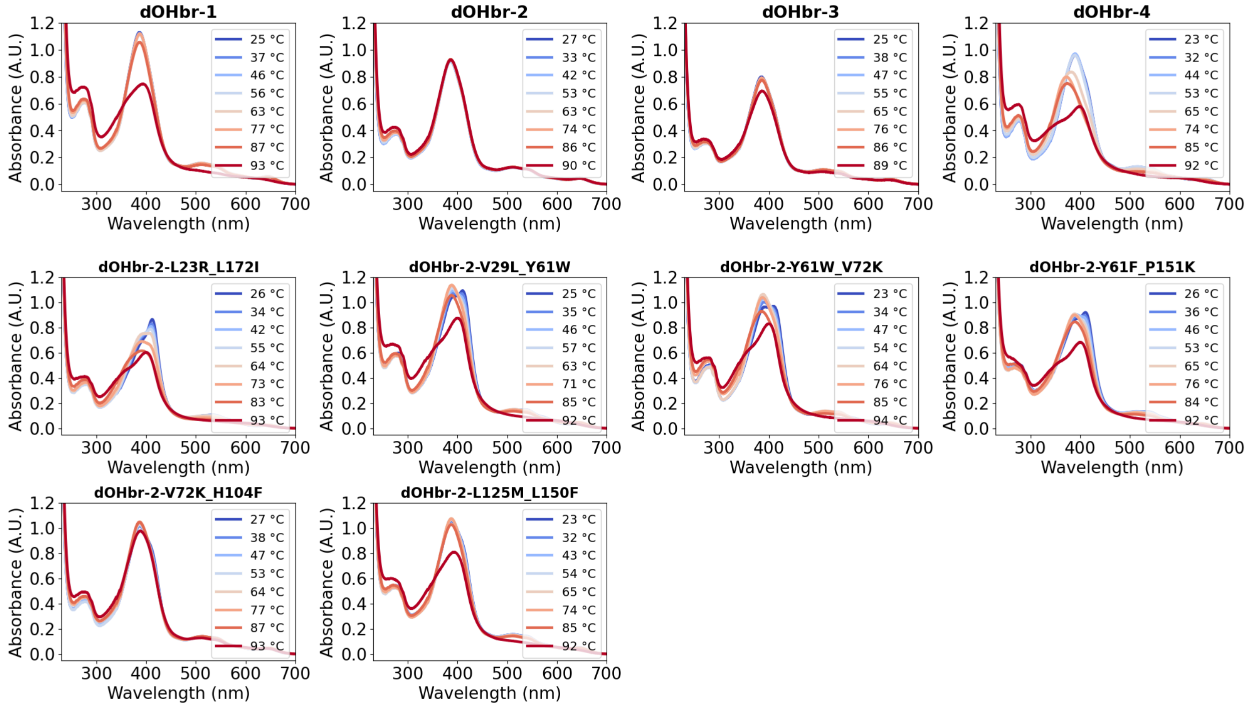


**Figure S31.** Temperature-dependent UV/Vis spectra of designed **BR** hydroxylases and **dOHbr-2** mutants. Spectra were measured in 50/200 KPi buffer at approximately 25 μM concentration.

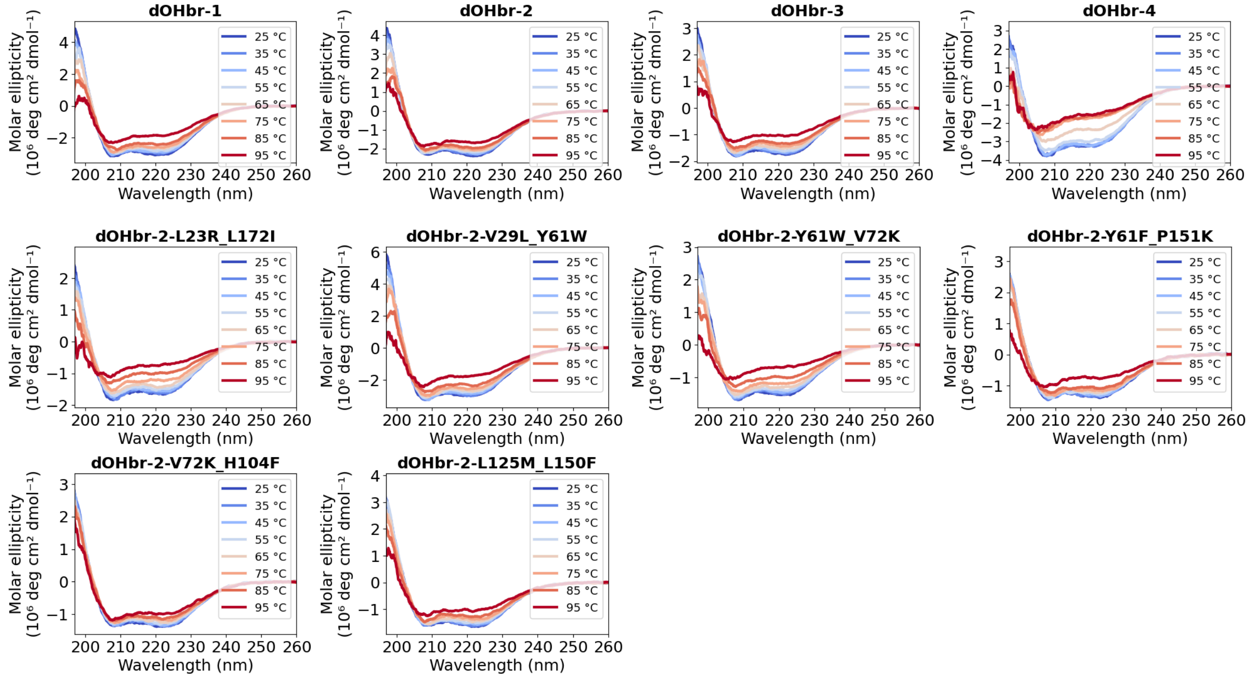


**Figure S32.** Temperature-dependent circular dichroism (CD) spectra of designed **BR** hydroxylases and **dOHbr-2** mutants. Spectra were measured in a 50/20 KPi buffer.

##### 3.3. Supplemental figures and tables

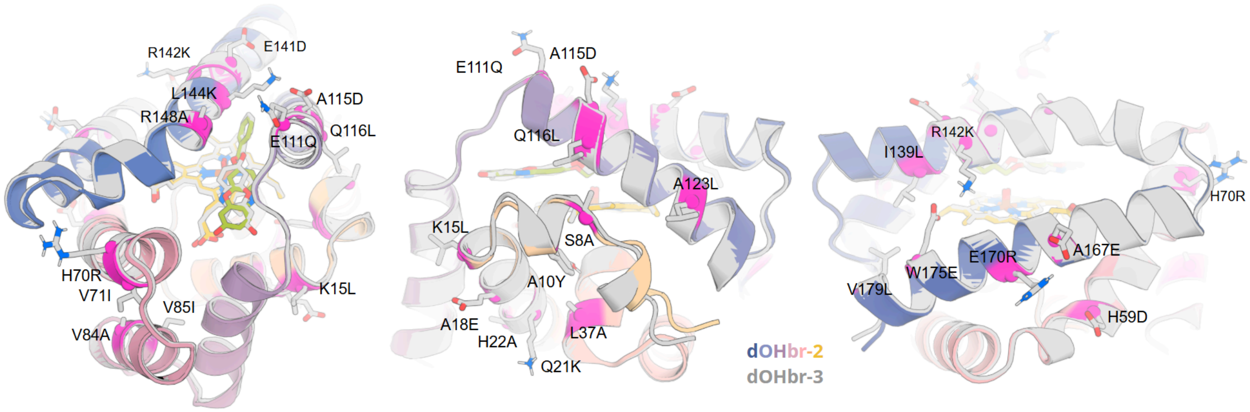


**Figure S33.** Visualization of **dOHbr-2** (colored) and **dOHbr-3** (gray) sequence differences at 25 positions. Residues in **dOHbr-3** as shown as gray sticks. The active site sequences are identical, with only the R148A, L144K and E111Q substitutions being somewhat close to the substrate. Rest of the mutations are all on the surface, introduced by the second sphere ligandMPNN sequence redesign in stage #4 of the design workflow.

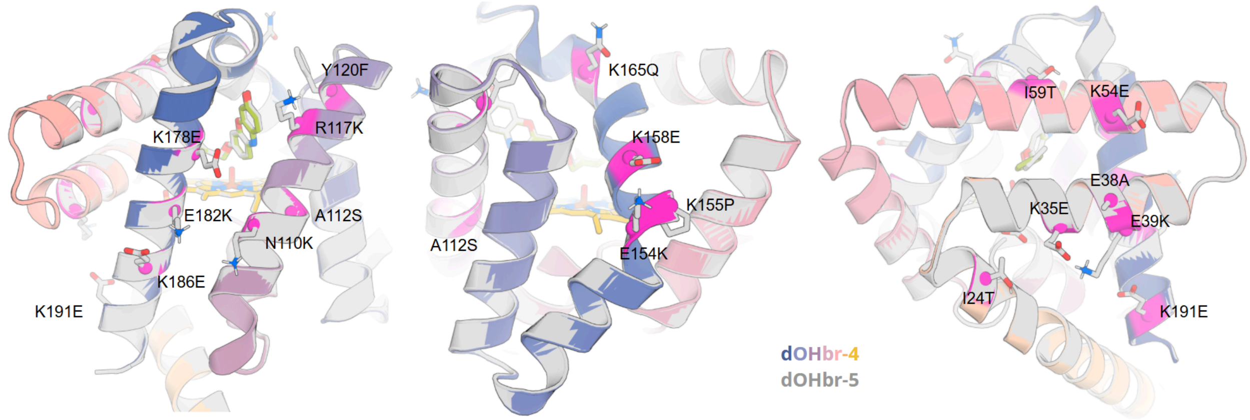


**Figure S34.** Visualization of **dOHbr-4** (colored) and **dOHbr-5** (gray) sequence differences at 15 positions. Residues in **dOHbr-5** as shown as gray sticks. The active site sequences are identical, and Y120F and R117K are the only other differences close to the substrate. Rest of the mutations are all on the surface, introduced by the second sphere ligandMPNN sequence redesign in stage #4 of the design workflow.

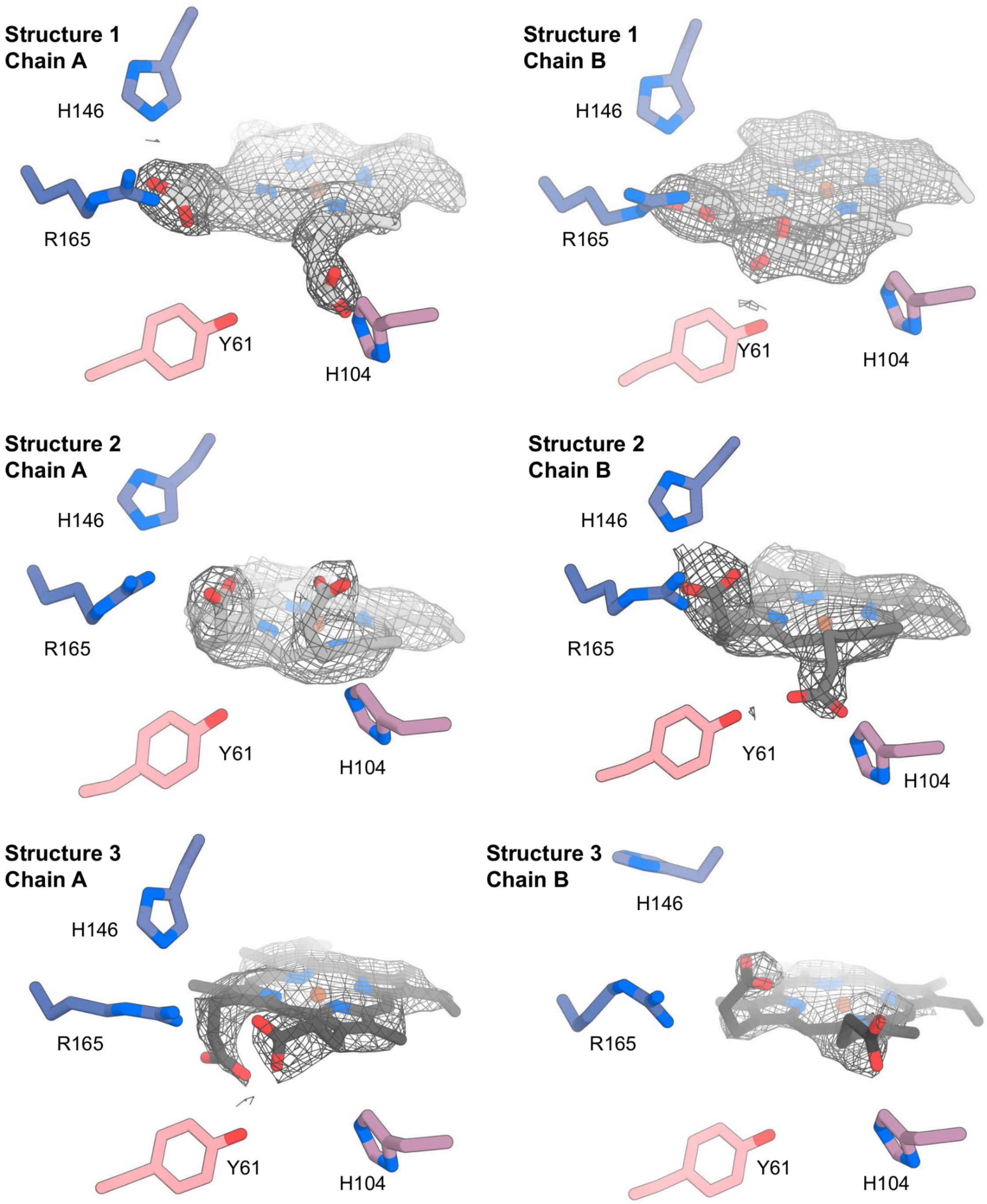


**Figure S35.** Structures and densities (2*F*o–*F*c map contoured at 1σ) of heme in the six asymmetric chains of three crystal structures of **dOHbr-2**. The structures 1-3 were solved to 2.0, 2.9 and 3.0 Å resolution, respectively. The heme propionate close to His104 shows a great degree of flexibility.

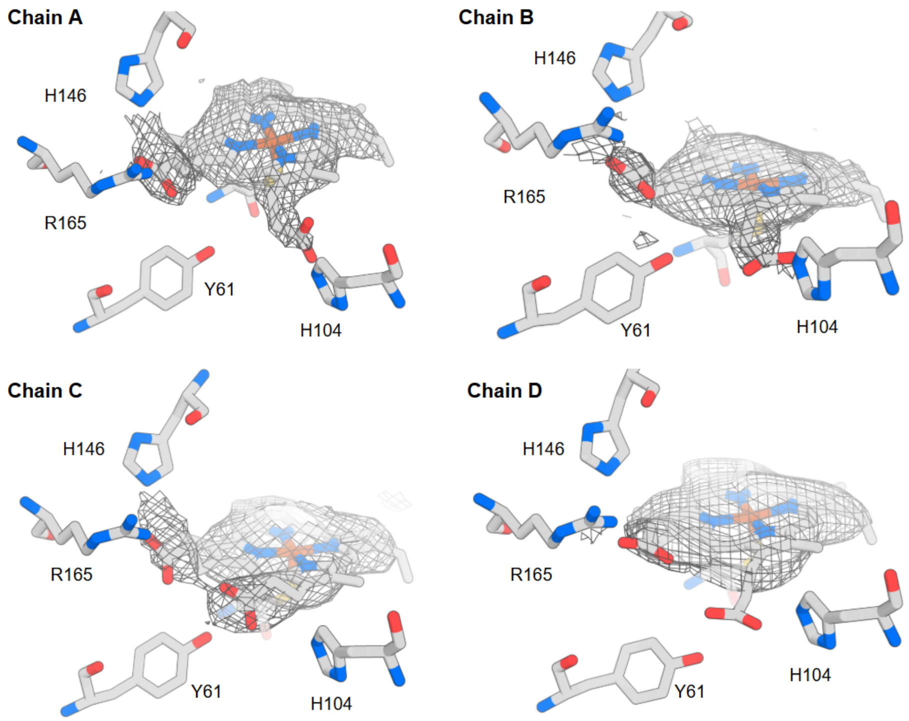


**Figure S36.** Structures and densities (2*F*o–*F*c map contoured at 1σ) of heme in the four asymmetric chains of the crystal structure of **dOHbr-2**-L125M-L150F. The structure was solved to 2.9 Å resolution. The structures show heme propionate closer to His104 adopting a similar conformation in chains A, B and D.

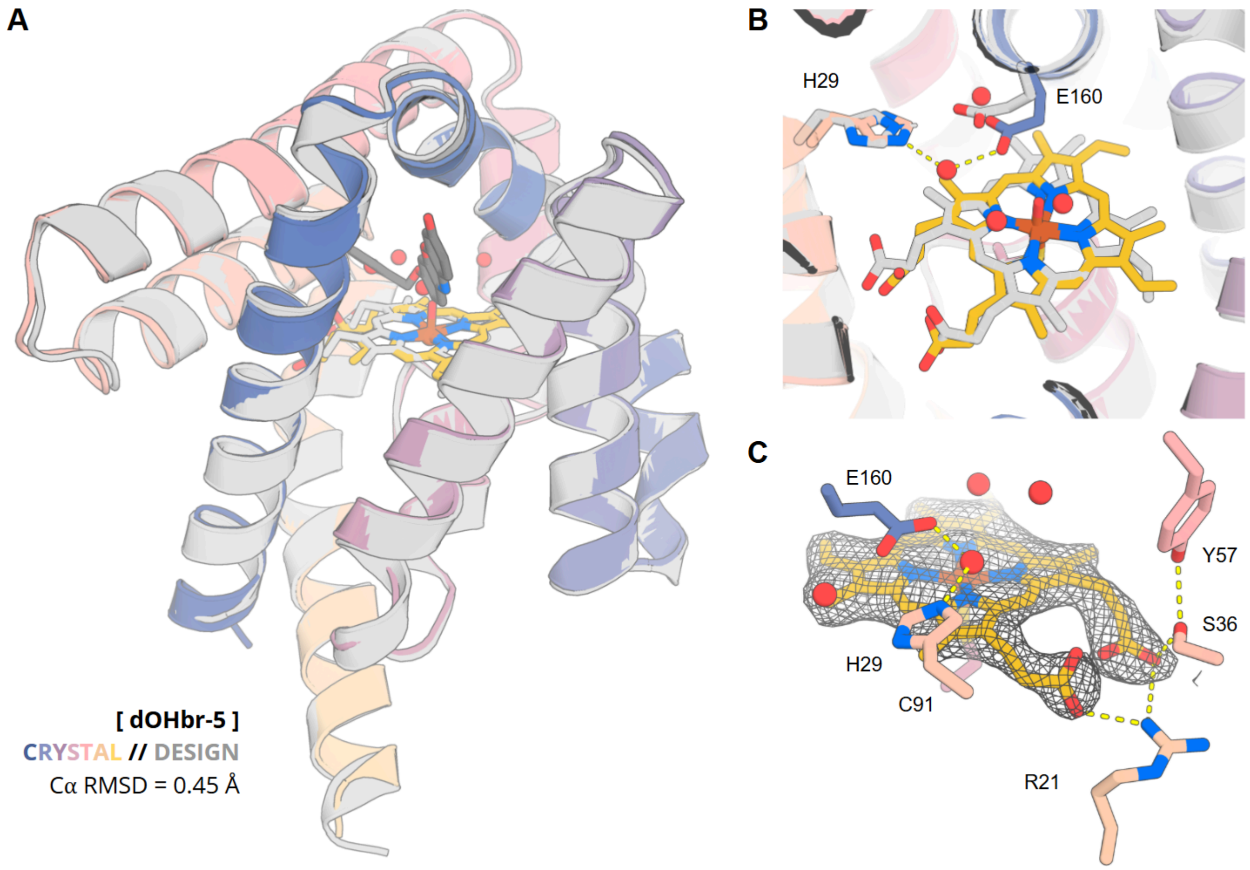


**Figure S37.** Crystal structure of **dOHbr-5**, solved to 2.2 Å resolution. **(A)** Overlay of the crystal structure (colored) with the design model (gray) shows a very close agreement, with $C_\alpha$ RMSD of 0.45 Å. **(B)** Positioning of the heme closely matches the design. The Glu160-His29 catalytic dyad is water-bridged in the crystal structure. **(C)** Density (2*F*o–*F*c map contoured at 1σ) of heme in the crystal structure, and its hydrogen bond contacts with the protein residues Arg-21, Ser-36 and Tyr-57.

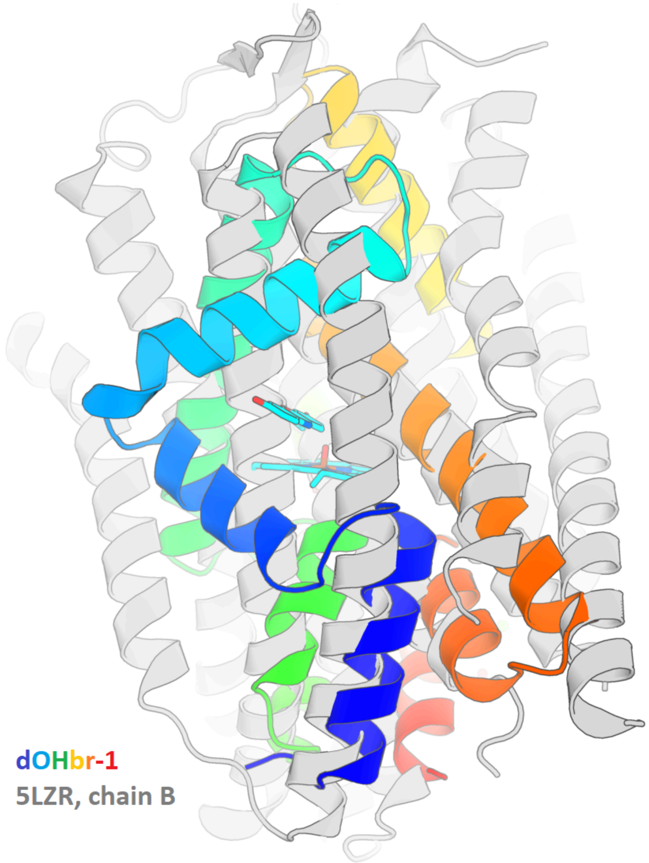


**Figure S38.** Alignment of **dOHbr-1** (rainbow) with its closest match in the PDB (5LZR, chain B; gray). TM-score of 0.53 indicates low similarity between the structures, and the alignment shows that only a few individual secondary structure elements match.

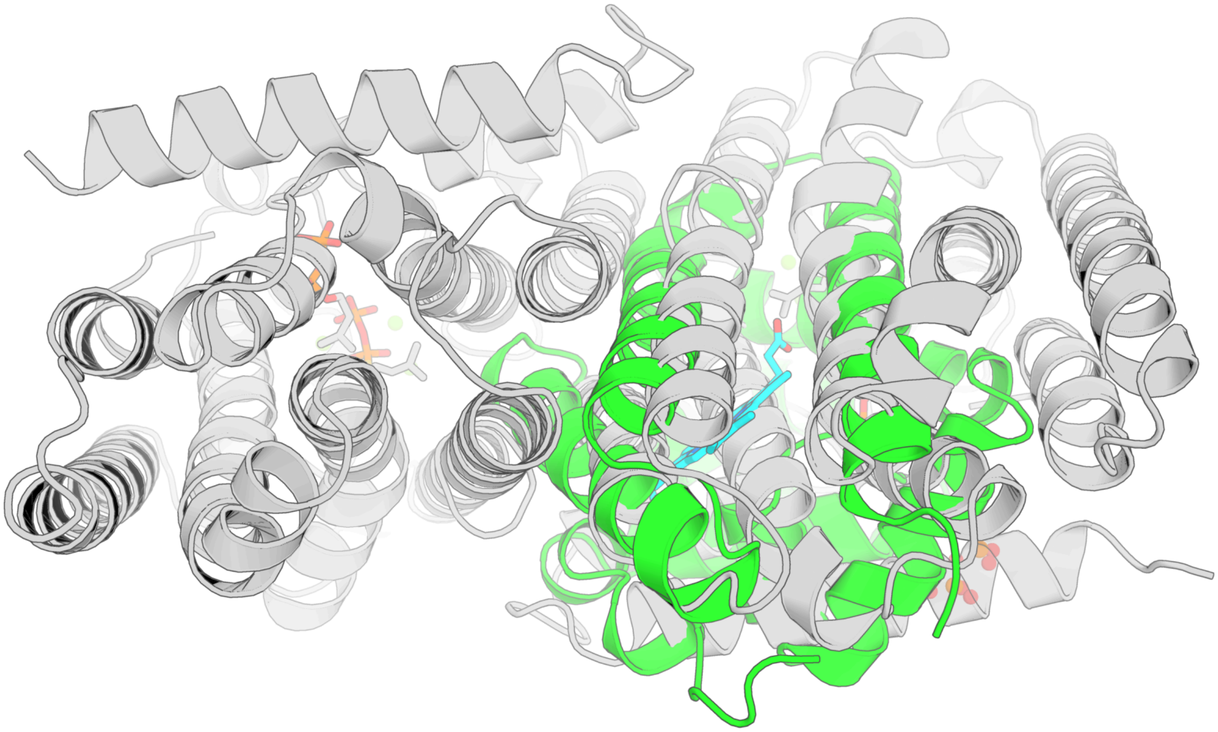

**Figure S39.** Alignment of **dOHbr-2** (green) with its closest match in the PDB (1RQI, gray). TM-score of 0.50 indicates no substantial similarity between the structures, and the alignment shows that only a few individual secondary structure elements match.

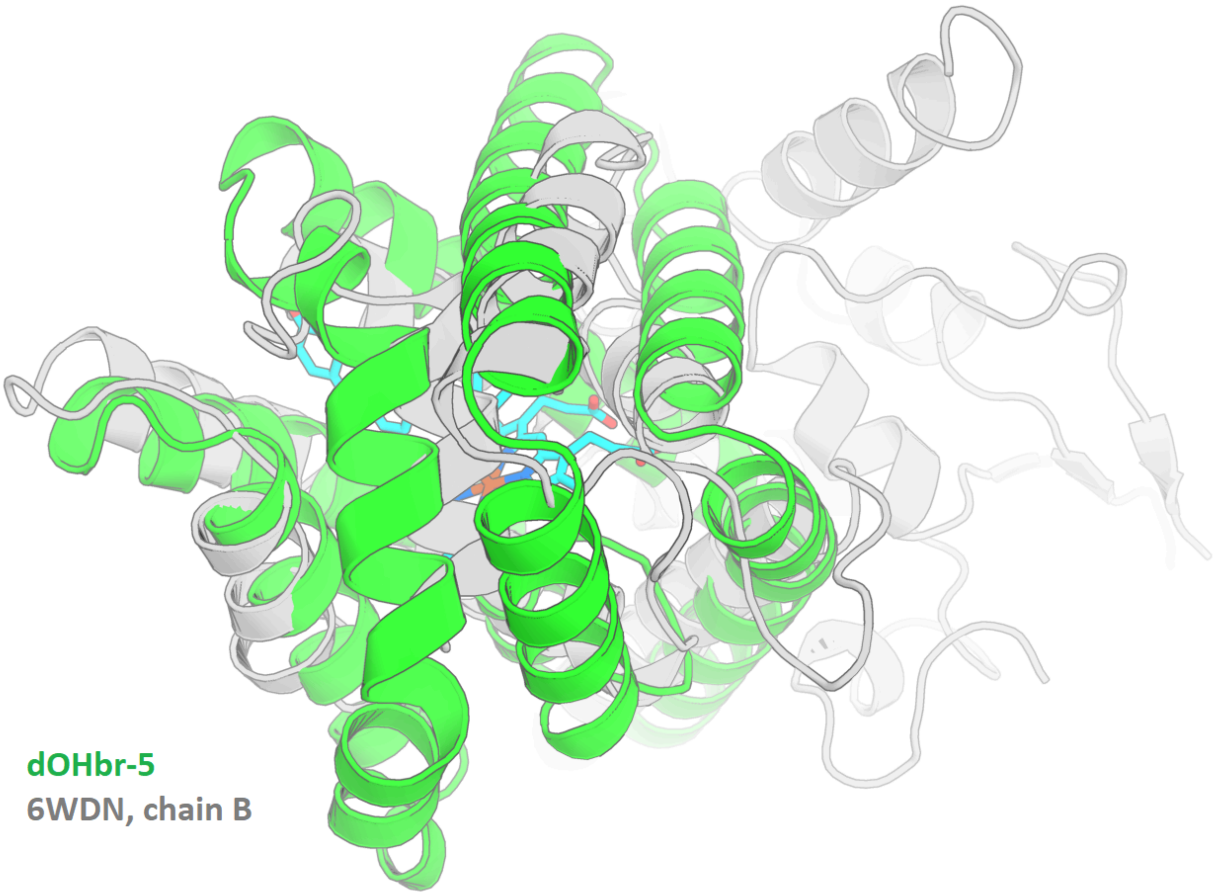


**Figure S40.** Alignment of **dOHbr-5** (green) with its closest match in the PDB (6WDN, chain B; gray). TM-score of 0.57 indicates some similarity between the structures. The alignment shows that the overall topology of **dOHbr-5** loosely follows that of 6WDN_B at positions 295-441, but the connections between secondary structure elements differ significantly.

**Table S12.** The extinction coefficients of Soret band for designs.

| | **Extinction coefficient (mM$^{-1}$ cm$^{-1}$)** |
|---|---|
| **dOHbr-2** | $A_{387nm}$ 102 ± 2 |
| **dOHbr-2** | $A_{387nm}$ 118 ± 9 |
| **dOHbr-2** | $A_{412nm}$ 105 ± 2 |
| **dOHbr-2**-V29L-Y61W | $A_{412nm}$ 132 ± 7 |
| **dOHbr-2**-Y61W-V72K | $A_{412nm}$ 127 ± 3 |

### 4. Estradiol hydroxylation

#### 4.1. Computational methods

Two design campaigns were run in parallel, differing only in the targeted regiochemistry of estradiol (RCSB `EST`) hydroxylation: campaign C7C6C5, which places C6 at the catalytic position, and campaign C17C16C15, which places C16 there. In the three-atom regiochemistry label the middle atom is the carbon being hydroxylated. The theozymes are generated in the same way as previous aliphatic hydroxylation pipelines. Both campaigns used the same ferryl heme-thiolate theozyme, the same pipeline, the same software versions and the same selection criteria. Everything below therefore describes both campaigns; where they differ, both values are given.

RFdiffusion3

Starting from 129 theozyme rotamer inputs, we generated backbones of 175–230 amino acids with RFdiffusion3. Each input was run with a diffusion batch size of 5 in 10 batches, giving 50 backbones per input and 6,450 per campaign run. Unindexed scaffolding was used for the catalytic motif. We used a single ORI token position per input, placed on the ferryl oxygen (`HOM` atom O5) in both campaigns. All heavy atoms of the heme (`HOM`, the Fe(IV)=O Compound I model) and of the substrate were held fixed, as were the catalytic side chains. Most of the substrate heavy atoms were additionally assigned to be buried. The two campaigns differ here, because the regiochemistry determines which end of the steroid faces the iron: in C7C6C5 the phenolic A-ring oxygen (O2) is buried and declared an H-bond donor while the D-ring carbonyl oxygen (O1) is only partially buried, and in C17C16C15 the assignment is reversed, with O1 buried and declared an H-bond acceptor.

RFD3 input JSON (C7C6C5; C17C16C15 values given in comments):

```
{"input": "input.pdb",
 "ligand": "HOM,EST",
 "unindex": "A1-5,A6,A8",
 "length": "175-230",
 "ori_token":[1.680,0.0,0.0]
 "dialect": 2,
 "select_unfixed_sequence": "A4-5",
 "select_fixed_atoms":{
        "A1":"N,CA,C,O,CB,CG,CD",
        "A2":"N,CA,C,O,CB,SG",
        "A3":"N,CA,C,O,CB,CG,CD",
        "A4":"N,CA,C,O",
        "A5":"N,CA,C,O",
        "A6":"ND1,CG,NE2,CD2,CB,CE1",
        "A8":"OE2,OE1,CD,CG,CB",
        "HOM":"ALL",
        "EST":"ALL"},
 "select_buried": {"HOM": "FE1,C17,C20,C24,C27,C13,C34",
        "EST": "C1,C2,C3,C4,C5,C6,C7,C8,C9,C10,C11,C12,C13,C14,C15,C16,C18,O2"},
                                # O1 in place of O2 for C17C16C15
 "select_partially_buried": {"HOM": "O1,O2,O3,O4,C5,C10"
        "EST": "O1"},           # O2 for C17C16C15
```

Diffused backbone filtering

All generated backbones were evaluated for backbone and pocket quality (Table S13), selecting proteins with no chain breaks or clashes against the ligands, reasonable side-chain geometry at the scaffolded positions, less than 35% loop content, no helix longer than 36 residues, a compact radius of gyration, both termini more than 15 Å from any ligand, and a buried ligand pocket. In the final diffusion run, 4.2% of backbones passed for C7C6C5 and 9.0% of backbones passed for C17C16C15.

**Table S13.** Generated backbone selection criteria. The partial-diffusion outputs (next section) were filtered on the same metrics with two tightened cutoffs, given in parentheses.

| Metric | Cutoff | Description |
|---|---|---|
| chainbreak | <= 4.5 Å | Largest adjacent Ca-Ca distance |
| rCA_nonadj | >= 3.0 Å (3.5 Å) | Smallest non-adjacent Ca-Ca distance. Always >3.6 Å in the PDB. |
| lig_dist | >= 2.5 Å | Smallest distance between a ligand heavyatom and a protein backbone heavyatom |
| bondlen_dev | <= 1.0 Å | Largest bond-length deviation from physical ground truth |
| cart_bonded_avg | <= 10.0 | Average Rosetta cart_bonded score of scaffolded residues. Shows side-chain geometry quality. |
| fa_dun_avg | <= 15.0 | Average Rosetta fa_dun score of scaffolded residues. Shows rotamer quality. |
| loop_frac | <= 35% | Percentage of loops in the protein, based on Rosetta DSSP analysis. |
| longest_helix | <= 36 | Longest helix in the protein, based on Rosetta DSSP analysis. |
| rog | <= 20.0 Å | Radius of gyration of the protein. |
| term_mindist | >= 15.0 Å | Distance between the termini and any ligand. |
| SASA_rel | <= 0.25 (0.10) | Relative SASA of the ligands (SASA_bound / SASA_free). |

Backbone diversification with partial diffusion

We used RFD3 partial diffusion to sample the backbone space around each surviving scaffold. At a low noise level the procedure generates perturbed variants in which the structural elements have shifted slightly. We used a `partial_t` noise level of 4.0 and generated 5 variants per input backbone. All heavy atoms of ligands were held fixed. The catalytic cysteine was fixed in all its heavy atoms, whereas for the distal histidine and glutamate only the side-chain atoms were fixed, allowing partial diffusion to continue to move their backbone.

RFD3 partial diffusion input JSON:
```
{"design_name":{"input":"diffusion_output.pdb",
 "ligand":"HOM,EST",
 "partial_t":4.0,
 "contig":"A42,A68,A123",   # example; the scaffolded positions of that input
 "select_fixed_atoms":{"A42":"N,CA,C,O,CB,SG",
                       "A68":"CB,CG,ND1,CD2,CE1,NE2",   # His and Glu: side chain only
                       "HOM":"ALL",
                       "EST":"ALL"}}}
```

The partial-diffusion outputs were analysed on the same backbone quality metrics (Table S1, tightened cutoffs), with 8.9% and 8.9% passing for both C7C6C5 and C17C16C15.

**Iterative sequence design and structure prediction**

To turn the generated backbones into proteins whose sequence folds into the intended shape while retaining precise positioning of the heme, the substrate and all catalytic residues, we used a four-stage design strategy that becomes progressively more refined in the sequence design and progressively more stringent in the selection.

In all stages, structural constraints were applied during Rosetta repacking, minimization and relaxation. The constraints were applied with the Rosetta AddOrRemoveMatchCsts mover and provided in the enzdes constraint format as four blocks: heme-Cys, the reactive geometry between the ferryl oxygen and the target carbon of the substrate, heme-Glu, and the Glu-His dyad.

<u>#1 Predesign</u>

In the first stage we used ligandMPNN (T=0.1) to generate a single sequence for each diffused and partially diffused backbone. Each structure was then repacked, minimized and coarsely relaxed in torsional space, all under enzdes constraints. Minimization was necessary to resolve bond length and angle problems in the catalytic side chains introduced by tip-atom-based scaffolding, and applying the constraints also absorbed the small deviations from ideal geometry that RFD3 can introduce in the motif atoms. The structures were then loosely analysed for active site quality in terms of constraint score, ligand SASA, contact molecular surface and ddG (Table S14) with cutoffs set only to exclude the worst backbones. This step retained 69% for C7C6C5 and 72% for C17C16C15.

**Table S14.** Stage #1 predesign selection criteria.

| Metric | Cutoff | Description |
|---|---|---|
| all_cst | <= 3.0 | Weighted constraint score - how well are constraints respected. |
| cms_per_atom_HOM | >= 5.5 | Contact molecular surface of heme, per ligand atom |
| cms_per_atom_EST | >= 4.0 | Contact molecular surface of estradiol, per ligand atom |
| L_SASA_HOM | <= 0.05 | Relative SASA of heme |
| L_SASA_EST | <= 0.15 | Relative SASA of estradiol |
| corrected_ddg | <= -55.0 | Rosetta ddG between the protein and all ligands (two-body energies). |

<u>#2 Backbone sequence design and folding</u>

We generated new sequences for each structure using enhancedMPNN at T=0.1, 0.2 and 0.3 with 3 sequences each (9 per structure), omitting Cys and Met and fixing the identities of the catalytic residues. The resulting sequences were predicted with AlphaFold3 without any ligands, focusing purely on folding of the *apo* protein; the intention was to find scaffolds and sequences that fold into the correct shape without the aid of the heme or substrate. Predictions were run in single-sequence mode with 6 recycles and 1 diffusion sample. Selection criteria (Table S15) focus on protein confidence (pLDDT, pTM) and accuracy (RMSD). 3.8% passed for C7C6C5 and 2.4% for C17C16C15, unique diffused backbones (16% and 13%, respectively).

**Table S15.** AlphaFold3 *apo* prediction selection criteria.

| Metric | Cutoff | Description |
|---|---|---|
| chain_ptm_ch0 | >= 0.75 | Global structural accuracy of protein chain A (predicted TM score) |
| rmsd | <= 1.5 Å | Cα RMSD between the design model and the prediction. |
| plddt | >= 75.0 | All-atom local structural accuracy of protein chains. |
| plddt_BB | >= 80.0 | Local structural accuracy of protein chains, averaged over backbone atoms (C,N,CA,O). |
| rmsd_SR1 | <= 1.5 Å | All-atom RMSD of CYS |
| plddt_SR1 | >= 70.0 | All-atom confidence of CYS |
| rmsd_SR3 | <= 2.0 Å | All-atom RMSD of GLU |
| plddt_SR3 | >= 70.0 | All-atom confidence of GLU |
| rmsd_SR4 | <= 2.5 Å | All-atom RMSD of HIS |
| plddt_SR4 | >= 70.0 | All-atom confidence of HIS |

Finally, the successful AF3 predicted structures were aligned to the corresponding design models (enhancedMPNN inputs), the heme and the substrate transplanted directly into the AF3 model, and this structure saved as a new PDB file.

#3 Active site sequence design

The third stage optimizes the sequence of the active site. For each structure from the previous step, positions with Cα within 9–15 Å of any ligand heavy atom were selected as designable (catalytic residues were excluded). The design used an iterative ligandMPNN and Rosetta repack/minimize protocol implemented through the FastMPNNdesign package. The protocol can be referred to the previous computational section using FastMPNNdesign. Because the regiochemistry determines which end of the steroid is solvent-facing, the atom defining those positions is the D-ring carbonyl oxygen O1 in the C7C6C5 campaign and the phenolic oxygen O2 in the C17C16C15 campaign.

FastMPNNdesign protocol, stage #3:

```
scale:coordinate_constraint 1.0
scale:fa_rep 0.150
mpnn 1.0 10
repack
scale:fa_rep 0.200
min 0.01
filter cst 0.0 8.0
keep_best 2 cms
mpnn_selector keep_hbonds_to_ligand_and_catres
scale:coordinate_constraint 0.2
scale:fa_rep 0.8
mpnn 0.4 3
repack
min 0.01
filter cst 0.0 6.0
keep_best 5 cms
keep_best 3 ddg
mpnn_selector keep_hbonds_to_ligand_and_catres
scale:coordinate_constraint 0.0
scale:fa_rep 1.0
mpnn 0.2 1
repack
min 0.01
filter cst 0.0 4.0
keep_best 2 cms
mpnn_selector keep_hbonds_to_ligand_and_catres
mpnn 0.1 5
repack
min 0.00001
```

The resulting design models were evaluated on Rosetta-based metrics describing the active site (Table S16). In addition, each model was relaxed again after deleting the substrate and analysed for the stability of the backbone and active site, the goal being to avoid designs in which the substrate is critical to the integrity of the pocket. These criteria retained 25% for C7C6C5 and 61% for C17C16C15, corresponding to 81 and 104 unique scaffolds.

**Table S16.** Stage #3 Rosetta filtering criteria.

| Metric | Cutoff | Description |
|---|---|---|
| all_cst | <= 4.0 | Weighted constraint score - how well are constraints respected. |
| cms_per_atom_HOM | >= 5.75 | Contact molecular surface of heme |
| cms_per_atom_EST | >= 5.0 | Contact molecular surface of estradiol |
| L_SASA_HOM | <= 0.05 | Relative SASA of heme |
| L_SASA_EST | <= 0.10 | Relative SASA of estradiol |
| corrected_ddg | <= -72.0 | Rosetta ddG between the protein and all ligands (two-body energies). |
| YW_dist | >= 6.0 Å | Distance between iron-oxo and the closest Tyr/Trp side chain |
| YW_count | <= 1.0 | Number of Tyr/Trp residues within 7 Å of iron-oxo. |
| KRE_count | <= 1.0 | Number of Lys, Arg, Glu residues touching the back of the substrate. Avoids salt bridges blocking substrate entry. |
| rmsd_CA_NoSub | <= 0.8 Å | Protein Cα RMSD after relaxing without the substrate. |
| rmsd_sc_NoSub | <= 1.0 Å | Average pocket residue all-atom RMSD after relaxing without the substrate. |
| rmsd_HOM_NoSub | <= 0.8 Å | Heme RMSD after relaxing without the substrate. |
| ddg_NoSub | <= -68.0 | Rosetta ddG between the protein and heme after relaxing without the substrate. |
| cms_per_atom_NoSub | >= 5.7 | Contact molecular surface of heme after relaxing without the substrate. |

LigandMPNN was then used to generate additional sequence diversity for the passing designs. For this we selected the outer layer of residues that are not designed in the FastMPNNdesign iteration. We generated 5 sequences each at T=0.1 and 0.2 giving 10 diversified sequences plus the original per design.

All sequences corresponding to passing designs were then co-folded with heme and estradiol using AlphaFold3 with 10 recycles and 5 diffusion samples with no MSA and no template. Heme was represented as the CCD_HEM model rather than the Compound I used for design, and estradiol was supplied as a userCCD mmCIF input with predefined atom names and bonding. Selection at this point focused on precise and confident recapitulation of the catalytic residue and heme-binding geometries, and did not yet filter on substrate binding (Table S17). Each of the five diffusion samples was assessed individually. These criteria retained 15% for C7C6C5 and 8.2% for C17C16C15.

**Table S17.** AlphaFold3 *holo* prediction selection criteria for heme binding in stage #3.

| Metric | Cutoff | Description |
|---|---|---|
| chain_ptm_ch0 | >= 0.80 | Global structural accuracy of protein chain A (predicted TM score) |
| chain_iptm_ch0 | >= 0.75 | Interface structural accuracy of protein chain A |
| chain_ptm_ch1 | >= 0.75 | Global structural accuracy of heme (predicted TM score) |
| chain_iptm_ch1 | >= 0.65 | Interface structural accuracy of heme |
| chain_pair_pae_min_inter | <= 5.0 | Largest interchain minPAE value (average of off-diagonal elements in the PAE matrix). |
| rmsd | <= 1.2 Å | Cα RMSD between the design model and the prediction |
| plddt | >= 85.0 | All-atom local structural accuracy of protein chains. |
| plddt_BB | >= 87.0 | Local structural accuracy of protein chains, averaged over backbone atoms (C,N,CA,O). |
| lig0_rmsd | <= 2.0 Å | Heme RMSD |
| lig0_plddt | >= 75.0 | Heme pLDDT |

| | | |
|---|---|---|
| lig1_rmsd | <= 4.0 Å | estradiol RMSD. Deliberately permissive at this stage. |
| rmsd_SR1 | <= 1.0 Å | All-atom RMSD of CYS |
| plddt_SR1 | >= 85.0 | All-atom confidence of CYS |
| rmsd_SR3 | <= 1.6 Å | All-atom RMSD of GLU |
| plddt_SR3 | >= 82.0 | All-atom confidence of GLU |
| rmsd_SR4 | <= 1.6 Å | All-atom RMSD of HIS |
| plddt_SR4 | >= 82.0 | All-atom confidence of HIS |

Finally, the successful AF3 predicted structures were aligned to the corresponding design models, the substrate transplanted directly into the AF3 model, and this structure saved as a new PDB file. The heme was taken from the AF3 prediction.

#4 Refinement of the substrate binding site

The fourth stage samples the sequence space around the substrate more deeply. Here the designable layer was narrowed to Cα within 7–13 Å of the ligands, and positions closer to the heme-coordinating cysteine than to the substrate. The protocol can be referred to the previous computational section using FastMPNNdesign. Slightly more stringent Rosetta selection criteria were applied to the design models (Table S18), which retained 20% for C7C6C5 and 8.4% for C17C16C15.

FastMPNNdesign protocol, stage #4:

```
scale:coordinate_constraint 0.5
scale:fa_rep 0.75
mpnn 0.4 10
repack
min 0.01
filter cst 0.0 6.0
keep_best 5 cms
mpnn_selector keep_hbonds_to_ligand_and_catres
scale:coordinate_constraint 0.0
scale:fa_rep 1.0
mpnn 0.2 1
repack
min 0.01
filter cst 0.0 4.0
keep_best 3 cms
keep_best 2 ddg
mpnn_selector keep_hbonds_to_ligand_and_catres
mpnn 0.1 5
repack
min 0.00001
```

**Table S18.** Stage #4 Rosetta filtering criteria.

| Metric | Cutoff | Description |
|---|---|---|
| all_cst | <= 3.0 | Weighted constraint score - how well are constraints respected. |
| cms_per_atom_HOM | >= 6.0 | Contact molecular surface of heme |
| cms_per_atom_EST | >= 5.1 (5.5) | Contact molecular surface of estradiol. C7C6C5 (C17C16C15). |
| L_SASA_HOM | <= 0.05 | Relative SASA of heme |
| L_SASA_EST | <= 0.05 | Relative SASA of estradiol |
| substrate_SASA | >= 0.01 | Absolute SASA of estradiol. Selects those with at least some exposed area. |
| corrected_ddg | <= -75.0 | Rosetta ddG between the protein and all ligands (two-body energies). |
| YW_dist | >= 8.0 Å | Distance between iron-oxo and the closest Tyr/Trp side chain |
| YW_count | <= 0.0 | Number of Tyr/Trp residues within 7 Å of iron-oxo. |
| KRE_count | <= 1.0 | Number of Lys, Arg, Glu residues touching the back of the substrate. |
| rmsd_CA_NoSub | <= 0.7 Å | Protein Cα RMSD after relaxing without the substrate. |
| rmsd_sc_NoSub | <= 1.0 Å | Average pocket residue all-atom RMSD after relaxing without the substrate. |
| rmsd_HOM_NoSub | <= 0.8 Å | Heme RMSD after relaxing without the substrate. |
| ddg_NoSub | <= -70.0 | Rosetta ddG between the protein and heme after relaxing without the substrate. |
| cms_per_atom_NoSub | >= 5.8 | Contact molecular surface of heme after relaxing without the substrate. |

All sequences corresponding to passing designs (1 original + 10 diversified) were again co-folded with heme and estradiol using AlphaFold3 with 10 recycles and 5 diffusion samples. Selection now addressed the entire active site, including the catalytic residues, the heme and the substrate (Table S19). Most metrics are averaged across the five diffusion samples, and the RMSDs are additionally filtered on the worst value among the five, ensuring that a sequence folds reliably into the same geometry rather than only on average. These criteria retained 10% for C7C6C5 and 16% for C17C16C15.

**Table S19.** AlphaFold3 *holo* prediction selection criteria in stage #4.

| Metric | Cutoff | Description |
|---|---|---|
| chain_ptm_ch0 | >= 0.85 | Global structural accuracy of protein chain A (predicted TM score), average of 5 models |
| chain_iptm_ch0 | >= 0.80 | Interface structural accuracy of protein chain A, average of 5 models |
| chain_ptm_ch1 | >= 0.75 | Global structural accuracy of heme (predicted TM score), average of 5 models |
| chain_iptm_ch1 | >= 0.70 | Interface structural accuracy of heme, average of 5 models |
| chain_iptm_ch2 | >= 0.70 | Interface structural accuracy of estradiol, average of 5 models |
| chain_pair_pae_min_inter_max | <= 3.0 | Largest average interchain minPAE value from 5 models. |
| rmsd | <= 1.2 Å | Cα RMSD between the design model and the prediction, average of 5 models |
| plddt | >= 85.0 | All-atom local structural accuracy of protein chains. |
| plddt_BB | >= 87.0 | Local structural accuracy of protein chains, averaged over backbone atoms (C,N,CA,O). |
| lig0_rmsd | <= 2.0 Å | Heme RMSD, average of 5 models |
| lig0_rmsd_max | <= 2.5 Å | Worst heme RMSD among 5 models. |
| lig0_plddt | >= 80.0 | Heme pLDDT, average of 5 models |
| lig1_rmsd | <= 3.0 Å | estradiol RMSD, average of 5 models |
| lig1_rmsd_max | <= 4.0 Å | Worst estradiol RMSD among 5 models. |
| lig1_plddt | >= 70.0 | estradiol pLDDT, average of 5 models |
| rmsd_SR1 | <= 0.8 Å | All-atom RMSD of CYS, average of 5 models |
| rmsd_SR1_max | <= 1.0 Å | Worst CYS RMSD among 5 models. |
| plddt_SR1 | >= 85.0 | All-atom confidence of CYS, average of 5 models |
| rmsd_SR3 | <= 1.2 Å | All-atom RMSD of GLU, average of 5 models |
| rmsd_SR3_max | <= 1.5 Å | Worst GLU RMSD among 5 models. |
| plddt_SR3 | >= 85.0 | All-atom confidence of GLU, average of 5 models |
| rmsd_SR4 | <= 1.4 Å | All-atom RMSD of HIS, average of 5 models |
| rmsd_SR4_max | <= 1.6 Å | Worst HIS RMSD among 5 models. |
| plddt_SR4 | >= 85.0 | All-atom confidence of HIS, average of 5 models |

#### Final analysis of designs

##### <u>Constrained relax</u>

All models that passed the stage #4 AF3 selection were relaxed with Rosetta FastRelax. Heavy coordinate constraints were applied to the backbone to retain the AF3-predicted geometry, and enzdes constraints were applied to nudge the catalytic residues, heme and substrate towards the idealized orientation. The relaxed structures were re-scored on the active site and ligand interaction terms (Table S20). This was done for all five AF3 samples, giving each sequence several chances to provide a good structural model. 7.8% of relaxed models passed for C7C6C5 11% for C17C16C15.

**Table S20.** Constrained-relax selection criteria.

| Metric | Cutoff | Description |
| --- | --- | --- |
| all_cst | <= 4.0 | Weighted constraint score - how well are constraints respected. |
| cms_per_atom_HOM | >= 5.9 | Contact molecular surface of heme |
| cms_per_atom_EST | >= 5.32 (5.8) | Contact molecular surface of estradiol. C7C6C5 (C17C16C15). |
| L_SASA_HOM | <= 0.05 | Relative SASA of heme |
| L_SASA_EST | <= 0.10 | Relative SASA of estradiol |
| sc_EST | >= 0.50 | Shape complementarity between the protein and estradiol. |
| corrected_ddg | <= -70.0 | Rosetta ddG between the protein and all ligands (two-body energies). |
| COO_hbond | = True | Do both propionate groups have at least one H-bond partner each. |
| HBres_not_STY | = True | Not all H-bonds to the propionate groups come from Ser, Thr, Tyr. |
| YW_dist | >= 7.0 Å | Distance between iron-oxo and the closest Tyr/Trp side chain |
| YW_count | <= 1.0 | Number of Tyr/Trp residues within 7 Å of iron-oxo. |
| KRE_count | <= 1.0 | Number of Lys, Arg, Glu residues touching the back of the substrate. |
| rmsd_CA_NoSub | <= 0.8 Å | Protein Cα RMSD after relaxing without the substrate. |
| rmsd_sc_NoSub | <= 1.0 Å | Average pocket residue all-atom RMSD after relaxing without the substrate. |
| rmsd_HOM_NoSub | <= 0.8 Å | Heme RMSD after relaxing without the substrate. |
| ddg_NoSub | <= -75.0 | Rosetta ddG between the protein and heme after relaxing without the substrate. |
| cms_per_atom_NoSub | >= 5.8 | Contact molecular surface of heme after relaxing without the substrate. |

Heme-only AlphaFold3 predictions

To confirm that the active site remains preorganized without the substrate, and that folding is not substrate-dependent, we ran further AlphaFold3 predictions of the passing designs omitting estradiol. Predictions used 10 recycles and 5 diffusion samples with heme as CCD_HEM, without MSA or template. The designs were selected on the stage #4 criteria (Table S19) excluding the substrate terms (the “ch2” and “lig1” scores), with `rmsd` tightened to 1.0 Å, `chain_pair_pae_min_inter_max` to 2.5 and `lig0_plddt` set to 79.0. 49% designs passed for C7C6C5 and 29% for C17C16C15.

PLACER analysis

We used PLACER to analyse the designed active sites for conformational peorganization. Estradiol was docked into the cropped active site while the heme was held fixed. We generated 100 samples per design using the relaxed AF3 models as input and ranked them by predicted RMSD. The predictions were analysed for the accuracy (RMSD) and confidence (pRMSD) of the catalytic side chains and the substrate. For the substrate we focused on the five highest-confidence predictions, since many lower-confidence samples had poorly predicted geometries. We also used the p_near metric of the substrate docking funnel (RMSD vs pRMSD), which measures its “funnelness”. Table S21 lists the criteria; 53% designs passed for C7C6C5 and 76% for C17C16C15.

**Table S21.** PLACER analysis selection criteria.

| Metric | Cutoff | Description |
| --- | --- | --- |
| rmsd_prmsd_lig_EST_top5 | <= 1.6 Å | Average RMSD of estradiol, among the top 5 models ranked by pRMSD |
| plddt_top5 | >= 0.85 | Average 1D track confidence (pLDDT) of the top 5 models, ranked by pLDDT |

| | | |
|---|---|---|
| plddt_pde_top5 | >= 0.75 | Average 2D track confidence (pLDDT-pde) of the top 5 models, ranked by pLDDT-pde |
| prmsd_top5 | <= 2.5 | Average substrate pRMSD of the top 5 models, ranked by pRMSD |
| prmsd_pnear_EST | >= 0.3 | Funnelness of the substrate RMSD-pRMSD relationship |
| rmsd_catres1 | <= 0.4 Å | Average RMSD of Cys, across 100 models |
| rmsd_catres2 | <= 1.3 Å | Average RMSD of Glu, across 100 models |
| rmsd_catres3 | <= 1.3 Å | Average RMSD of His, across 100 models |
| u_catres1 | <= 0.4 | Average uncertainty (pRMSD) of Cys, across 100 models |
| u_catres2 | <= 1.0 | Average uncertainty (pRMSD) of Glu, across 100 models |
| u_catres3 | <= 1.0 | Average uncertainty (pRMSD) of His, across 100 models |
| u_pocket | <= 0.5 | Average uncertainty (pRMSD) of all pocket residues (within 5 Å of the substrate), across 100 models |
| rmsd_pocket_worst | <= 3.0 Å | Worst average side-chain RMSD of any pocket residue |
| u_pocket_worst | <= 2.0 | Worst average pRMSD of any pocket residue |

RoseTTAFold3 predictions

As an final orthogonal structure-prediction check, the designs were additionally co-folded with heme and estradiol using RoseTTAFold3, with heme supplied as CCD_HEM and estradiol as a custom mmCIF. The criteria (Table S22) mirror the stage #4 AF3 selection. 36% designs passed for C7C6C5 and 6.9% for C17C16C15.

**Table S22.** RoseTTAFold3 selection criteria.

| Metric | Cutoff | Description |
|---|---|---|
| ptm | >= 0.85 | Global structural accuracy (predicted TM score), average of 5 models |
| iptm | >= 0.85 | Interface structural accuracy, average of 5 models |
| chain_pair_pae_min_inter_max | <= 5.0 | Largest average interchain minPAE value from 5 models. |
| rmsd | <= 1.2 Å | Cα RMSD between the design model and the prediction, average of 5 models |
| plddt | >= 85.0 | All-atom local structural accuracy of protein chains. |
| plddt_BB | >= 88.0 | Local structural accuracy of protein chains, averaged over backbone atoms (C,N,CA,O). |
| lig0_rmsd | <= 2.0 Å | Heme RMSD, average of 5 models |
| lig0_rmsd_max | <= 2.5 Å | Worst heme RMSD among 5 models. |
| lig0_plddt | >= 85.0 | Heme pLDDT, average of 5 models |
| lig1_rmsd | <= 1.6 Å | estradiol RMSD, average of 5 models |
| lig1_rmsd_max | <= 2.0 Å | Worst estradiol RMSD among 5 models. |
| lig1_plddt | >= 80.0 | estradiol pLDDT, average of 5 models |
| rmsd_SR1 | <= 0.8 Å | All-atom RMSD of CYS, average of 5 models |
| rmsd_SR1_max | <= 1.0 Å | Worst CYS RMSD among 5 models. |
| plddt_SR1 | >= 85.0 | All-atom confidence of CYS, average of 5 models |
| rmsd_SR3 | <= 1.4 Å | All-atom RMSD of GLU, average of 5 models |
| rmsd_SR3_max | <= 1.7 Å | Worst GLU RMSD among 5 models. |
| plddt_SR3 | >= 82.0 | All-atom confidence of GLU, average of 5 models |
| rmsd_SR4 | <= 1.5 Å | All-atom RMSD of HIS, average of 5 models |
| rmsd_SR4_max | <= 1.8 Å | Worst HIS RMSD among 5 models. |
| plddt_SR4 | >= 85.0 | All-atom confidence of HIS, average of 5 models |

#### 4.2. Experimental methods

##### Initial screening of estradiol hydroxylation

The synthetic DNA encoding the 96+67 designs was constructed using the IPDassembly approach, with pooled 350nt DNA fragments obtained from Twist Biosciences and spatially separated into corresponding designs using two PCR reactions. The fragments were assembled into full designs and cloned into pDTstrep1 vector (Addgene #262810) encoding a C-terminal StrepTag sequence, and giving the final expressed sequence as `MSG-<design>-GSAWSHPQFEK`. The proteins were transformed into *E. coli* and expressed (with IPTG induction; same protocol as described in Section 3.2.1) as polyclonal cultures. Cells from 4x1 mL expression cultures were harvested by centrifugation, frozen overnight at -20 °C, thawed and lysed using 600 µL of lysis buffer consisting of 50 mM KPi (pH 7.2), 200 mM NaCl, 1 mg/mL lysozyme, 10 µg/mL DNAse I, 0.5 mg/mL Polymyxin B, 1 mM PMSF. Lysis reactions were supplemented with 100 µM hemin (from 2 mM solution in 0.5 M NaOH) and incubated at 37 °C for 1 hour, and proteins were purified by Strep-tag affinity chromatography using 75 µL of Strep-Tactin®XT 4Flow® High Capacity resin. The resin-bound proteins were washed with 2x 0.5 mL KPi buffer, 2x 0.5 mL KPi buffer containing 25 mM imidazole (to wash off any non-specifically bound heme), and again with 2x 0.5 mL KPi buffer. The proteins were eluted off the resin using 300 µL of 50 mM KPi (pH 7.2), 200 mM NaCl buffer containing 50 mM biotin. Enzyme concentrations were not standardized for the initial screen.

Screening reactions contained 200 µL of enzyme eluate, 250 µM estradiol, and 500 µM $H_2O_2$ in 50/200 KPi buffer (pH 7.2) containing 10% (v/v) acetonitrile, in a total volume of 400 µL. Reactions were incubated at room temperature for 16 h and quenched with 400 µL of acetonitrile containing 1,2,3-trimethoxybenzene, giving a final internal-standard concentration of 200 µM in the UPLC sample. Samples were briefly centrifuged, and the supernatants were collected for analysis.

Samples were analyzed without derivatization by UPLC with UV detection at 275 nm using an InfinityLab Poroshell 120 EC-C18 column (2.1 × 100 mm, 1.9 µm) and a 5 µL injection volume. Mobile phases A and B were water and 90% (v/v) acetonitrile in water, respectively, with a flow rate of 0.5 mL $min^{-1}$. The gradient was held at 20% B for 1 min, increased linearly to 95% B over 6 min, held for 1 min, and returned to 20% B over 1 min. Retention times were 2.56 min for 6α-hydroxyestradiol, 2.82 min for estriol, and 3.42 min for the internal standard.

##### Analytical-scale reactions of estradiol hydroxylation

Reactions contained 2.5 µM purified enzyme, 250 µM estradiol, and 500 µM $H_2O_2$ in 50/200 KPi buffer containing 10% (v/v) acetonitrile, in a total volume of 400 µL. Reactions were incubated at room temperature for 16 h and processed as described for the initial screen. Estriol concentrations were determined by UPLC–UV using calibration curves and internal-standard-normalized peak areas. Yield and total turnover number (TTN) were calculated relative to the initial substrate and enzyme concentrations, respectively. Reactions were performed in four independent replicates, and quantitative results are reported as mean ± SD. Formation of 6α-hydroxyestradiol was too low for reliable UV quantification and was confirmed by mass spectrometry.

Product identities were confirmed following dansyl chloride derivatization by comparison of retention times and mass-to-charge ratios with identically derivatized authentic standards using HPLC coupled to a triple-quadrupole mass spectrometer (HPLC–MS/MS), monitoring *m/z* 522.2. Diastereomeric excess was calculated from the integrated areas of resolved diastereomer peaks in the corresponding mass chromatograms. Analysis was performed using reverse-phase LC-MS on Agilent 6475 LC/MS/MS with InfinityLab Poroshell 120 EC-C18, 2.1 x 100 mm, 1.9 μm (A: $H_2O$ with 0.1% Formic Acid, B: acetonitrile with 0.1% Formic Acid). 100 μL of enzymatic reaction was performed with 5 μM of enzyme, 50 μM substrate, and 1 mM $H_2O_2$. After 1 hour of enzymatic reaction, the enzymatic reaction was stopped by the addition of 100 μL of acetonitrile. After centrifugation at 14,000 rpm for 20 min, the supernatant of enzyme reaction solution was filtered with 0.45 μm Nylon membrane filter.

**Activity data for estradiol**

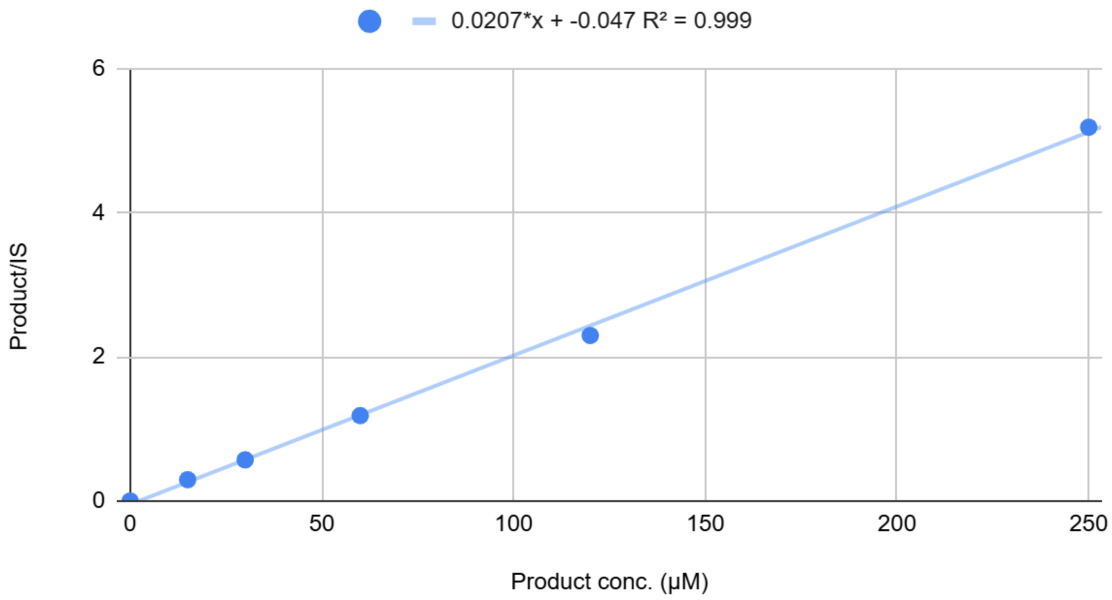


**Figure S41.** Calibration curve of estriol and internal standard (1,2,3-trimethoxybenzene) ratio compared against the concentration of estriol.

**Table S23.** UPLC analysis results of estradiol hydroxylation by **dOHes-2, dOHes-2**-NKVQ.

| **Enzyme** | **Product Area** | **Internal Std Area** | **P/I** | **Concentration µM** | **Yield %** | **TTN** |
|---|---|---|---|---|---|---|
| **dOHes-2** | 1.1 | 27 | 0.04 | 4.2 | 1.7 | 2 |
| **dOHes-2** | 1 | 26 | 0.038 | 4.1 | 1.7 | 2 |
| **dOHes-2** | 1.1 | 27 | 0.040 | 4.2 | 1.7 | 2 |
| **dOHes-2** | 1.1 | 27 | 0.040 | 4.2 | 1.7 | 2 |
| **dOHes-2-NKVQ** | 47 | 27 | 1.74 | 86.3 | 34.6 | 35 |
| **dOHes-2-NKVQ** | 50 | 27 | 1.85 | 91.7 | 36.7 | 37 |
| **dOHes-2-NKVQ** | 52 | 27 | 1.93 | 95.3 | 38.1 | 38 |
| **dOHes-2-NKVQ** | 52 | 26 | 2.00 | 98.9 | 39.6 | 40 |

#### Derivatization of estradiol and related products by dansyl chloride

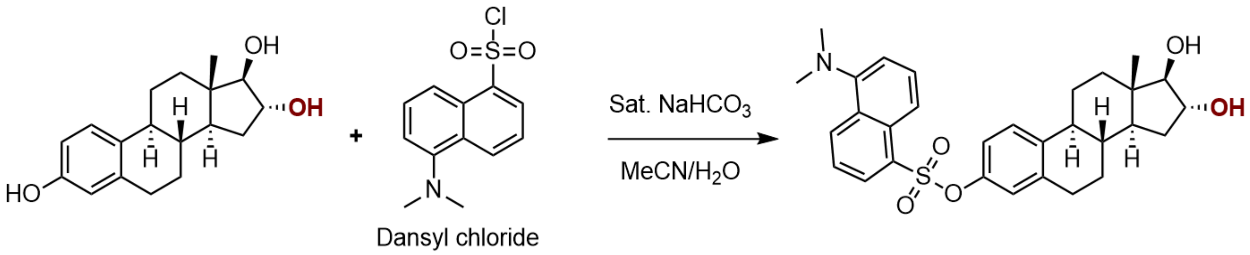


Reaction samples and authentic standards of estradiol, estriol, 16-epiestriol, and 6α- and 6β-hydroxyestradiol were derivatized before LC–MS analysis. Authentic standards were prepared at 1 mM in potassium phosphate buffer (pH 7.2). A 100 µL aliquot of each reaction sample or standard was mixed with freshly prepared dansyl chloride (100 µL, 10 mM in acetonitrile) and saturated aqueous sodium bicarbonate (100 µL). Mixtures were incubated at room temperature for 16 h, diluted with 400 µL water, and centrifuged at 14,000 × *g* for 15 min. Supernatants were analyzed by LC–MS, and products were identified by comparison of retention times and mass-to-charge ratios with authentic standards processed identically.

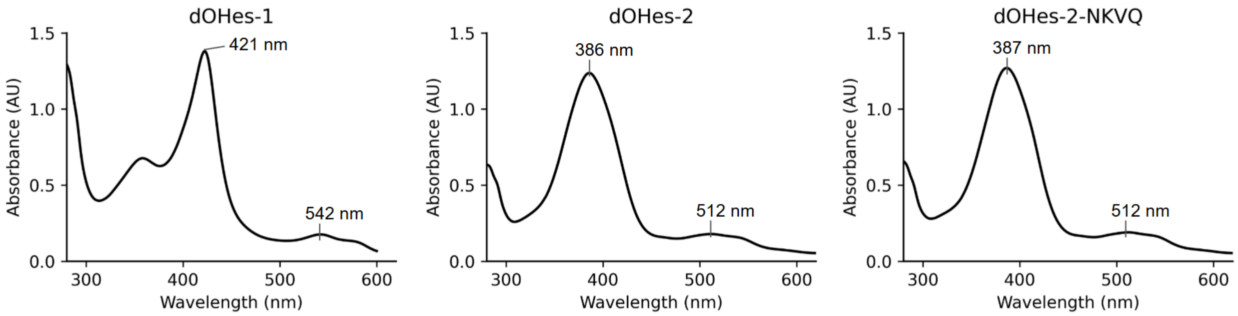


**Figure S42.** UV/Vis spectra of purified enzymes for estradiol hydroxylation.

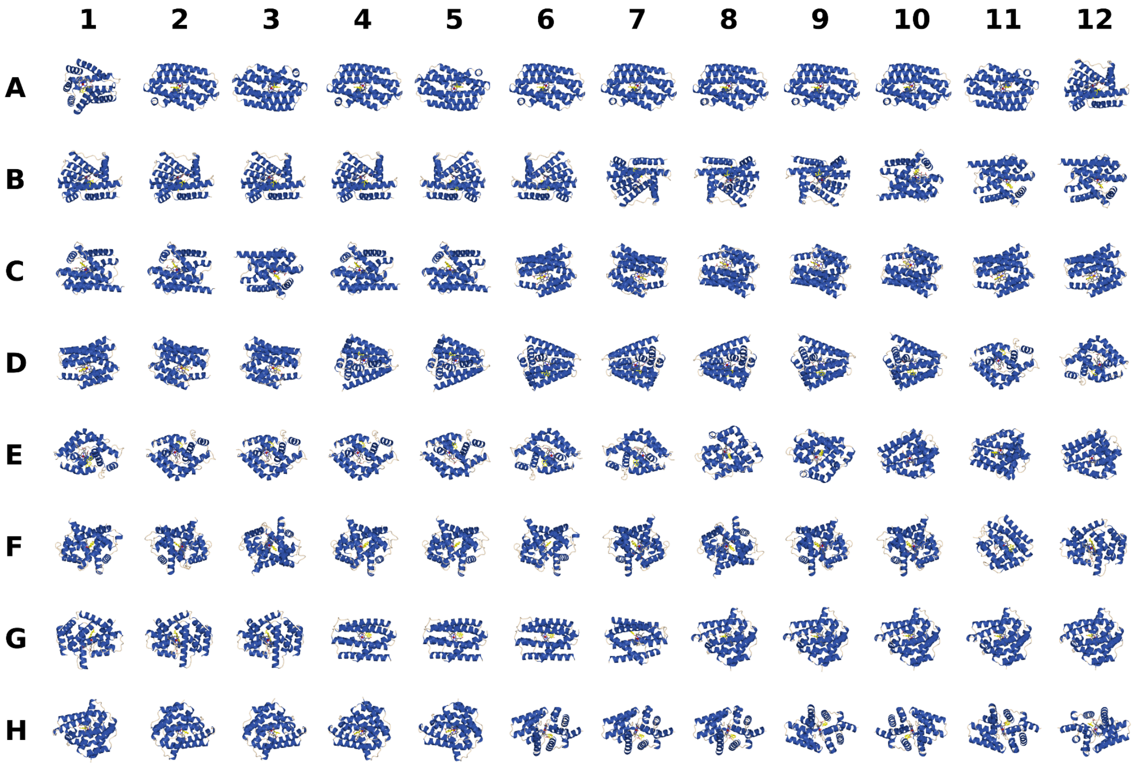


**Figure S43.** Panel of designed enzymes for estradiol C6a hydroxylation.

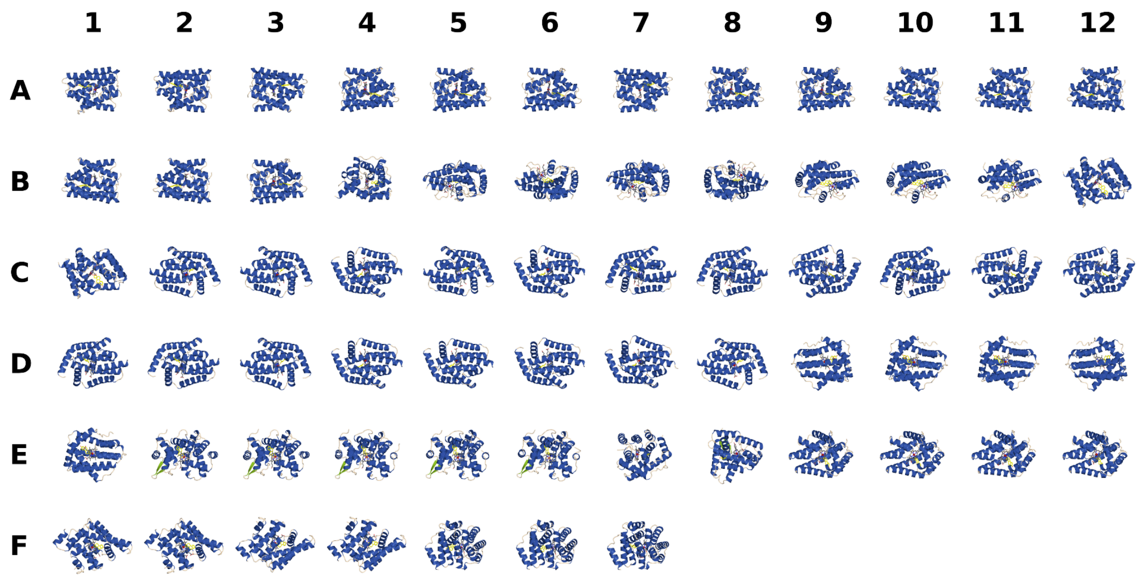


**Figure S44.** Panel of designed enzymes for estradiol C16a hydroxylation.

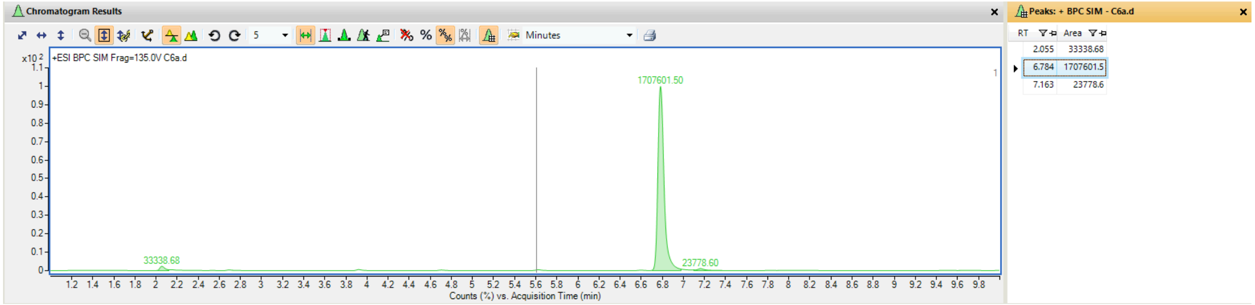


*Commercial standard of C6a*

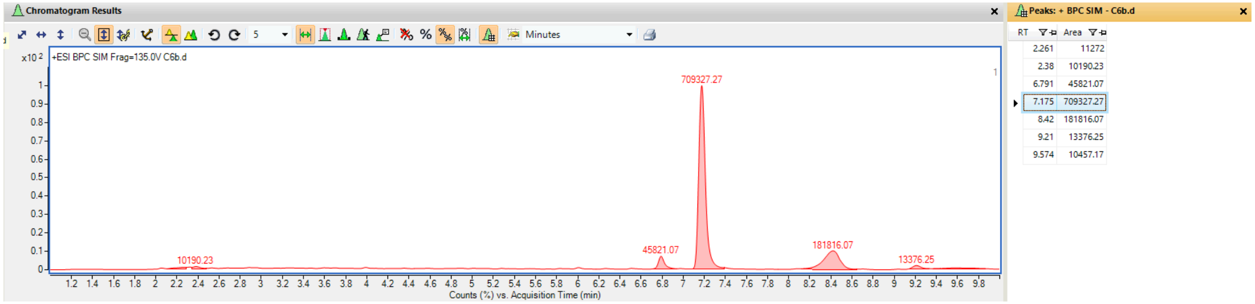


*Commercial standard of C6b*

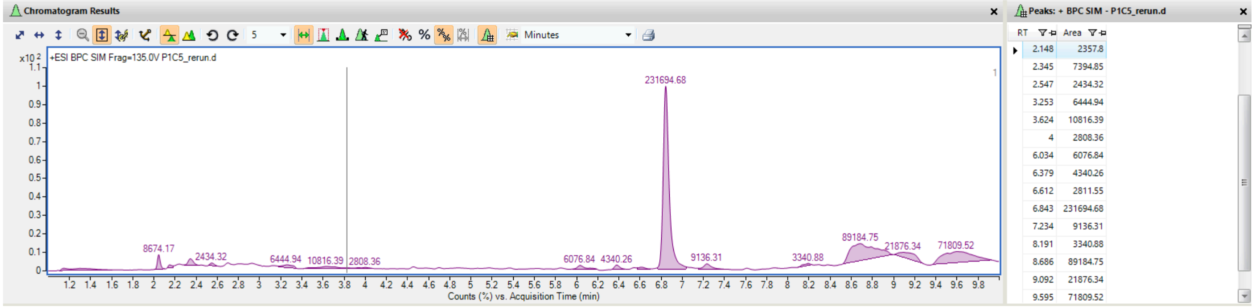


*Enzymatic reaction with dOHes-1*

**Figure S45.** LCMS spectrum for commercial standard of C6 and **dOHes-1** enzymatic reaction.

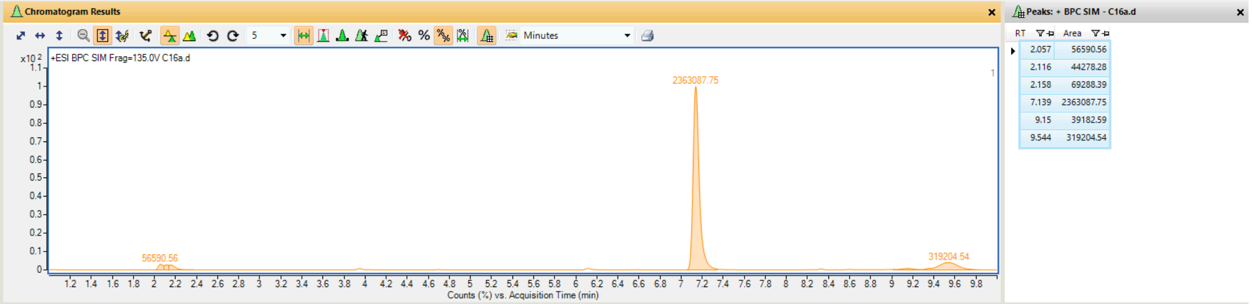


*Commercial standard of C16a*

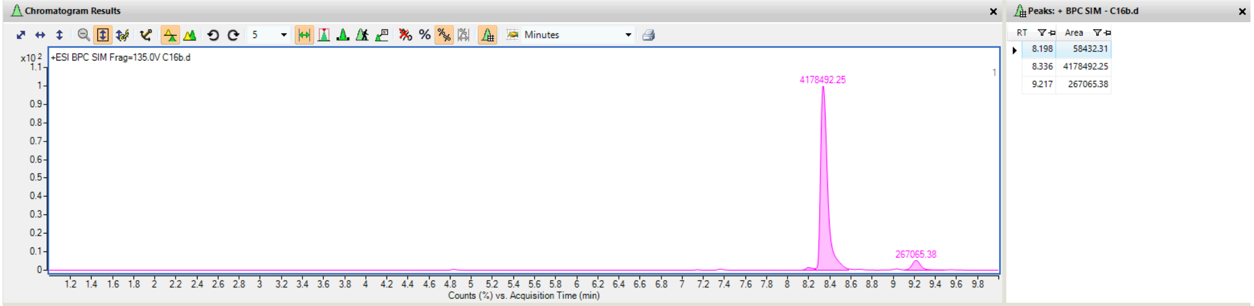


*Commercial standard of C16b*

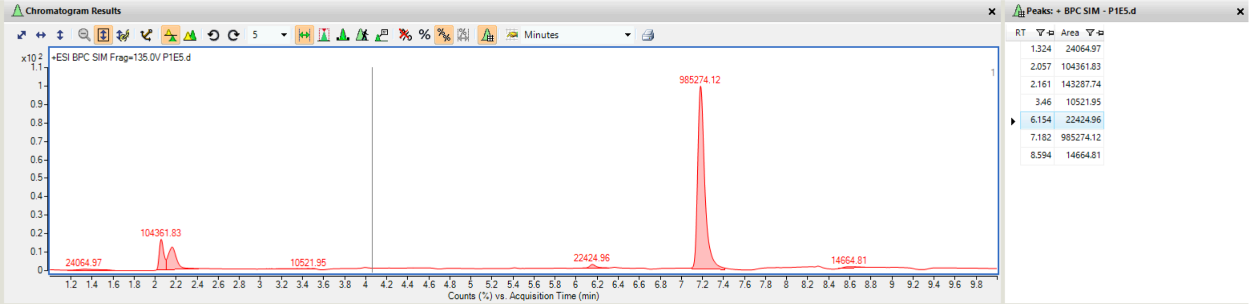


*Enzymatic reaction with dOHes-2*

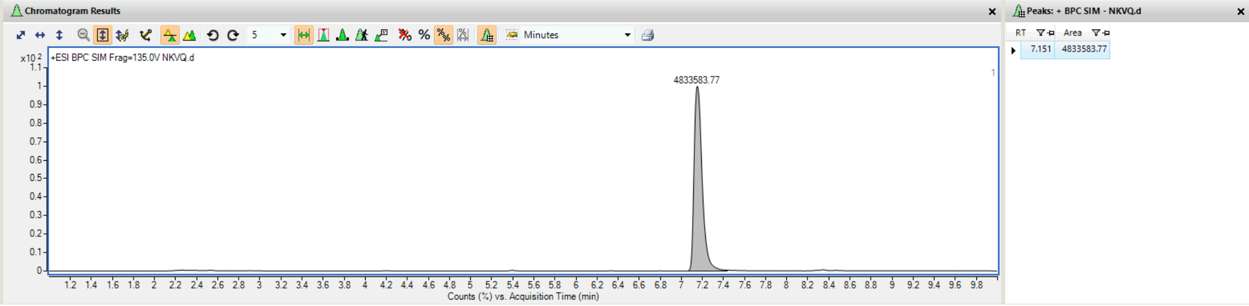


*Enzymatic reaction with dOHes-2-NKVQ*

**Figure S46.** LCMS spectrum for commercial standard of C16 and **dOHes-2/dOHes-2-NKVQ** enzymatic reaction

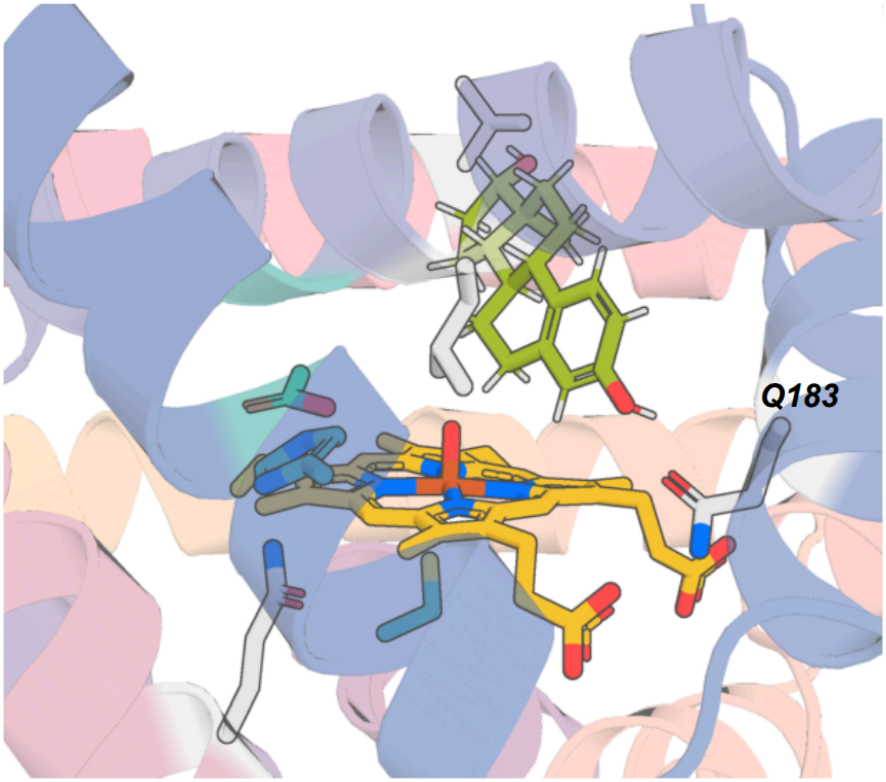


**Figure S47.** Detailed view of **dOHes-1** active site AF3 prediction.

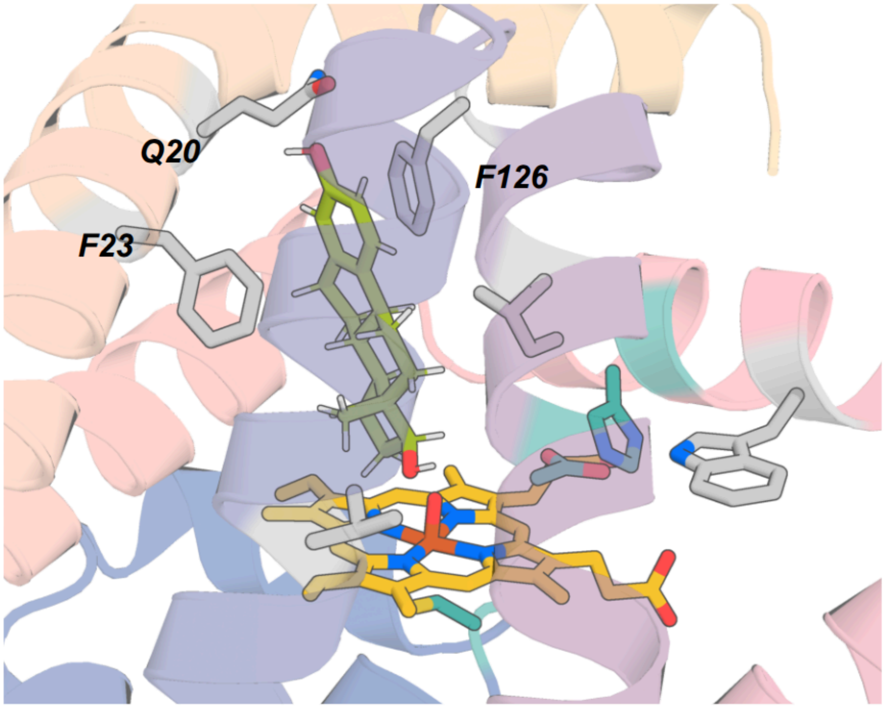


**Figure S48.** Detailed view of **dOHes-2** active site AF3 prediction.

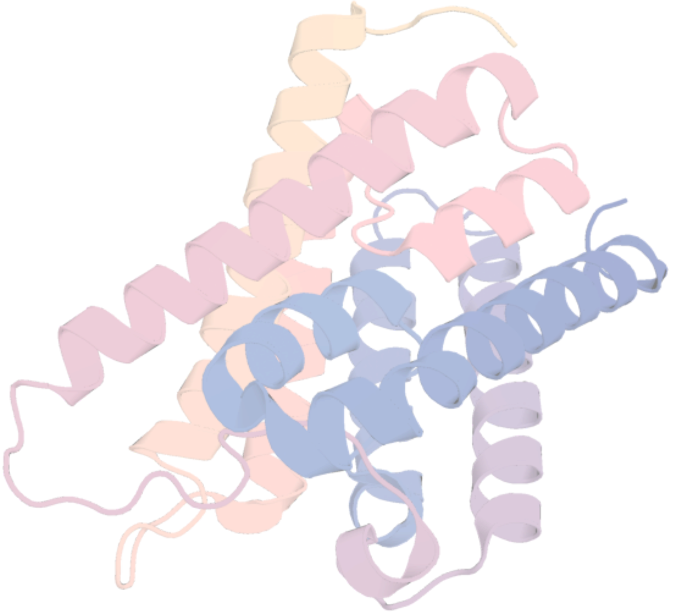

**Figure S49.** Closest structure to **dOHes-1** in PDB, ID: 3LLQ,TM score: 0.45.

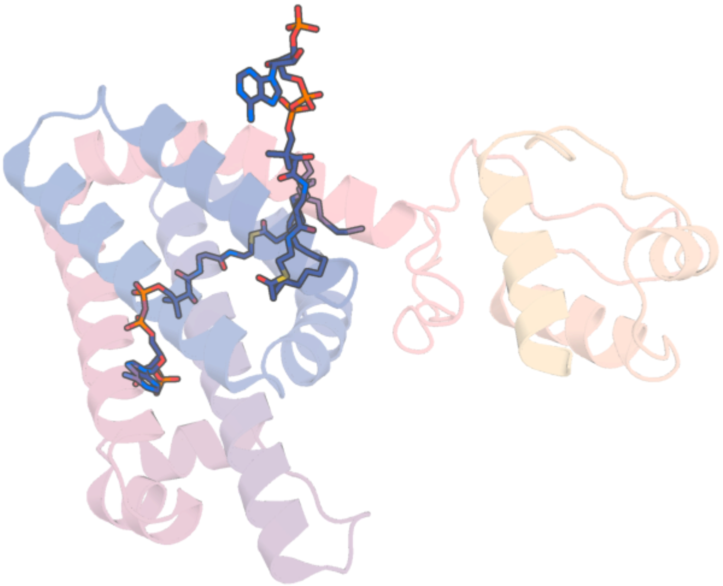

**Figure S50.** Closest structure to **dOHes-2** in PDB, ID: 4PDK,TM score: 0.45.

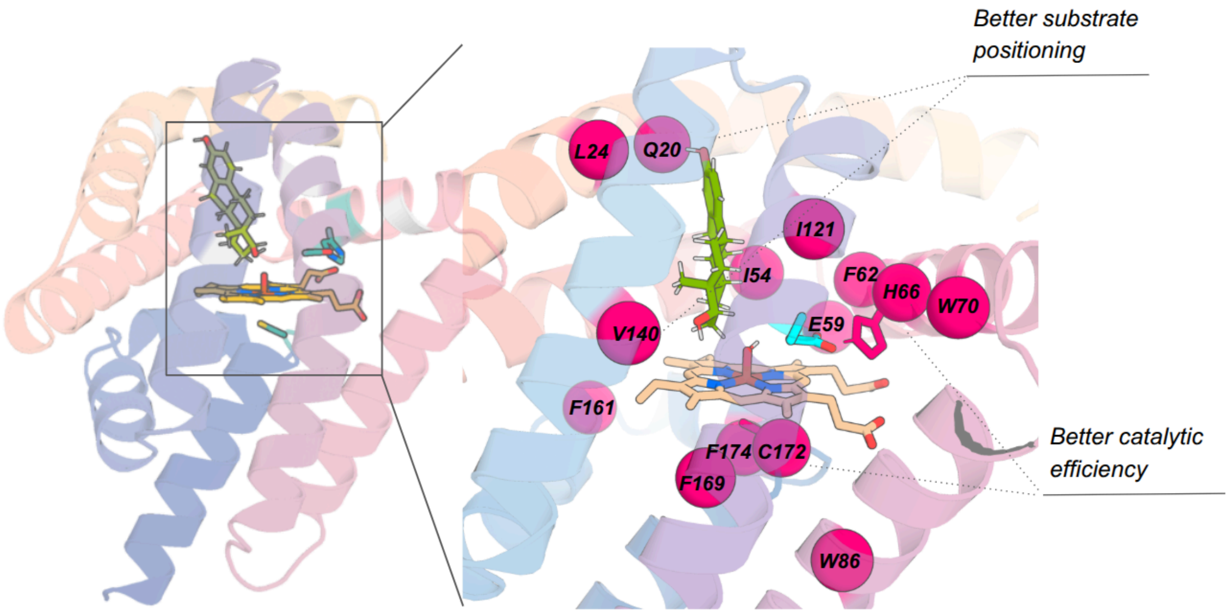


**Figure S51.** Mutagenesis site selection for **dOHes-2**.

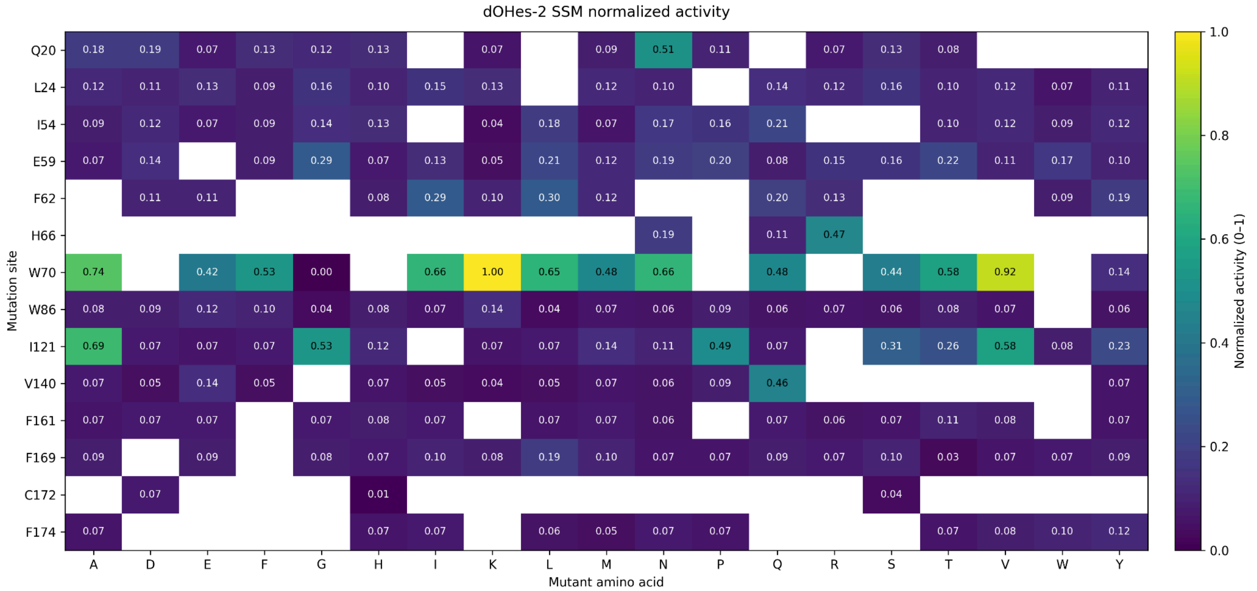


**Figure S52.** AF3 guided site-saturation mutagenesis of **dOHes-2**. Data normalized.

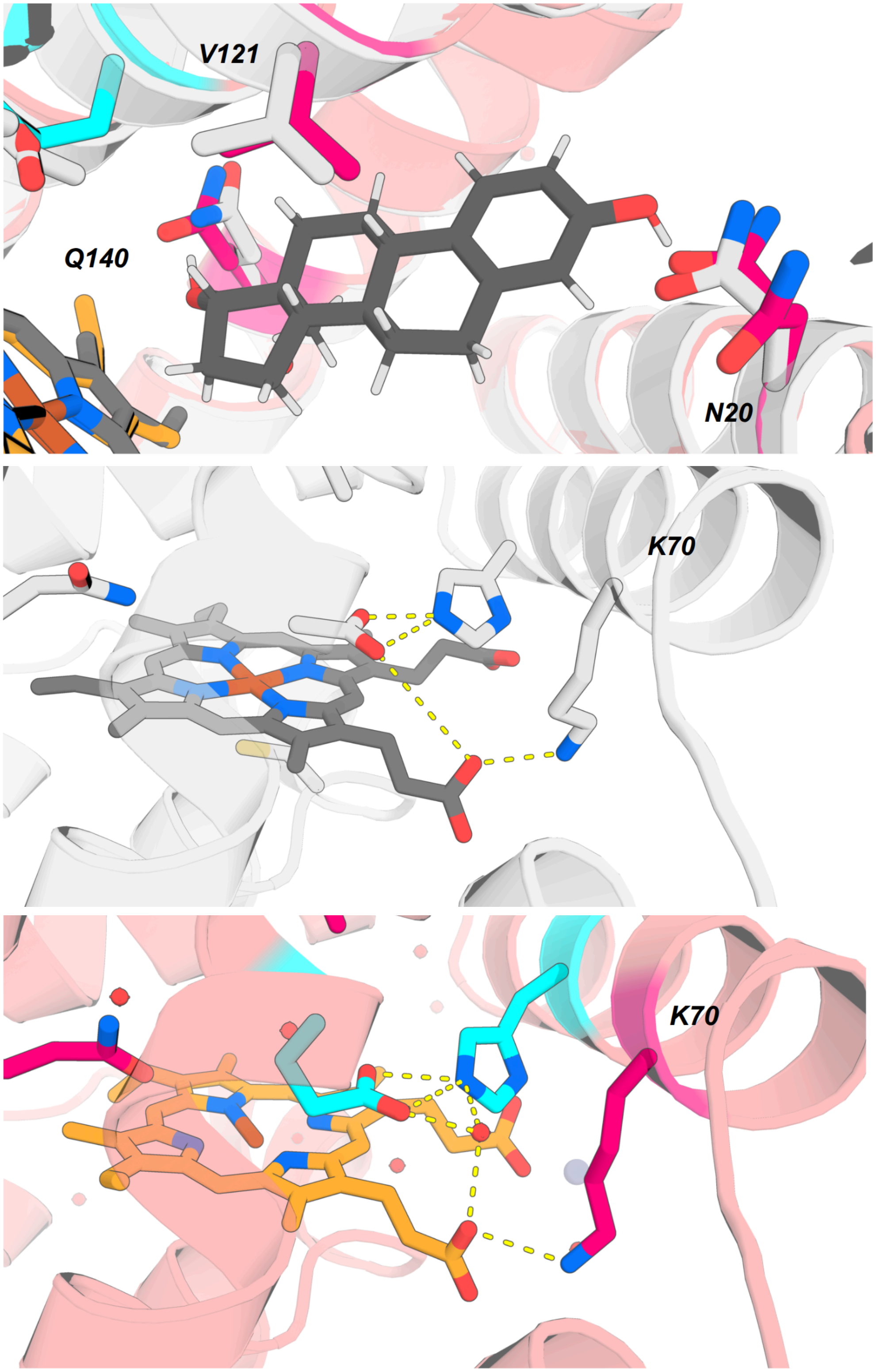


**[ dOHes-2-NKVQ ]**

CRYSTAL // AF3

**Figure S53.** Detailed view of **dOHes-2-NKVQ** active site AF3 model overlay with crystal structure.

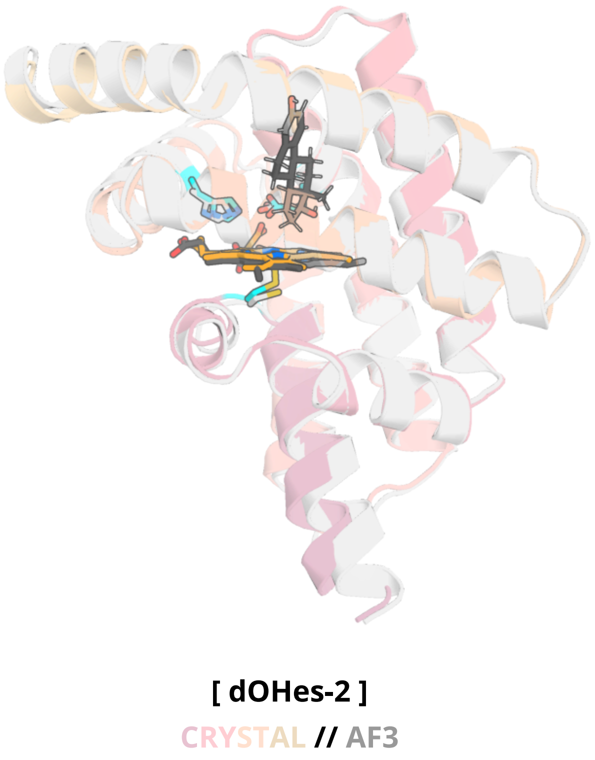


**Figure S54. dOHes-2** AF3 model (gray/black) overlay with crystal structure (colored).

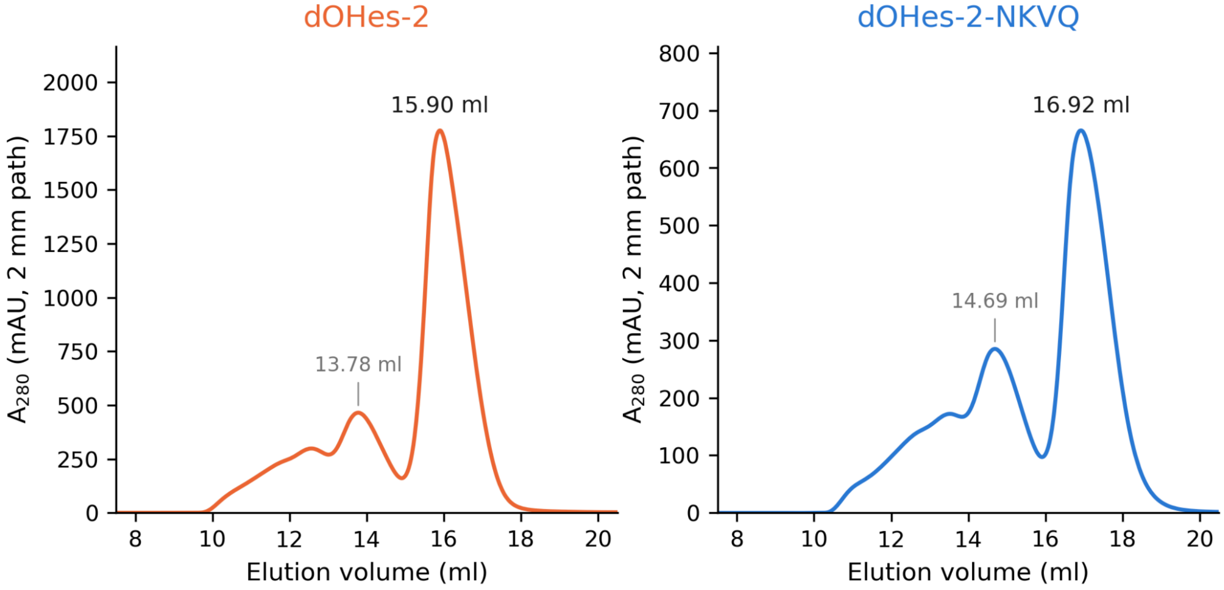


**Figure S55.** Size-exclusion chromatograms of purified **dOHes-2**, **dOHes-2**-NKVQ using Superdex Increase 75 10/300 GL column and 50 mM KPi 200 mM NaCl pH 7.2 running buffer. Monomeric protein used for crystallography is collected from fractions near 15.9 and 16.9 mL peaks, respectively.

### 5. Crystallography

All crystallization experiments were conducted using the sitting drop vapor diffusion method. Crystallization trials were set up in 200 nL drops using the 96-well plate format at 20 °C. Crystallization plates were set up using a Mosquito LCP from SPT Labtech, then imaged using UVEX microscopes and UVEX PS-256 from JAN Scientific.

Diffraction quality crystals formed from protein samples prepared using the following procedures and under the following conditions:

**dOHar-2**

A sample of an uncleaved protein (containing N-terminal His and TEV tags) was purified by size-exclusion chromatography with 50/200 KPi elution. The protein was buffer-exchanged into 25 mM Tris, 100 mM NaCl (pH 8), and concentrated to 19 mg/mL (as judged by A280).

Morpheus Fusion H2: 4 mM Divalent cations mix (0.005 M Mn(II) chloride, 0.005 M Co(II) chloride, 0.005 M Ni(II) chloride, 0.005 M Zn acetate), 0.1 M Buffer System 1 (0.1 M Imidazole; MES (acid)), pH 6.5, 37.5% Precipitant Mix 4 (25% v/v MPD; 25% PEG 1000; 25% w/v PEG 3350). Crystals formed at 2:1 protein/buffer ratio.

**dOHbr-2**

A sample of an uncleaved protein (containing C-terminal SNAC and His tags) was purified by size-exclusion chromatography with 25 mM Tris, 100 mM NaCl (pH 8) elution. The protein was concentrated to 25 mg/mL (as judged by A280).

Index H7 and optimized with GRAS D3: 0.15 M DL-Malic acid, pH 7.0, 20% (w/v) Polyethylene glycol 3,350 + 1.1 mM Pyruvic acid. Crystals formed at 1:1 protein/buffer ratio.

Additional crystals formed under the following conditions. BCS C9: 0.15 M Magnesium acetate tetrahydrate, 0.1 M Sodium citrate, pH 5.6 , 20% v/v PEG Smear Broad; 1:1 protein/buffer. Morpheus H1: 0.1 M amino acids mix (DL-Glutamic acid; DL-Alanine; Glycine; DL-Lysine; DL-Serine), 0.1 M Buffer System 1 (Imidazole; MES), pH 6.5, 30% v/v Precipitant Mix 1 (40% PEG 500 MME; 20% PEG 20000); 1:2 protein/buffer.

**dOHbr-2**-L125M-L150F

A sample of an uncleaved protein (containing C-terminal SNAC and His tags) was purified by size-exclusion chromatography with 25 mM Tris, 100 mM NaCl (pH 8) elution. The protein was concentrated to 25 mg/mL (as judged by A280).

Morpheus H1: 0.1 M Amino acids Mix (0.1 M DL-Glutamic acid; DL-Alanine; Glycine; DL-Lysine; DL-Serine), 0.1 M Buffer System 1 (0.1 M Imidazole;MES), pH 6.5, 30% (v/v) Precipitant Mix 1 (40% PEG 500 MME; 20% PEG 20,000). Crystals formed at 1:1 protein/buffer ratio.

**dOHbr-5**

A sample of an uncleaved protein (containing N-terminal His and TEV tags) was purified by size-exclusion chromatography with 25 mM Tris, 100 mM NaCl (pH 8) elution. The protein was concentrated to 20 mg/mL (as judged by A280).

Index A2 and optimized with GRAS C11: 0.1 M Sodium acetate, pH 4.5, 2.0 M Ammonium sulfate, 4 mM α-Ketoglutaric acid disodium salt. Crystals formed at 1:1 protein/buffer ratio.

**dOHes-2**

A sample of an uncleaved protein (containing C-terminal Streptag) was purified by size-exclusion chromatography with 25 mM Tris, 100 mM NaCl (pH 8) elution. The protein was concentrated to 20 mg/mL (as judged by A280).

Classics1 G11: 0.2 M Ammonium sulfate, 0.1 M Sodium acetate, pH 4.6, 30% (w/v) PEG 2000 MME. Crystals formed at 1:1 protein/buffer ratio.

**dOHes-2-NKVQ**

A sample of an uncleaved protein (containing C-terminal Streptag) was purified by size-exclusion chromatography with 25 mM Tris, 100 mM NaCl (pH 8) elution. The protein was concentrated to 20 mg/mL (as judged by A280).

Classics1 G7: 0.01 M Zinc sulfate, 0.1 M MES, pH 6.5, 25% (w/v) PEG 550 MME. Crystals formed at 1:1 protein/buffer ratio.

Diffraction data were collected at the Advanced Photon Source beamline on 24-ID-E and National Synchrotron Light Source II on FMX/AMX. Diffraction data were processed using xia2/DIALS (*17*, *18*) and anisotropy-correction was performed using the STARANISO server (*19*). Structure determination and refinement starting phases were obtained by molecular replacement using DIMPLE (*20*) and their respective designed model. Iterative model building and refinement were performed manually using COOT (*21*) following molecular replacement and subsequent structure refinement was performed with CCP4 Cloud Refmacat (*22*, *23*). Initial refinement was conducted using BUSTER (*24*) to rapidly fix Ramachandran, rotamer, and density fit outliers, refining to convergence. PDB-REDO (*25*) was used to assess the model quality in between refinements and to fix any rotamer and density fit outliers. Final refinement was carried out in Phenix (*26*), with manual model building performed in COOT. The model quality was evaluated using MolProbity (*27*). Data collection and refinement statistics are recorded in Tables S24-25. Structural figures were prepared using PyMOL.

**Table S24.** Crystallographic data of designs **dOHar-2**, **dOHbr-2**, **dOHbr-2**-L125M-L150F, **dOHbr-5**.

| | **dOHar-2** | **dOHbr-2** | **dOHbr-2**-L125M-L150F | **dOHbr-5** |
|---|---|---|---|---|
| **Data collection** | | | | |
| Space group | $P\ 2_1\ 2_1\ 2_1$ | $C\ 1\ 2\ 1$ | $P\ 1$ | $P\ 4_1\ 2_1\ 2$ |
| Cell dimensions | | | | |
| *a, b, c* (Å) | 47.32, 51.32, 71.71 | 159.91, 59.42, 45.50 | 45.61, 60.48, 84.34 | 55.20, 55.20, 138.14 |
| α, β, γ (°) | 90, 90, 90 | 90, 106.12, 90 | 107.23, 105.49, 90.41 | 90, 90, 90 |
| Resolution (Å) | 41.74 - 1.86 (2.04 - 1.86) | 55.66 - 1.94 (2.12 - 1.94) | 40.41 - 2.75 (3.17 - 2.75) | 51.27 - 1.77 (1.99 - 1.77) |
| $R_{merge}$ | 0.102 (1.644) | 0.0950 (0.597) | 0.147 (0.912) | 0.142 (1.552) |
| $I / \sigma I$ | 8.4 (1.5) | 10.7 (1.6) | 10.1 (1.4) | 14.6 (1.5) |
| Spherical Completeness (%) | 86.7 (46.5) | 65.4 (10.3) | 65.8 (17.8) | 78.0 (24.8) |
| Ellipsoidal Completeness (%) | 93.6 (68.0) | 89.1 (46.7) | 89.7 (62.1) | 93.9 (60.2) |
| Redundancy | 6.6 (6.9) | 3.6 (3.9) | 3.9 (3.8) | 26.1 (24.9) |
| $CC_{1/2}$ | 0.998 (0.678) | 0.997 (0.789) | 0.993 (0.599) | 0.999 (0.878) |
| **Refinement** | | | | |
| Resolution (Å) | 41.74 - 1.86 (1.94 - 1.86) | 43.71 - 1.94 (2.03 - 1.94) | 38.64 - 2.75 (2.97 - 2.75) | 51.26 - 1.77 (1.88 - 1.77) |
| No. reflections | 13,184 (375) | 22,472 (273) | 14,032 (344) | 16,941 (371) |
| $R_{work} / R_{free}$ | 0.1990 (0.3778) /0.2462 (0.4260) | 0.2005 (0.2414) /0.2327 (0.3115) | 0.2045 (0.3124) /0.2612 (0.3002) | 0.1834 (0.3434) /0.2255 (0.4712) |
| No. atoms | | | | |
| Protein | 1417 | 3059 | 5686 | 1555 |
| Water | 42 | 95 | 36 | 123 |
| Ligand | 49 | 104 | 185 | 56 |
| B-factors | | | | |
| Protein | 47 | 34 | 58 | 39 |
| Water | 51 | 35 | 39 | 43 |
| Ligand | 31 | 25 | 43 | 34 |
| R.m.s. deviations | | | | |
| Bond lengths (Å) | 0.012 | 0.018 | 0.006 | 0.005 |
| Bond angles (°) | 1.110 | 1.590 | 0.920 | 0.830 |

*Single Crystal used for each data collection. *Values in parentheses are for highest-resolution shell.

**Table S25.** Crystallographic data of designs **dOHes-2** and **dOHes-2-NKVQ**.

| | **dOHes-2** | **dOHes-2-NKVQ** |
|---|---|---|
| **Data collection** | | |
| Space group | *C 1 2 1* | $P\ 2_1\ 2_1\ 2_1$ |
| Cell dimensions | | |
| *a*, *b*, *c* (Å) | 133.35, 57.32, 43.80 | 44.29, 52.95, 64.54 |
| α, β, γ (°) | 90, 106.21, 90 | 90, 90, 90 |
| Resolution (Å) | 42.06 - 3.66 (4.19 - 3.66) | 36.48 - 2.00 (2.21 - 2.00) |
| $R_{merge}$ | 0.143 (0.551) | 0.113 (1.294) |
| $I$ / σ$I$ | 3.7 (1.5) | 10.6 (2.1) |
| Spherical Completeness (%) | 62.6 (17.5) | 93.1 (73.4) |
| Ellipsoidal Completeness (%) | 83.0 (46.7) | 96.9 (86.6) |
| $CC_{1/2}$ | 0.979 (0.833) | 0.997 (0.827) |
| **Refinement** | | |
| Resolution (Å) | 42.06 - 3.66 (4.61 - 3.66) | 36.48 - 2.00 (2.20 - 2.00) |
| No. reflections | 2,248 (550) | 9,983 (1,900) |
| $R_{work}$ / $R_{free}$ | 0.3295 (0.3525) /0.3581 (0.3672) | 0.2065 (0.2571) /0.2416 (0.2894) |
| No. atoms | | |
| Protein | 3078 | 1551 |
| Water | n/a | 21 |
| Ligand | 86 | 53 |
| *B*-factors | | |
| Protein | 203 | 51 |
| Water | n/a | 52 |
| Ligand | 138 | 45 |
| R.m.s. deviations | | |
| Bond lengths (Å) | 0.006 | 0.008 |
| Bond angles (°) | 0.900 | 0.920 |

*Single Crystal used for each data collection. *Values in parentheses are for highest-resolution shell.

5. Matching and enzyme design geometric constraint file format. https://docs.rosettacommons.org/docs/latest/rosetta_basics/file_types/match-cstfile-format.

6. I. Kalvet, *Invrotzyme: Utility for Building Inverse Rotamer Assemblies out of a Rosetta Matcher/enzdes Constraint File* (Github; https://github.com/ikalvet/invrotzyme).

7. B. L. T. Edward A. Berry, Simultaneous determination of hemes a, b, and c from pyridine hemochrome spectra. *Analytical Biochemistry* **161**, 1–15 (1987).

8. Z. Gan, J. Feng, J. Yin, J. Huang, B. Wang, J. Z. H. Zhang, Diverse mechanisms for the aromatic hydroxylation: Insights into the mechanisms of the coumarin hydroxylation by CYP2A6. *ACS Catal.* **14**, 16277–16286 (2024).

9. J. Qian, L. F. Milles, B. I. M. Wicky, R. J. Ragotte, A. Motmaen, A. J. Borst, R. Skotheim, S. Ols, B. Coventry, X. Li, R. D. Kibler, I. Goreshnik, M. Expòsit, K. Loré, L. Stewart, D. Baker, Accelerating protein design by scaling experimental characterization. *Nat. Commun.* **17**, 9945 (2026).

10. C. Xu, Y. Zhang, H. Sun, J. Ai, M. Ren, Development of a two-photon fluorescent probe for imaging hydrogen sulfide (H2S) in living cells and zebrafish. *Anal. Methods* **15**, 1948–1952 (2023).

11. R.-J. He, Z.-H. Tian, J. Huang, M.-R. Sun, F. Wei, C.-Y. Li, H.-R. Zeng, F. Zhang, X.-Q. Guan, Y. Feng, X.-M. Meng, H. Yang, G.-B. Ge, Rationally engineered CYP3A4 fluorogenic

substrates for functional imaging analysis and drug-drug interaction studies. *J. Med. Chem.* **66**, 6743–6755 (2023).

12. I. Kalvet, *Fastmpnndesign: A Framework for Integrated LigandMPNN and Rosetta Repack/minimize/relax Sequence Design Applications, Mimicking Rosetta FastDesign* (Github; https://github.com/ikalvet/fastmpnndesign).
